# Shine: A novel strategy to extract specific, sensitive and well-conserved biomarkers from massive microbial genomic datasets

**DOI:** 10.1101/2021.11.11.468318

**Authors:** Cong Ji, Junbin (Jack) Shao

## Abstract

**Background:** Concentrations of the pathogenic microorganisms’ DNA in biological samples are typically low. Therefore, DNA diagnostics of common infections are costly, rarely accurate, and challenging. Limited by failing to cover updated epidemic testing samples, computational services are difficult to implement in clinical applications without complex customized settings. Furthermore, the combined biomarkers used to maintain high conservation may not be cost effective and could cause several experimental errors in many clinical settings. Given the limitations of recent developed technology, 16S rRNA is too conserved to distinguish closely related species, and mosaic plasmids are not effective as well because of their uneven distribution across prokaryotic taxa.

**Results:** Here, we provide a computational strategy, Shine, that allows extraction of specific, sensitive and well-conserved biomarkers from massive microbial genomic datasets. Distinguished with simple concatenations with blast-based filtering, our method involves a *de novo* genome alignment-based pipeline to explore the original and specific repetitive biomarkers in the defined population. It can cover all members to detect newly discovered multicopy conserved species-specific or even subspecies-specific target probes and primer sets. The method has been successfully applied to a number of clinical projects and has the overwhelming advantages of automated detection of all pathogenic microorganisms without the limitations of genome annotation and incompletely assembled motifs. Using on our pipeline, users may select different configuration parameters depending on the purpose of the project for routine clinical detection practices on the website https://bioinfo.liferiver.com.cn with easy registration.

**Conclusions:** The proposed strategy is suitable for identifying shared phylogenetic markers while featuring low rates of false positive or false negative. This technology is suitable for the automatic design of minimal and efficient PCR primers and other types of detection probes.

## Introduction

Rapid detection of pathogenic organisms is a crucial undertaking related to health, safety and wellbeing, especially for the early detection of pathogens for diagnosing and preventing diseases[1–3]. While the field of diagnostics is rapidly evolving, polymerase chain reaction (PCR) remains the gold standard for nucleic acid-based diagnostic assays, in part due to its reliability, flexibility and wide deployment[4]. The development of an emergency-use molecular laboratory-developed test (LDT) will be useful to laboratories for future outbreaks and will help lower barriers to establishing fast and accurate diagnostic testing in crisis conditions[4]. Nevertheless, the deoxyribonucleic acid (DNA) concentrations from pathogenic microorganisms in biological samples are mostly very low and close to the detection limit, so pathogen detection has become one of the most challenging task in clinical applications[5]. For instance, traditional PCR and real-time PCR often lack detection sensitivity[6, 7], and other methods, such as two-step nested PCR, may have better sensitivity, but they are not feasible for routine tests and present a high risk of contamination[8]. Consequently, these methods are relatively time consuming and costly and are not very accuracy. Thus, it is necessary to explore biomarkers with high performance to improve the quality of reagents.

Accuracy and sensitivity are important in the quality of nucleic acid detection reagents, with accuracy attributed mainly to high specificity and conservation. Due to the lack of shared universal phylogenetic biomarkers, small changes in the concentrations of single biomarkers are not sufficient for the accurate prediction of viral/bacterial community-acquired pneumonia[9]. Although automated identification of species-specific repetitive DNA sequences and their utilization for detecting microbial organisms by MultiMPrimer3 have been widely reported[10], this tool is limited by the lack of customized settings, especially for clinical applications. For instance, when unknown microorganisms cause epidemic outbreaks[11], the pathogenic microorganism database may lead to the original probe primer design failing to cover the epidemic pathogenic microorganisms. To improve the predictive power of detection, biomarker combinations have become the primary choice in many studies[12–14]. However, this approach may not be cost effective and can lead to several experimental errors in clinical settings. Therefore, the importance of exploring minimal shared universal phylogenetic and repetitive biomarkers with primers and probes to improve the detection sensitivity and accuracy for any pathogen cannot be overestimated.

Traditional method is to select 16S ribosomal RNA (16S rRNA) sequences[15] and specific plasmid[16] from known literatures as the PCR designing templates for pathogenic microorganisms. Thereof, 16S rRNA gene sequence analysis can be routinely used for the identification of mycobacteria and facilitate the exploration of novel pathogens and non-cultured bacteria[17–20]. However, few studies have reported consensus definitions of genera or species based on 16S rRNA gene sequence data. Several studies have highlighted that rRNA and other marker genes cannot be directly used to fully predict the functional potential of the bacterial community[21]. The main reason for this is that not all rRNA genes are species-specific, because the sequences of them are too conserved to be distinguishable, especially between closely related species. The mechanisms and selective pressures causing the formation of mosaic plasmids do not occur evenly across all species, and plasmids may provide different levels of potential variation to different species that are abundant and unevenly distributed across prokaryotic taxa[22]. Finally, many clinical laboratories have to validate the quality of assays by other primers and probes since plasmid PCR testing has obviously high risks of generating false positive or false negative results.

Here, we propose a new strategy, Shine, which is essential for performing comparative genomics in routine tests and rapid detection of pathogenic organisms for improved performance. It learns from previous tools which have involved reconstructing the evolutionary histories of species and inferring the selective forces shaping the evolution of genes and species by comparative analysis of molecular sequence data, such as those using Molecular Evolutionary Genetics Analysis (MEGA5)[23] or Phylogenetic Analysis by Maximum Likelihood (PAML4)[24]. We hypothesized that the more comprehensive the genomic data are, the more effective detection biomarkers. To explore specific, sensitive and conserved biomarkers from massive microbial genomic data within populations, we aimed to develop a computational strategy to improve the quality of nucleic acid detection reagents, which has been validated in several clinical projects.

## Results

### Illustration of our strategy

We developed a general de novo genome alignment-based pipeline to explore the original and specific multicopy biomarkers in the defined populations to cover all the members. Either repetitive regions or specific regions were preferred, and raw data were divided by two selection methods, i.e., prioritizing multicopy regions for large genomes and prioritizing specific regions for small genomes. Then, the data were processed in the other modules separately. That is, by the first method, the data are processed into the next module to search for specific regions. If the second method is used, the data are pushed into the next module to search for multicopy regions. Finally, we focused on searching for consensus sequences and designing the best primer and probe sets. Correspondingly, it was necessary to perform a double-check validation in every module, as shown in Figure 1.

**Figure 1.**
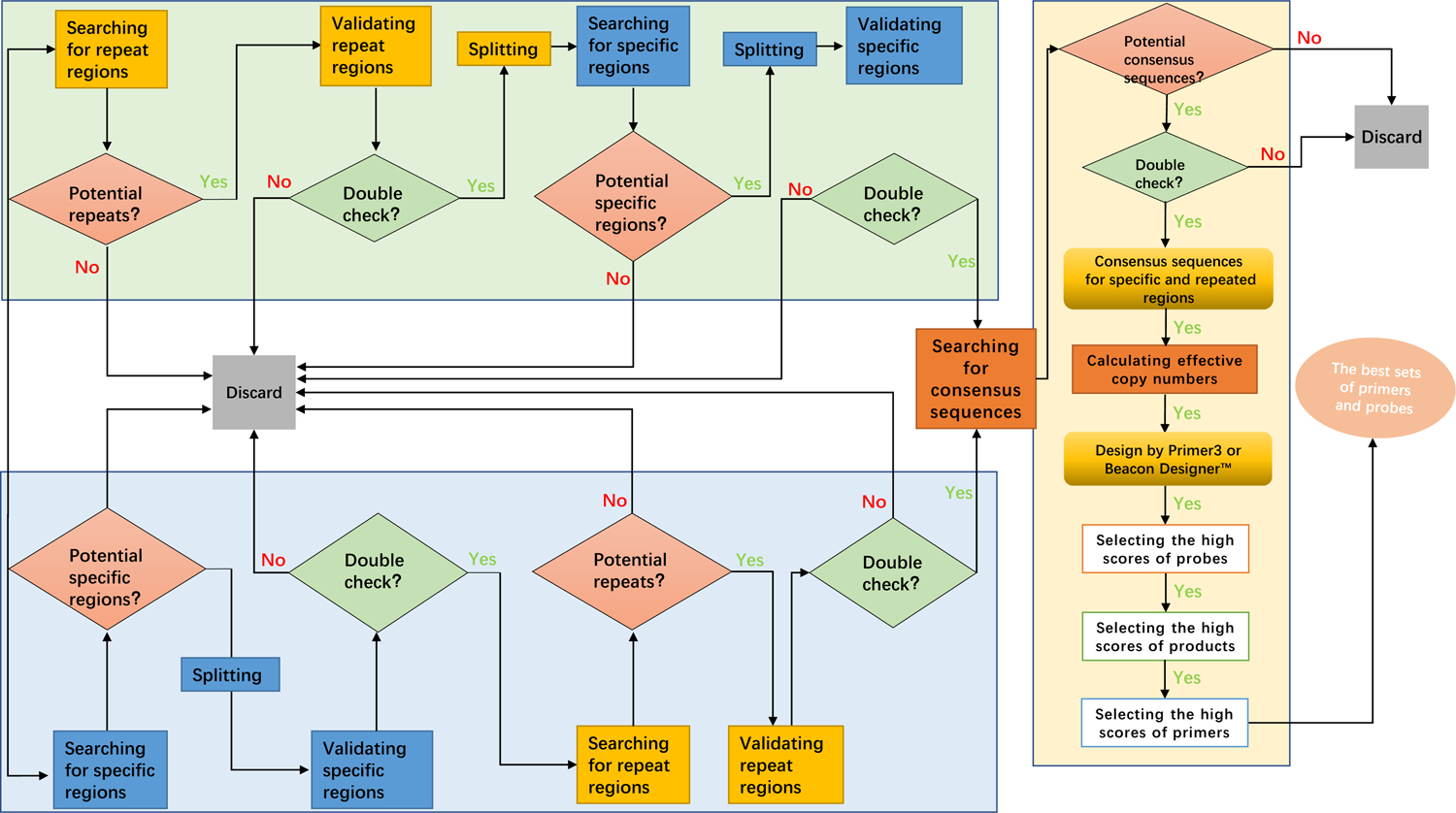
Schematic map of Shine. This new strategy was used to explore specific, sensitive and conserved biomarkers to cover all members of defined populations. The total pipeline was divided into two directions, i.e., to search specific regions preferentially or to search sensitive regions preferentially. At last, searching for consensus sequences for specific and repeated regions was to be available for the best sets of primers and probes.

One of the important details was common block deletions used to search the specific regions, and each genome of the target strains was compared with every genome of the control strains for N calculations, distinguished with a blast-based filtering method. Common block deletions lasted for X generations with multiple threads to search specific regions or subspecific regions for M target strains, as illustrated in Figure 2. To accelerate the comparison, in a preferred embodiment, the first-round divided fragments T1-Tn were each compared with the whole-genome sequences of the remaining comparison strains by group iterations, as shown in Figure 2: 1) upon dividing the target comparison strains into M groups, each group included a plurality of the same comparison strains; 2) simultaneously comparing the first-round divided fragment T^1^ with the whole-genome sequences of comparison strain 1 in the first group one to one and removing fragments for which the similarity exceeded the preset value, the plurality of residual fragments was obtained as the first-round candidate sequence library of the first-round divided fragment T^1^; 3) simultaneously comparing the previous-round candidate sequence library of the first-round divided fragment T^1^ with whole-genome sequences of comparison strain 2 in the next group one to one and removing fragments for which the similarity exceeded the preset value, the plurality of residual fragments was obtained as the next-round candidate sequence library of the first-round divided fragment T^2^; 4) operations from the second-round candidate sequence library were repeated until the N-round candidate sequence library was obtained as the candidate specific sequence library of the N-round divided fragments Tn; 5) the collection of all the candidate specific sequence libraries of the N-round divided fragments were the candidate specific regions. Further validation comprised two parts of scoring and filtering the potential specific regions: blast against all organisms and comparison of genomic sequences with those of control strains so that we could identify the specific regions with stringent quality standards.

**Figure 2.**
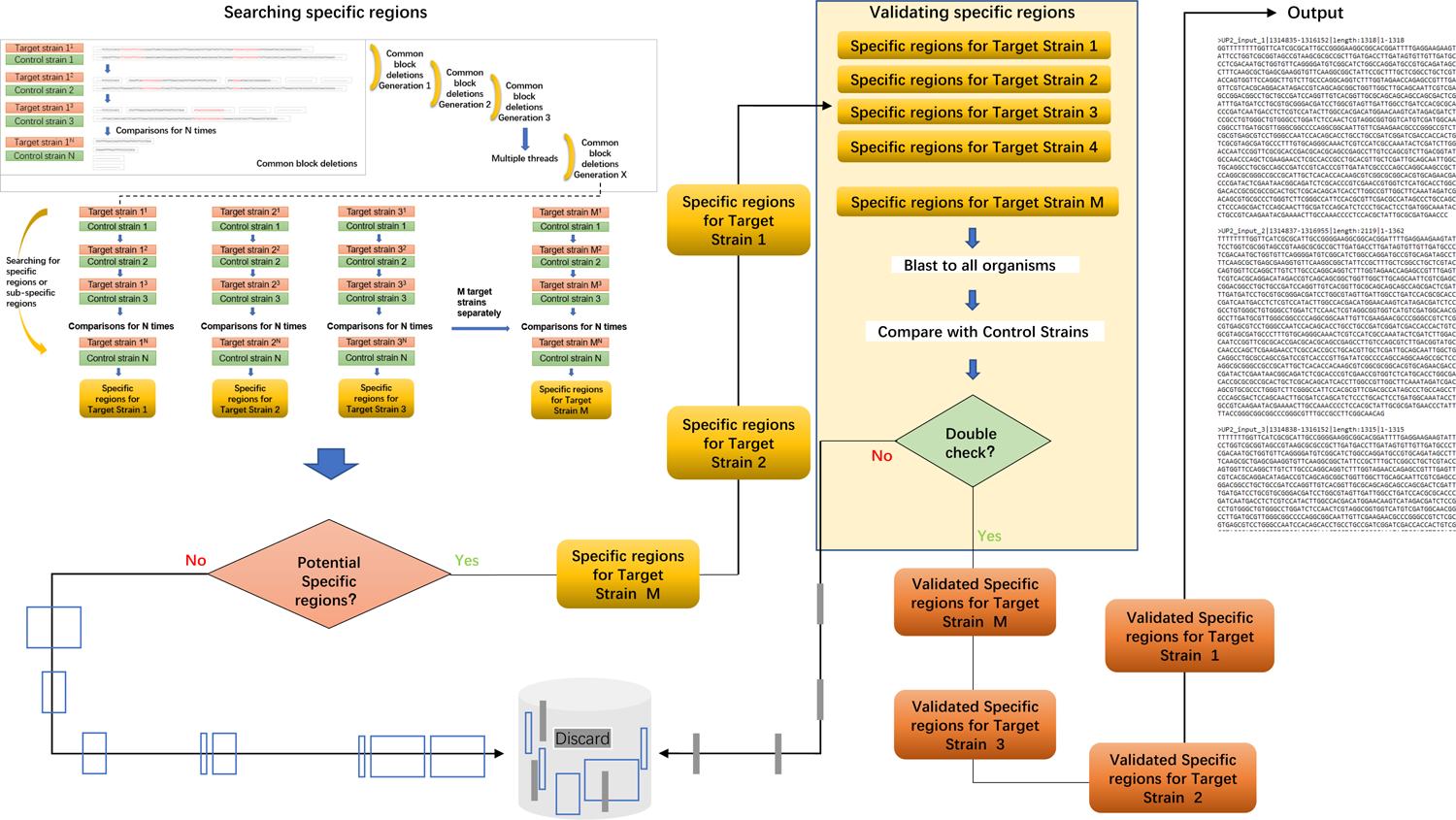
Illustration of the submodule preferentially searching for the specific regions. 1) the microbial target fragments were compared with the whole-genome sequences of one or more comparison strains one to one to obtain several residual fragments as the first-round cut fragments T1-Tn; 2) then compared with whole-genome sequences of the remaining comparison strains, to obtain the collection of residual cut fragments as the candidate specific regions of the microbial target fragments; and 3) the specific regions were then verified and obtained to determine.

The other key aspect was the search for repetitive regions with different copy numbers in every target strain and extracting potential repeats for validation by remapping and statistically summarizing the mean copy numbers and variations for each repeat, while the rest were discarded, as shown in Figure 3. Here, each repeat was filtered into the classified summation so that we could obtain the minimal accurate sets of repetitive sequences from the whole genome. When identifying multicopy regions in microbial target fragments, the motifs are connected together before searching for candidate multicopy regions, in which the microbial target fragments often have multiple incomplete motifs.

**Figure 3.**
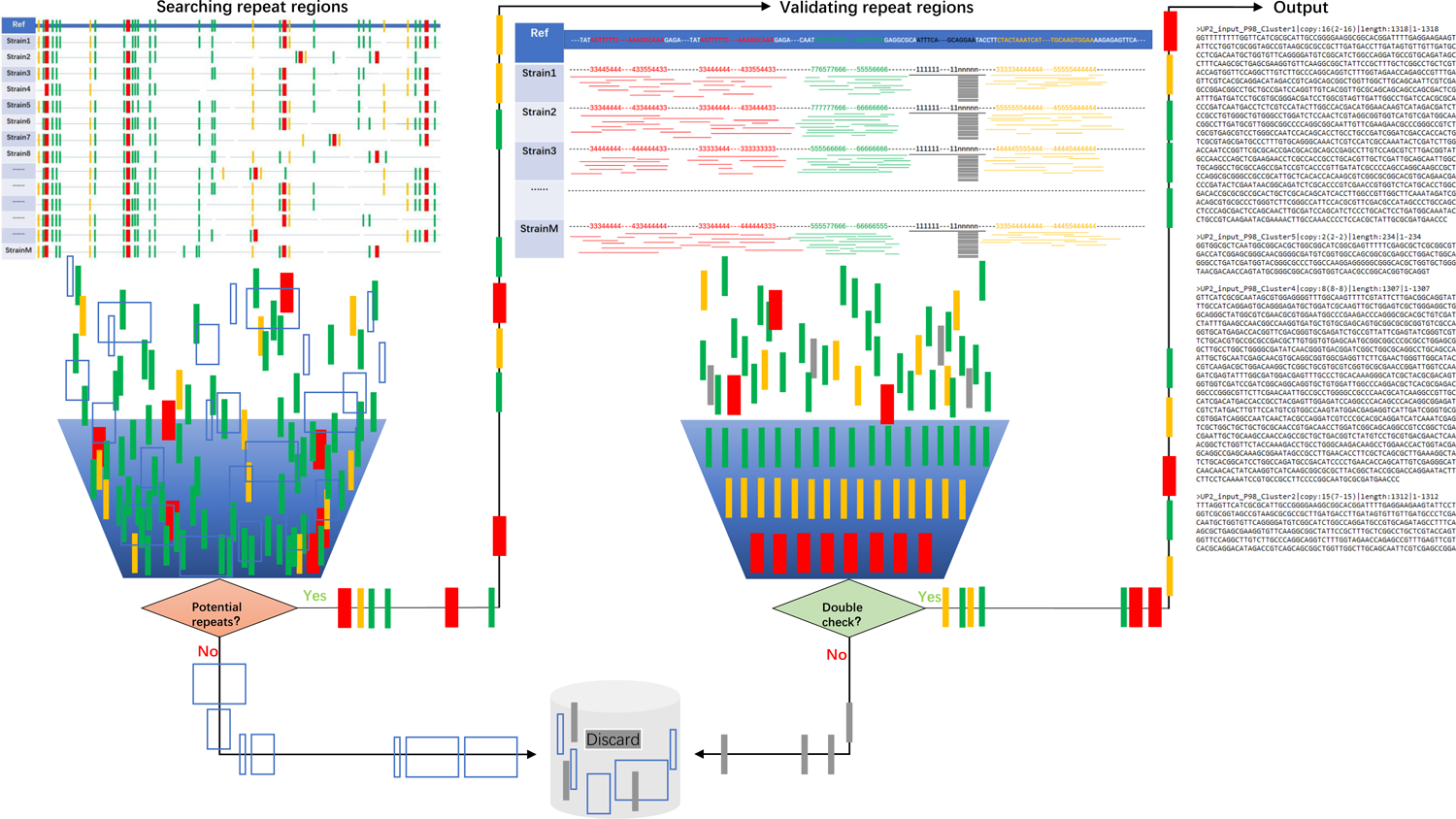
Illustration of the submodule preferentially searching for multicopy regions. 1) for searching candidate multicopy regions, internal alignments were performed on the microbial target fragments; 2) for verifying and obtaining the multicopy regions, including by determining the positions of each candidate multicopy region on the microorganism target fragments, obtaining the numbers of other candidate multicopy regions covering the positions of each base of the to-be-verified candidate multicopy regions, and calculating the median values of the copy numbers of the to-be-verified candidate multicopy regions.

To be conservative, the maximum possible values of the strain coverage rate were achieved with the fewest consensus sequences. All the logic modules were verified multiple times. As demonstrated in Table 2 and Supplementary Table1-8, the distribution of each cluster is summarized by the count numbers in parentheses for all the copy numbers out of the parentheses. The obvious choice was to pick the target cluster sequences as the design templates based on the percentage of strains and weighted averages of copy numbers, as detailed in the Materials and Methods. The templates of the primary-screened species-specific consensus sequence were designed, and the best sets of primers and probes were identified.

Finally, operational tasks could be submitted by providing the names of the target strains and the comparison strains or by uploading sequence files locally on the website https://bioinfo.liferiver.com.cn with easy registration. Users may select different configuration parameters depending on the purpose of the project. The configuration parameters mainly include the name of the workflow, target species, comparison species, uploaded local FASTA files, target fragment length, species specificity, repeated region similarity, target fragment strain distribution, host sequence filtering, priority scheme (prioritizing multicopy regions vs. prioritizing specific regions), calculation of target strain and alarm threshold similarities, and primer probe design parameters.

### Application and practice on a series of clinical projects

The most striking finding of this method was the contribution of specific, sensitive, and conservative biomarkers for each species or subspecies, especially those available for microbial genomes. First, the obvious advantage of our strategy was that it was capable of detecting species-specific or even subspecies-specific target fragments that contained forward primers, reverse primers and probes separately in several projects, such as those for human coronavirus HKU1 (HKU1), human coronavirus OC43 (OC43), human coronavirus NL63 (NL63), human coronavirus 229E (229E), Middle East respiratory syndrome (MERS) coronavirus, severe acute respiratory syndrome coronavirus (SARS-CoV) and severe acute respiratory syndrome coronavirus 2 (SARS-CoV-2). Notably, if there were no hits with the above biomarker genes or probes and no annotation, the sets were defined de novo, as presented in Table 1, and were obviously distinguished from other species or subspecies.

**Table 1.**
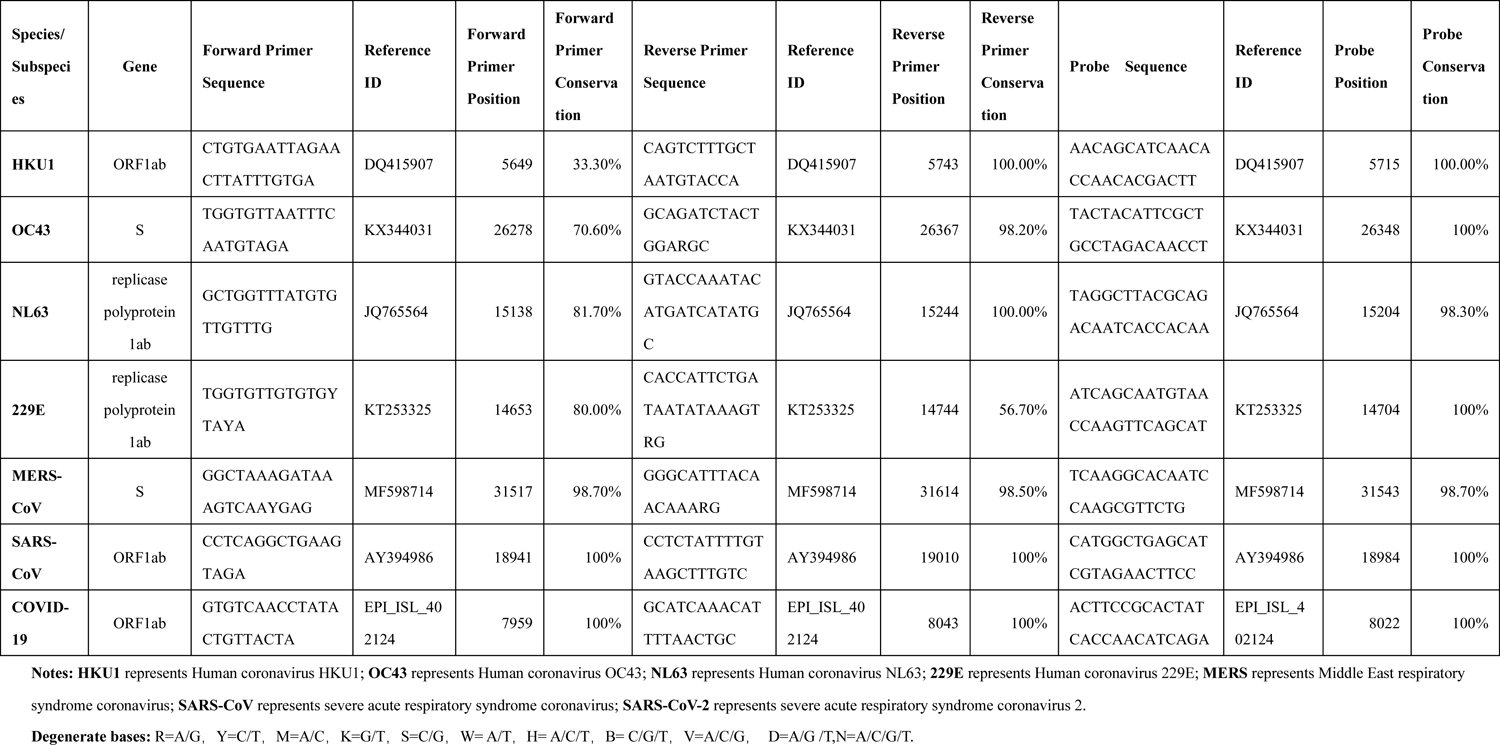
Final sets of primers and probes obtained by our strategy for different species or subspecies of Coronaviridae. HKU1 represents Human coronavirus HKU1; OC43 represents Human coronavirus OC43; NL63 represents Human coronavirus NL63; 229E represents Human coronavirus 229E; MERS represents Middle East respiratory syndrome coronavirus; SARS-CoV represents severe acute respiratory syndrome coronavirus; SARS-CoV-2 represents severe acute respiratory syndrome coronavirus 2.

Compared with the previous method, our strategy was highly accurate and sensitive, and undiscovered multicopy regions could be identified, as demonstrated in Table 2. For instance, IS6110 was identified by Shine as an insertion element that was found exclusively within members of the *Mycobacterium tuberculosis* complex (MTBC), Subsequently, IS6110 has become an important biomarker for the identification of MTBC species[25, 26]. Comparison with known 16s rRNA genes, i.e., IS1081 and PGRS of MTBC, IS1663 of *Bordetella pertussis* (BP), IS1001 of BP and *Bordetella parapertussis* (BPP), RepMP 4/5/1 of *Mycoplasma pneumonia* (MP), and cagA of *Helicobacter pylori* (HP) showed results that were all consistent with those of our study. Interestingly, IS1002 was present in both BP and BPP strains isolated from humans detected by Shine, inconsistent with the reported copy numbers in a recent study[27], in which only 47.2% of strains in BP and BPP had an IS1002-specific region, with 0.52 being the effective copy number in heterozygous taxa. When IS1002 was selected as the design template, false negative results were obtained that indicated that IS1002 could not cover all the BP and BPP strains. In addition, we identified several cases with false negative results; for example, the conservation of IS481 of BP and BPP was only 93.2%, with an effective copy number of 1.2. It was also clear that GBSil of *Streptococcus agalactiae* could not be used as the core template to cover all *S. agalactiae* strains, since the conservation of GBSil was only 28.6%.

**Table 2.**
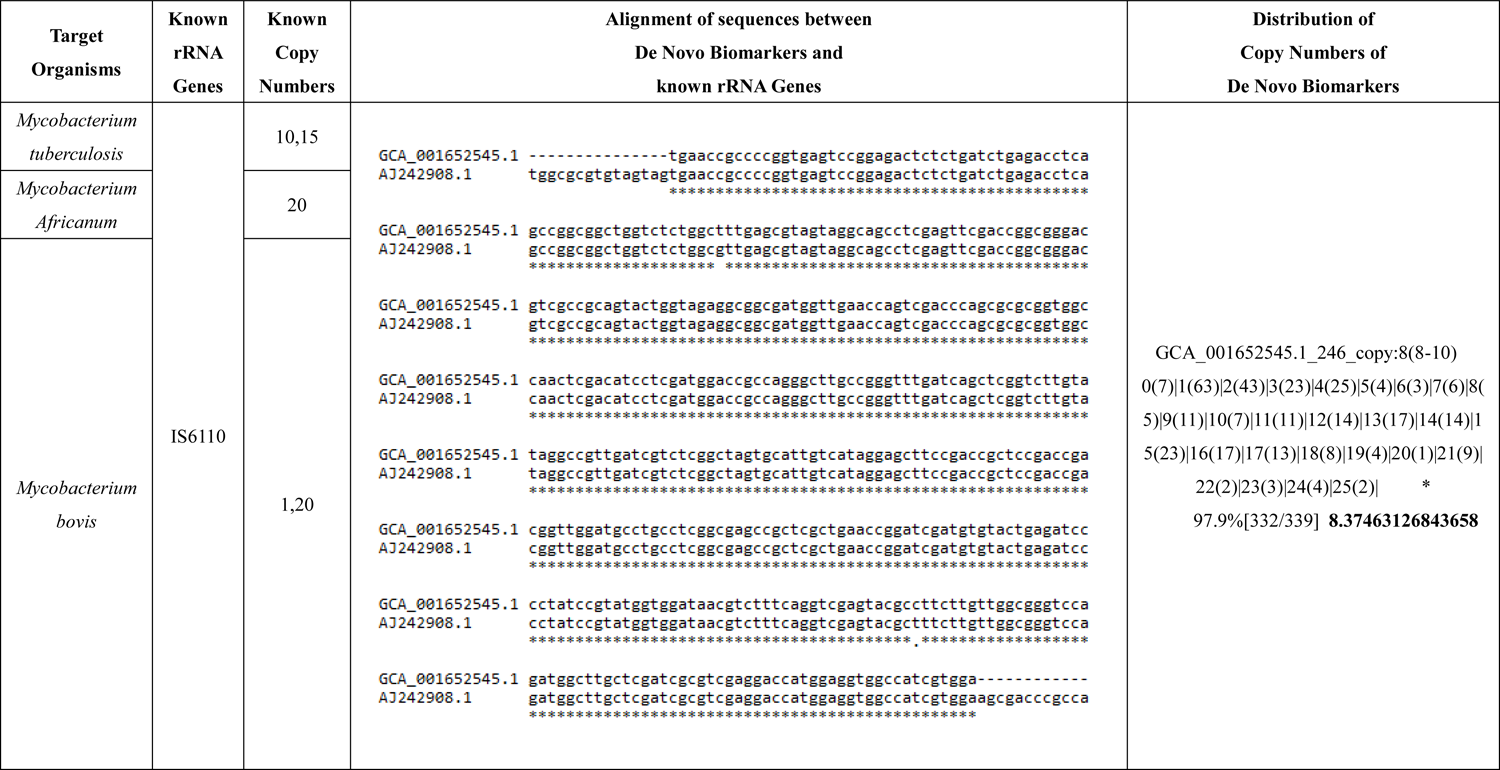
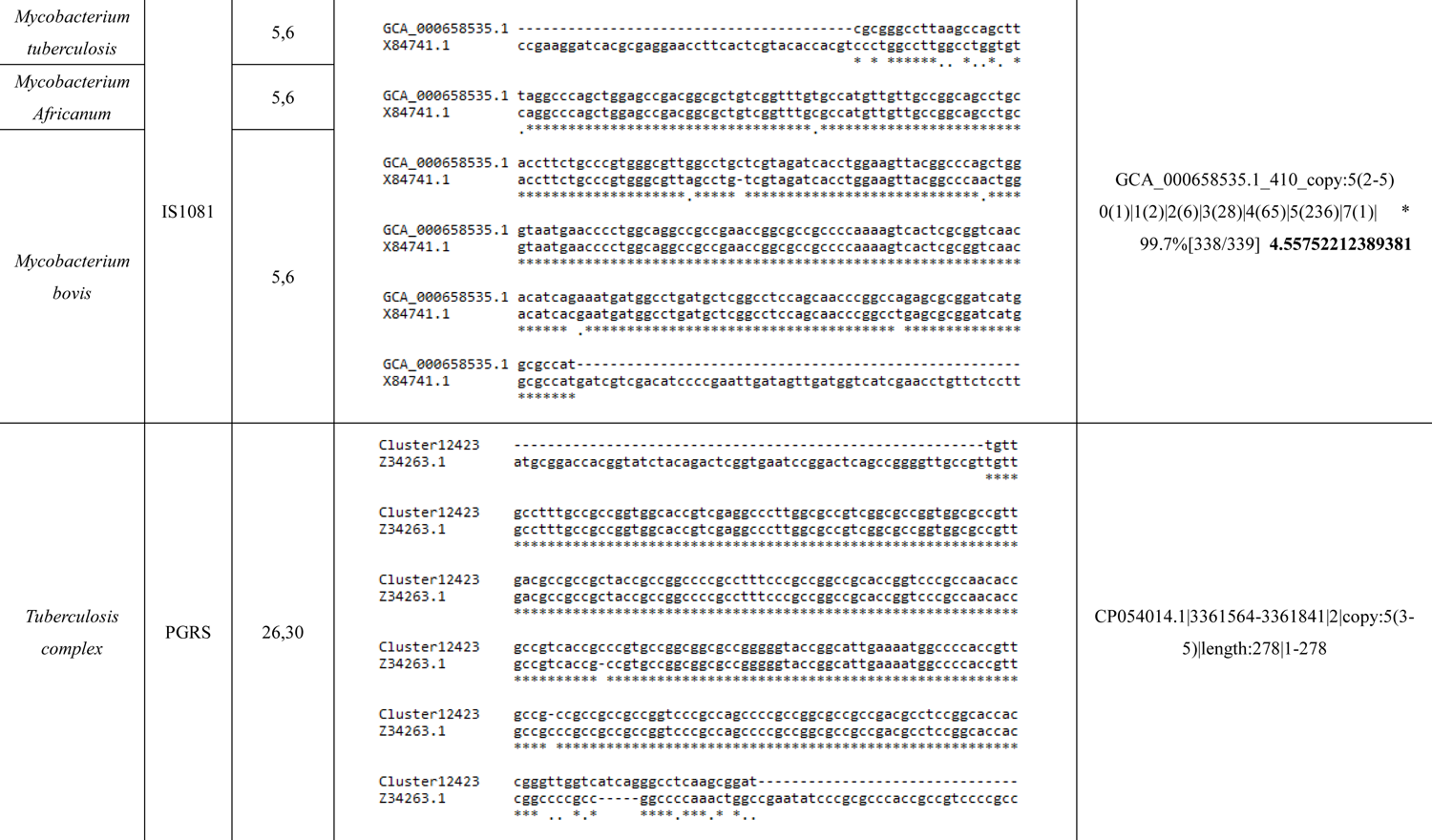
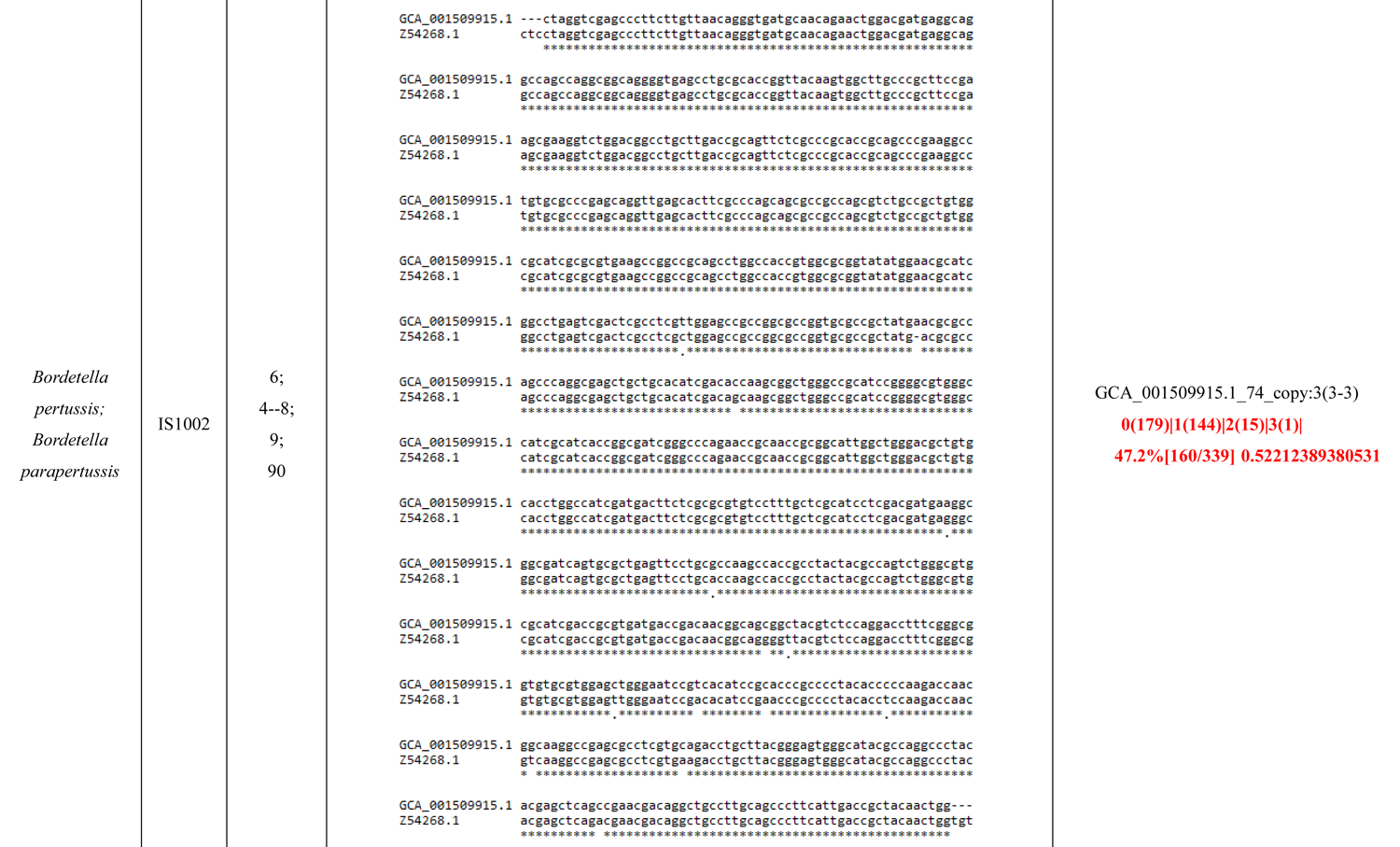
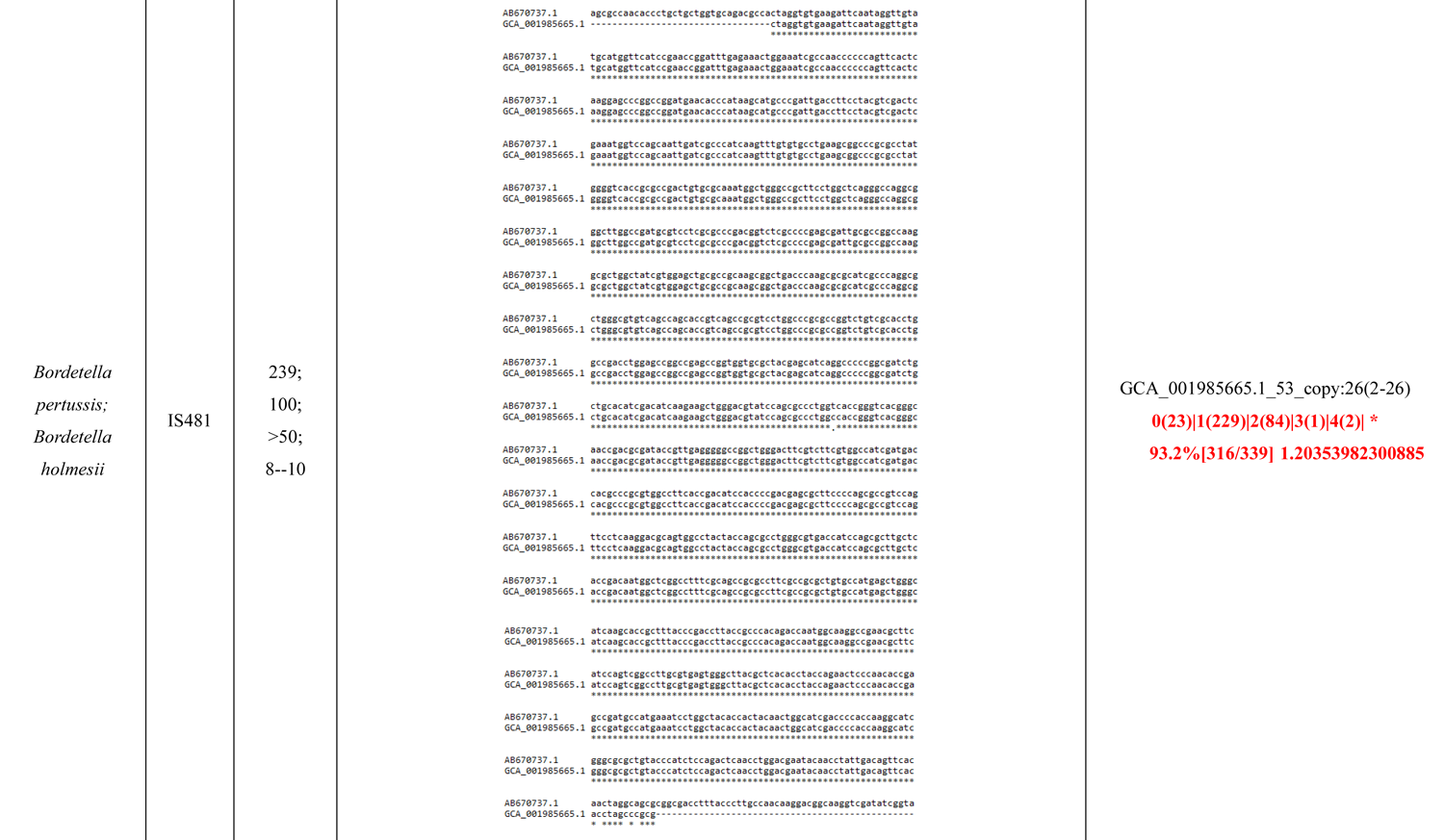
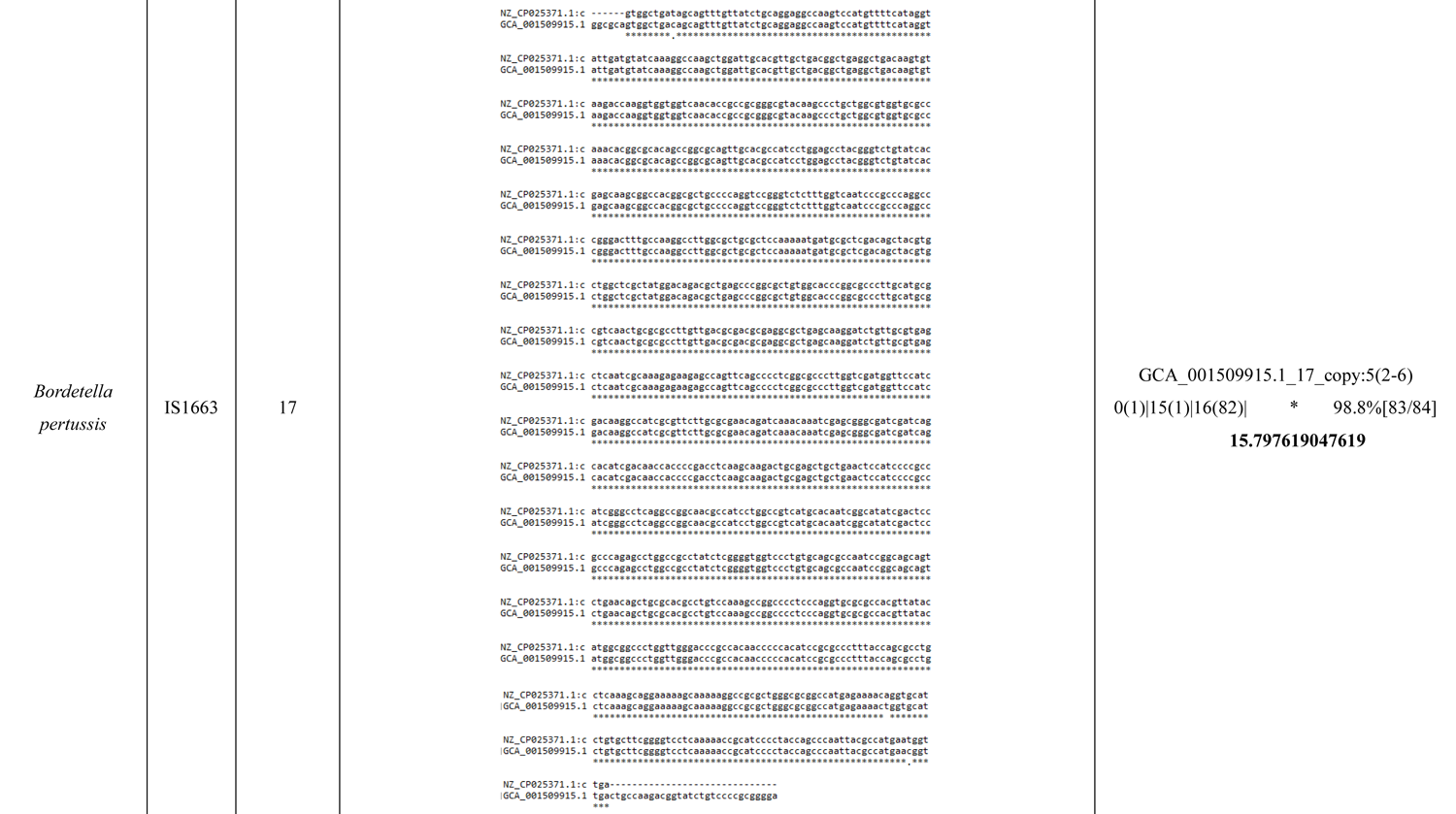
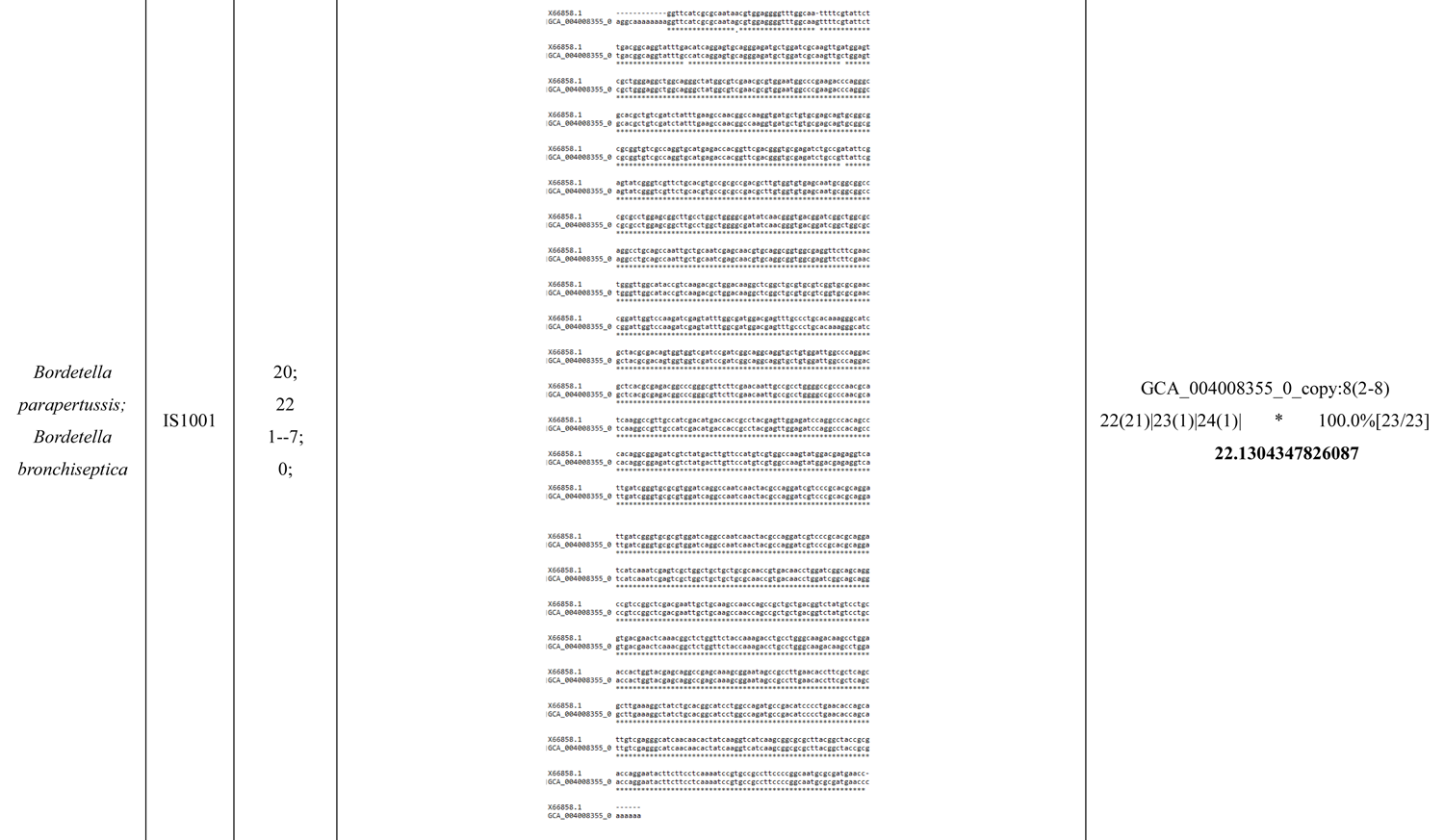
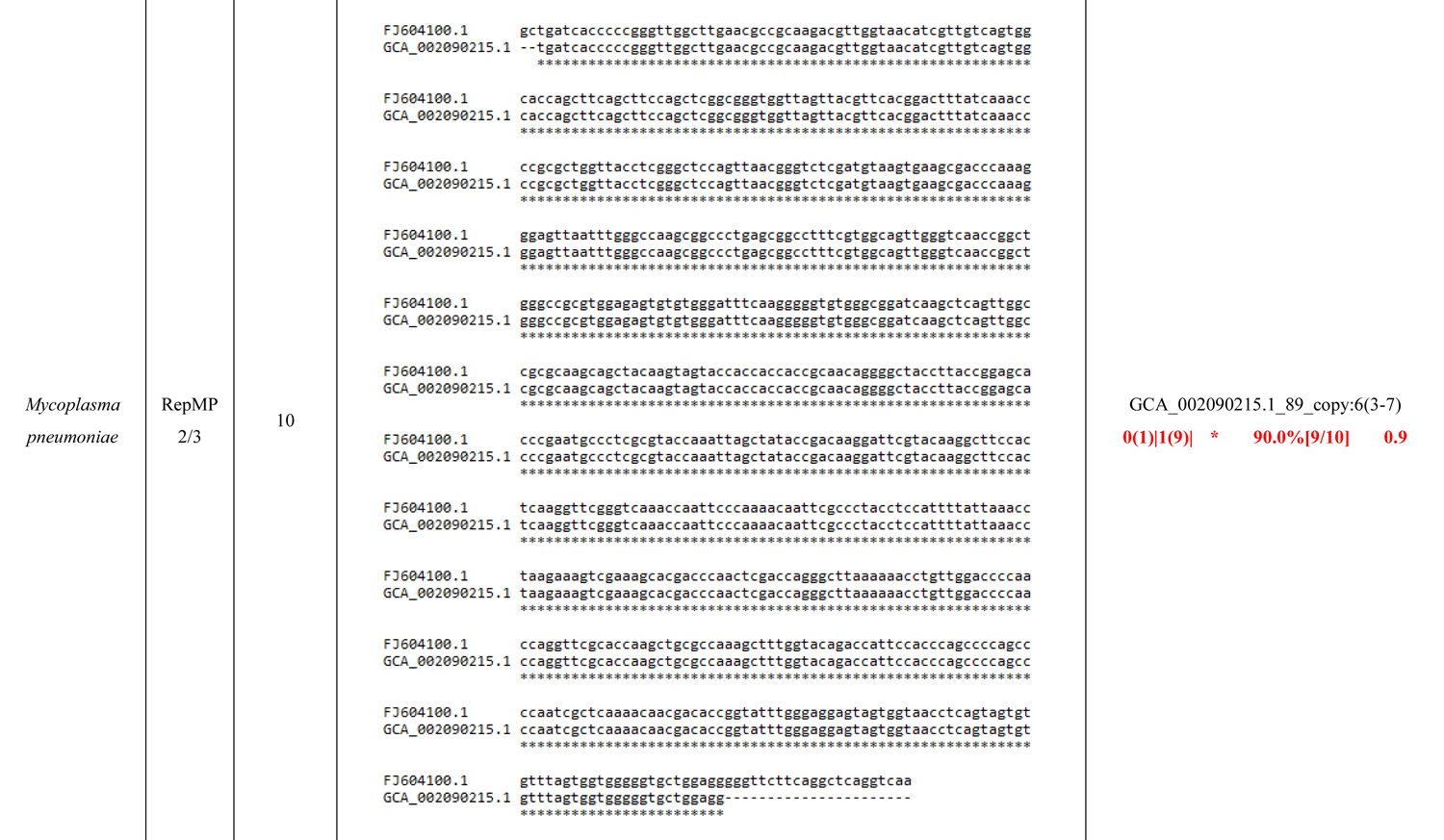
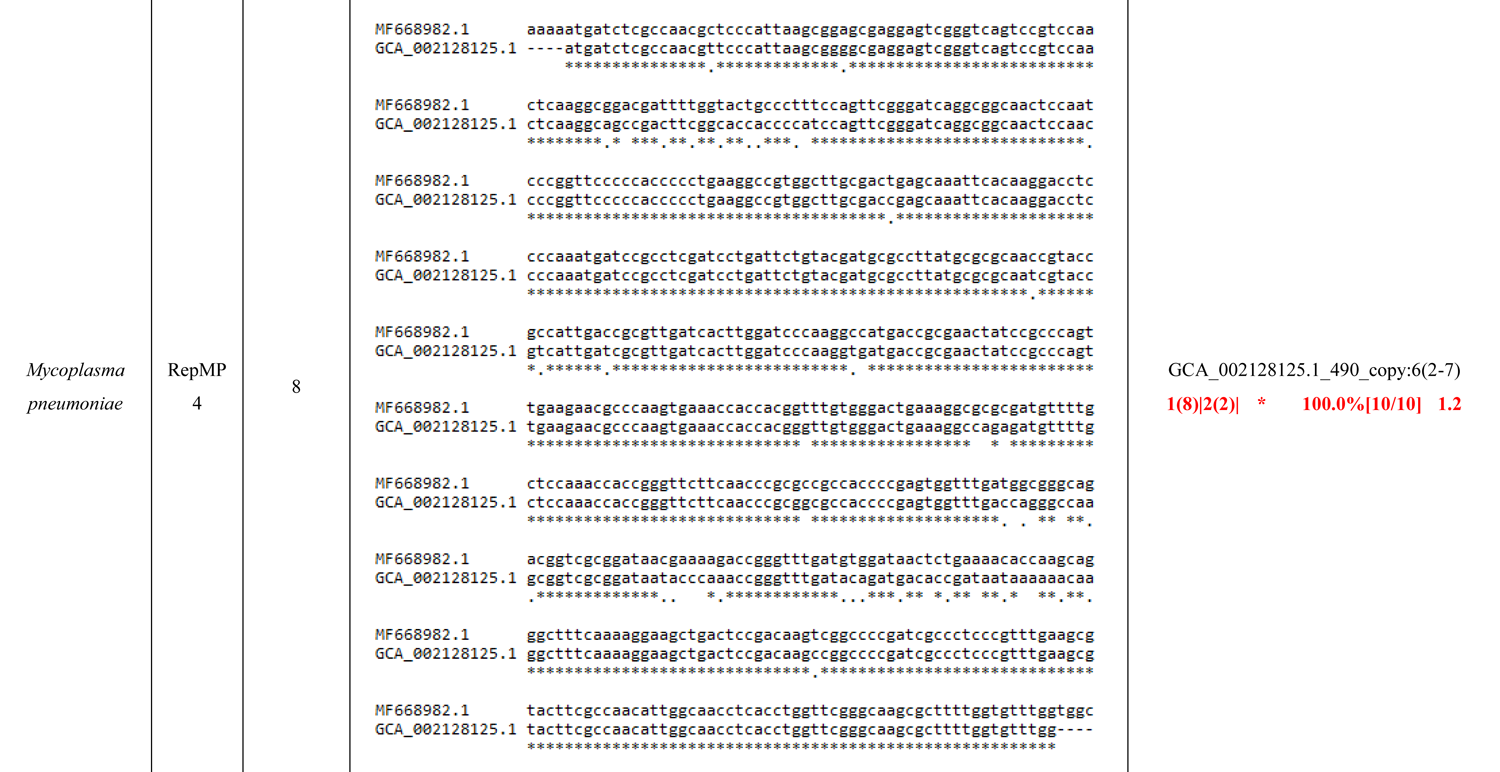
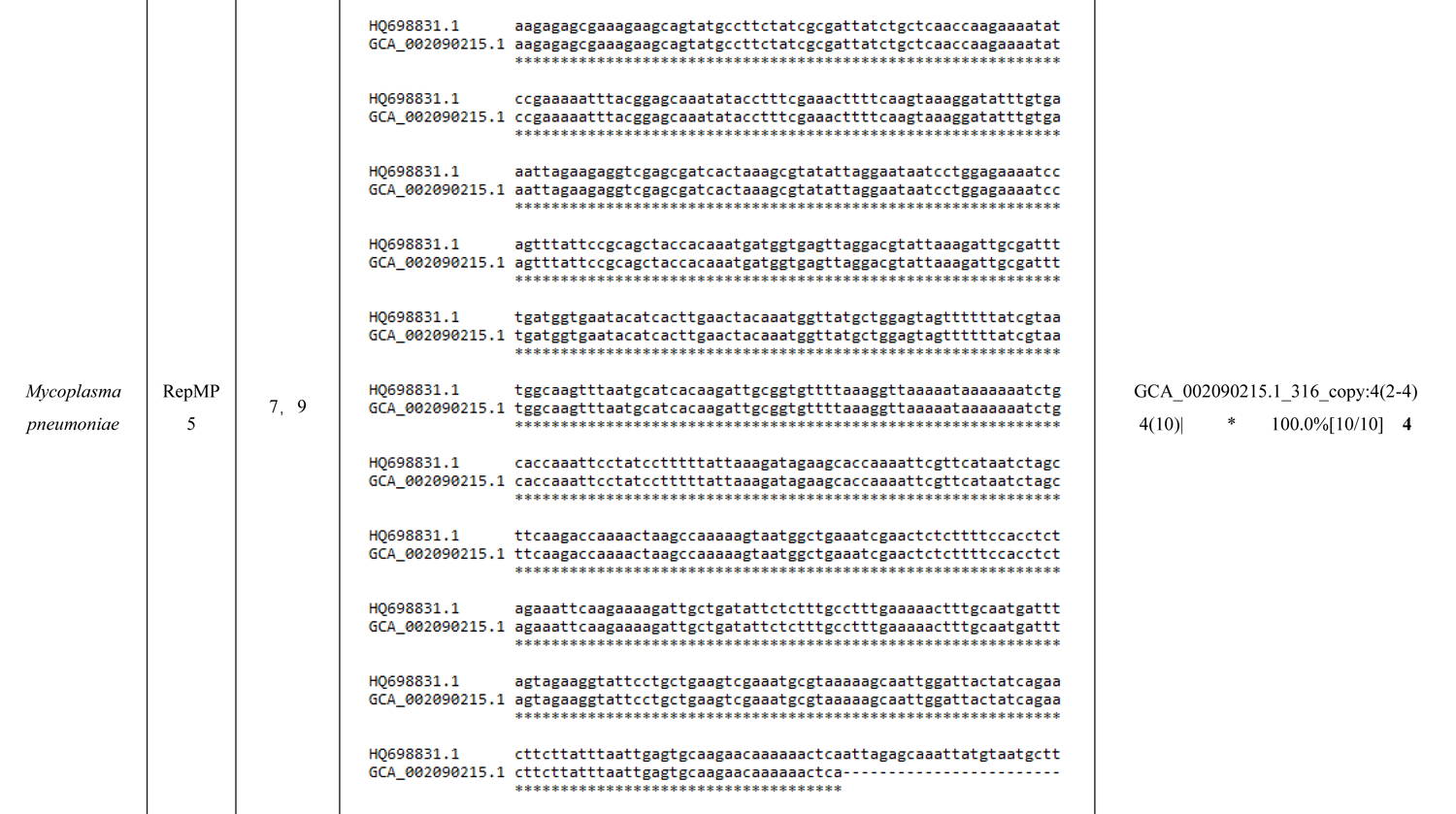
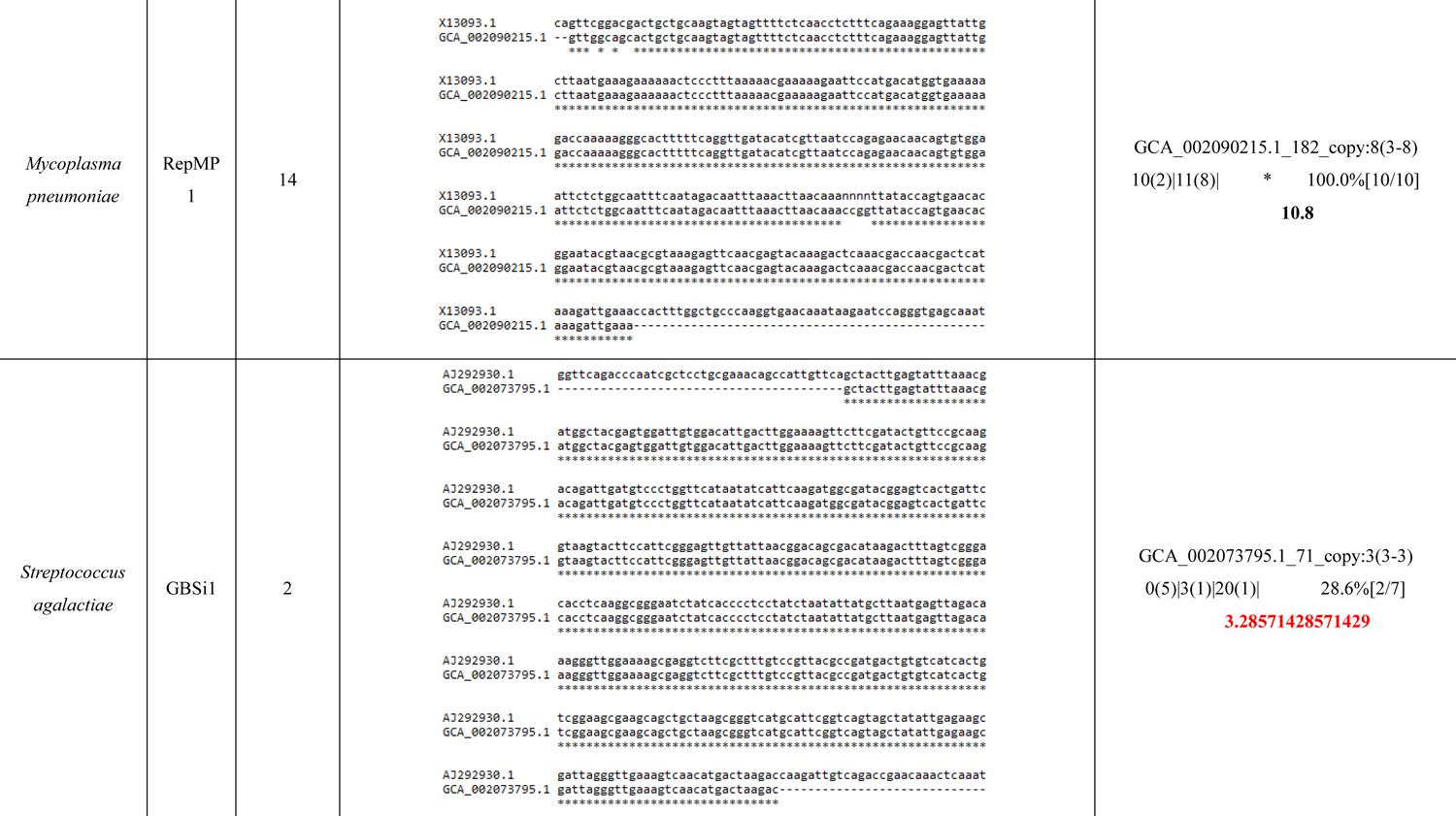
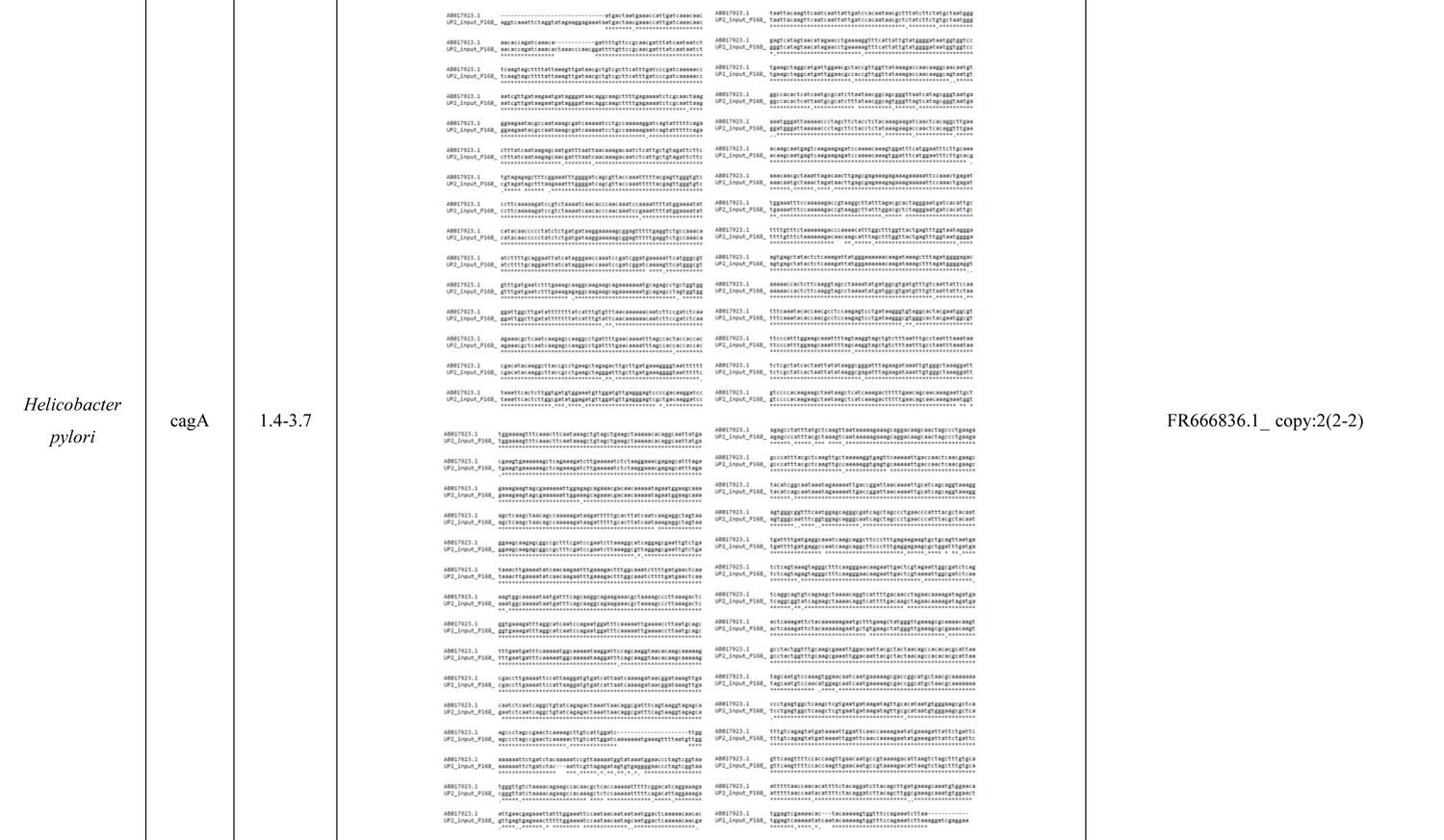
The performance of our strategy in reducing the false negative rate for sample sets of identified multicopy regions compared with known 16S rRNA genes. Here, the distribution of copy numbers of de novo biomarkers separately shows the sample ID; cluster ID; mean copy numbers (minimum copy numbers - maximum copy numbers); copy number corresponding to the i-th candidate species-specific consensus sequences (the number of strains with the i-th candidate species-specific consensus sequences); markers; conversation [summation of numbers of covered strains/total number of target strains]; and weighted average copy numbers.

Besides, we demonstrated the distribution of copy numbers of de novo biomarkers by the weighted average copy numbers and a series of sequences of newly discovered species- or subspecies-specific multicopy regions as potential design templates to avoid false positive results for any sequenced pathogenic organisms. It was clear that the performance of our strategy in reducing the false positive rate for the sample sets of newly discovered species- or subspecies-specific multicopy regions shown in Table 3, such as *M. tuberculosis, BP, BPP, MP, S. agalactiae, HP, Legionella pneumophila*, and *Candida auris.* This should be a very competitive strategy to identify newly discovered biomarkers.

**Table 3.**
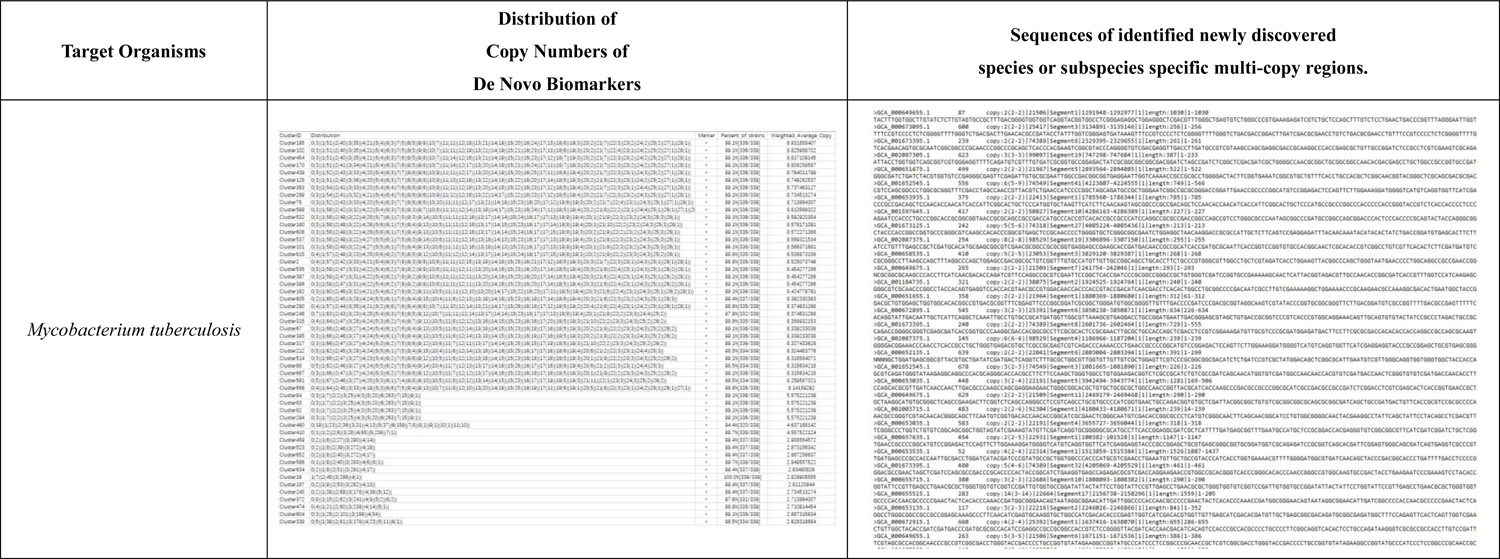
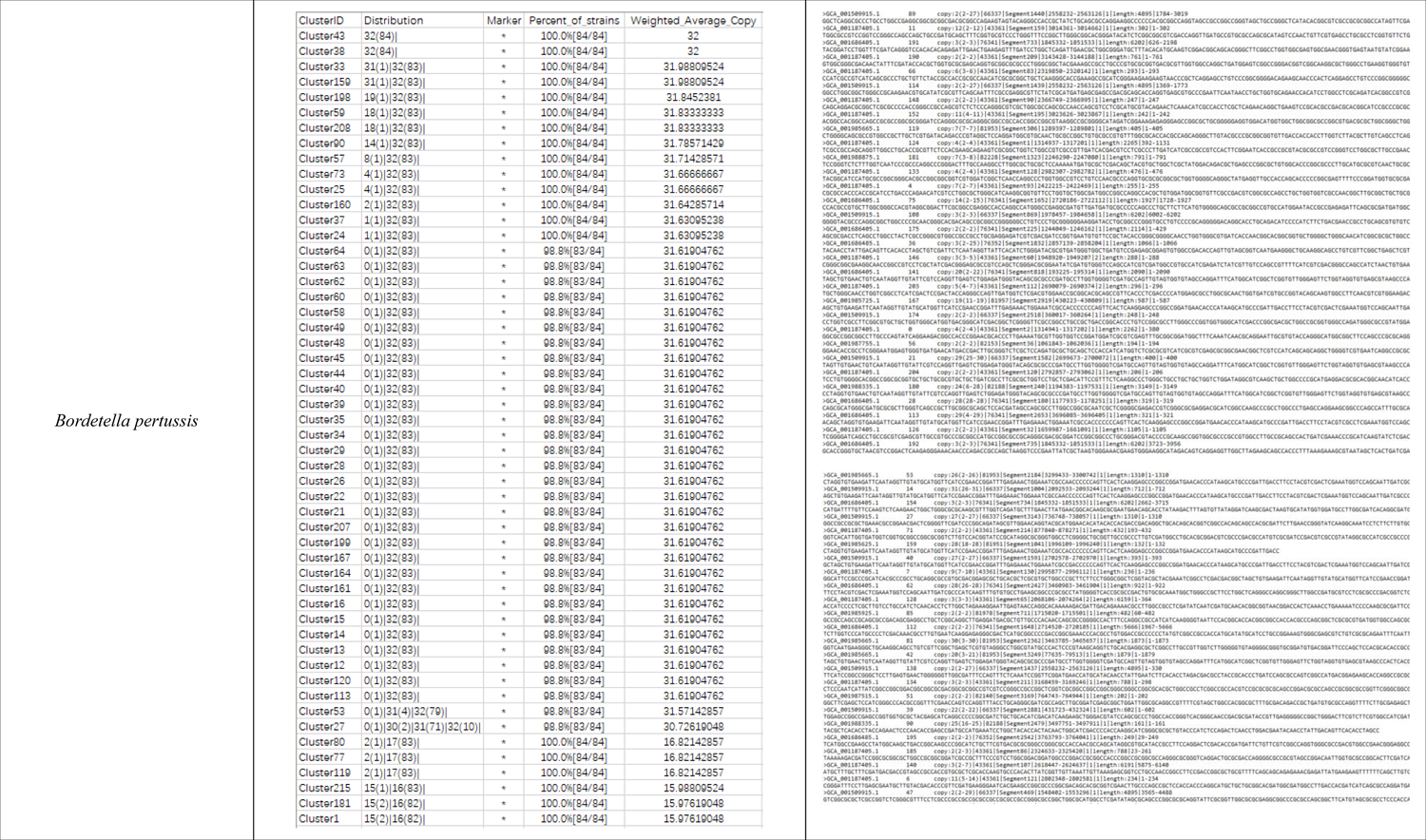
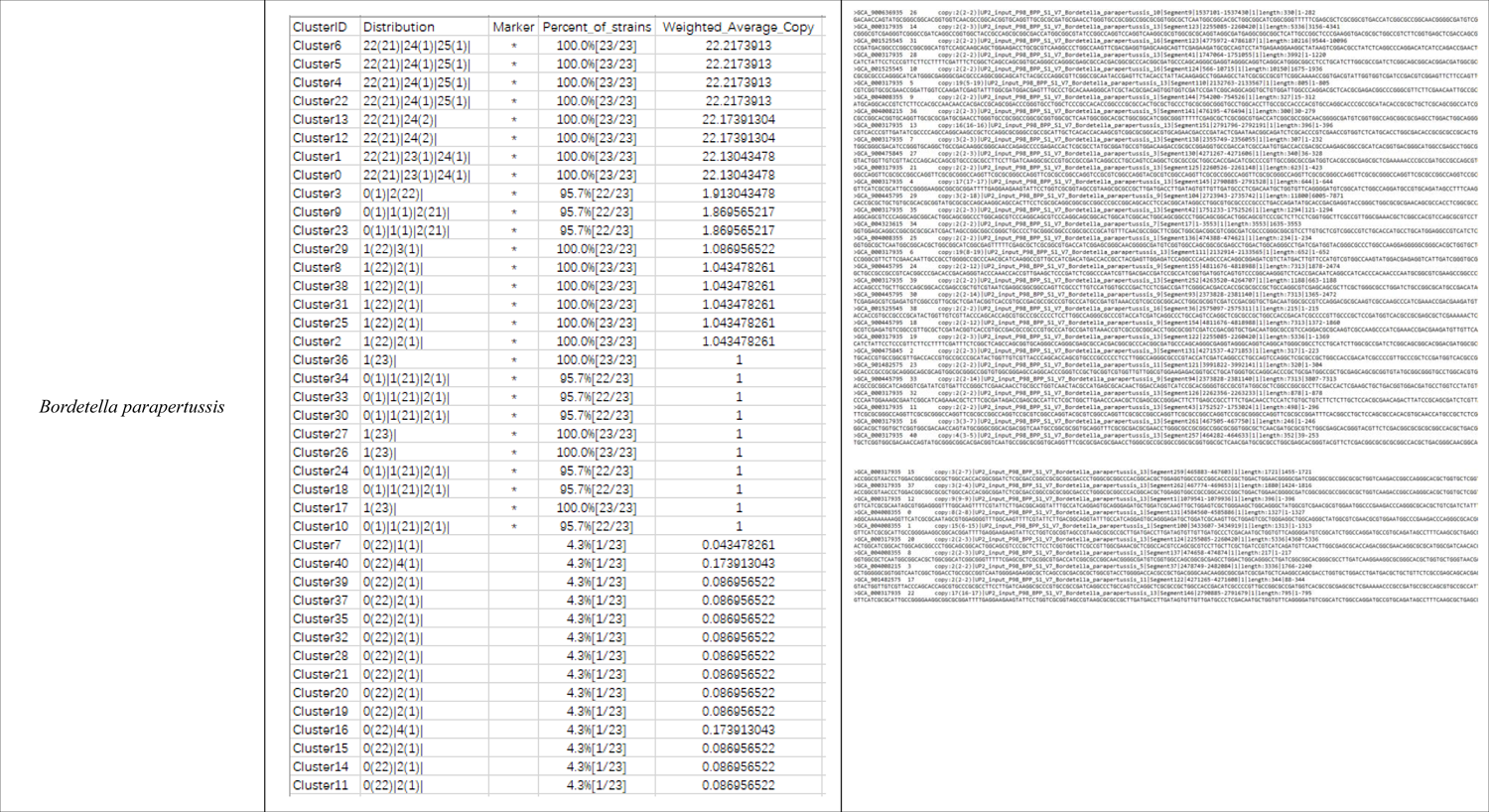
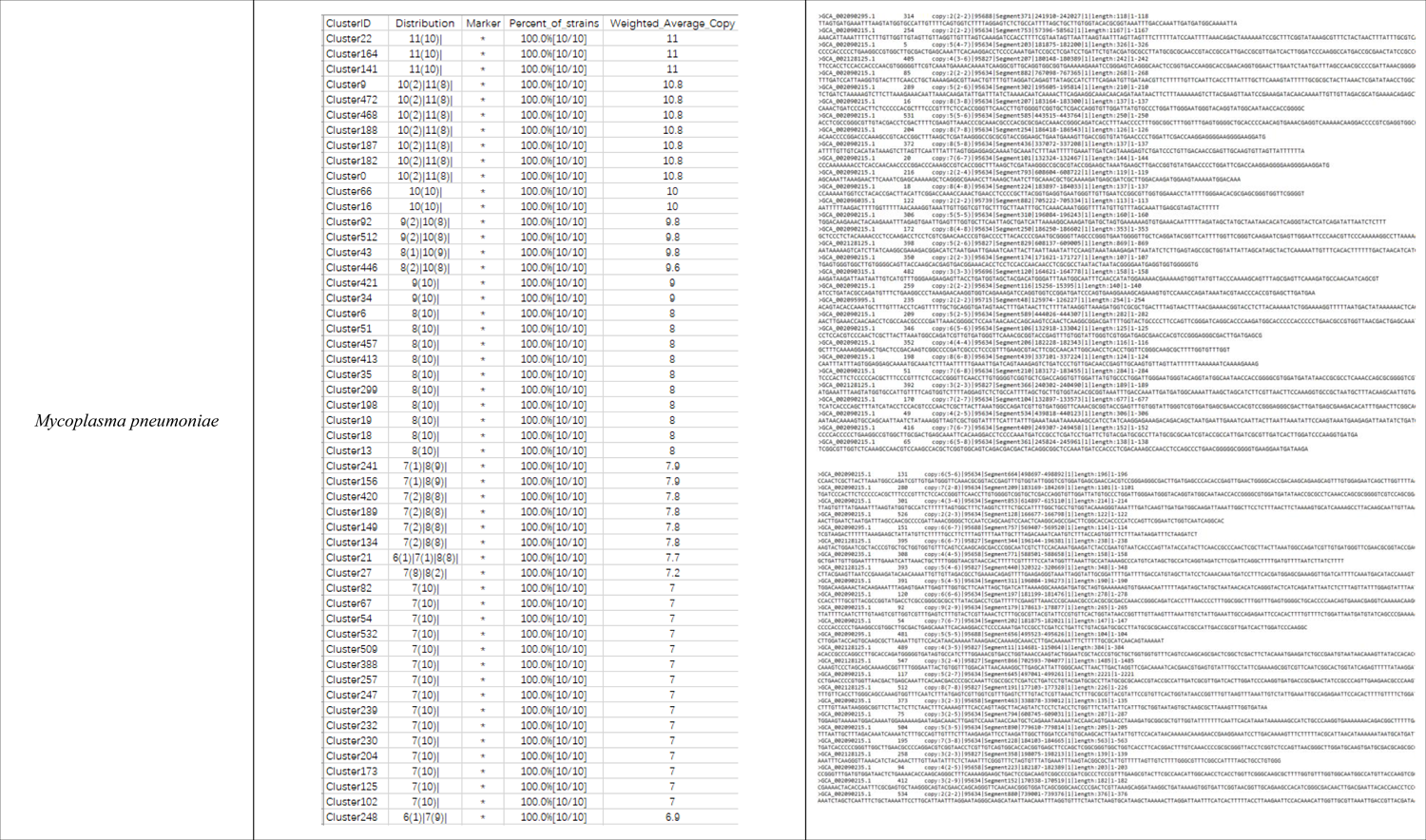
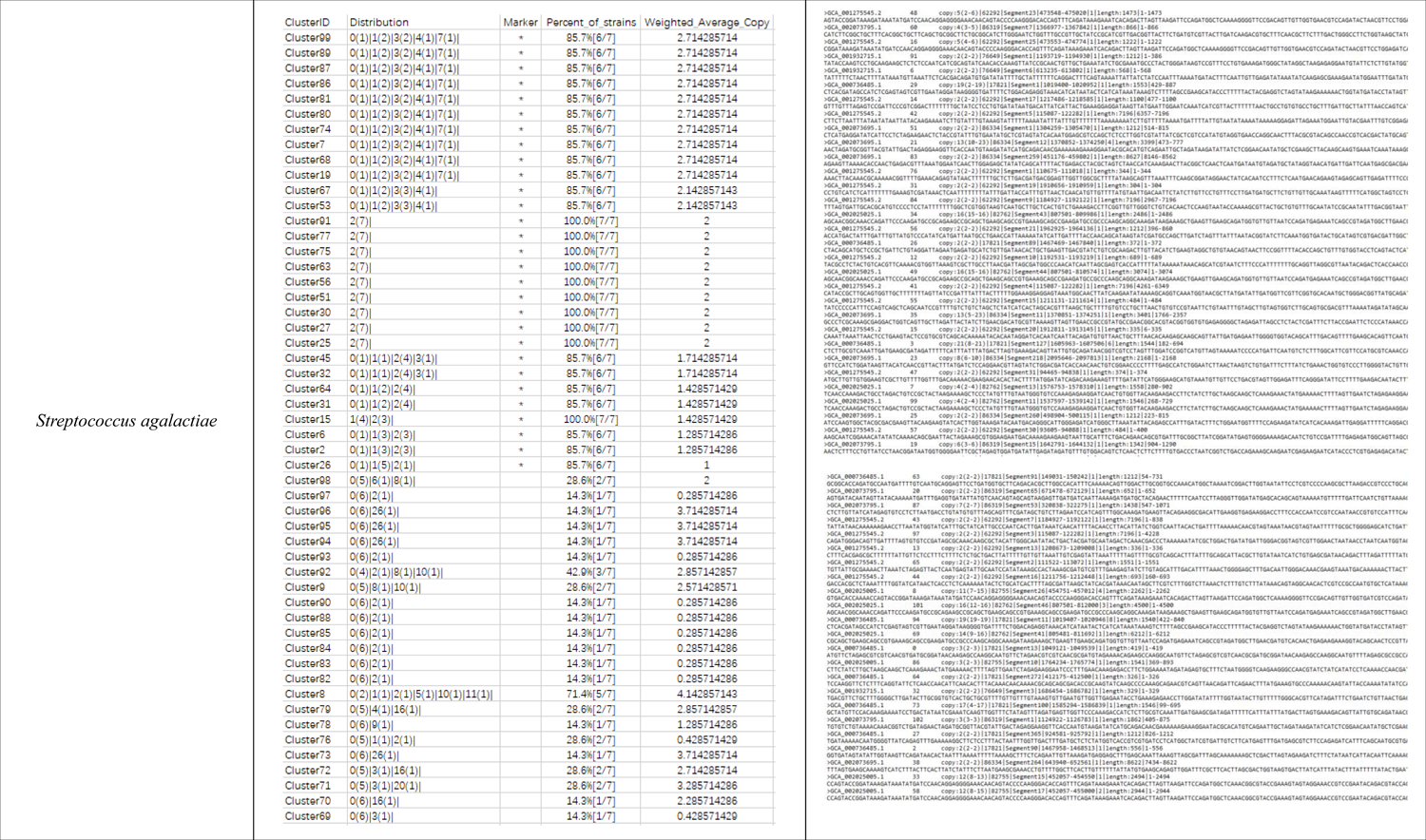
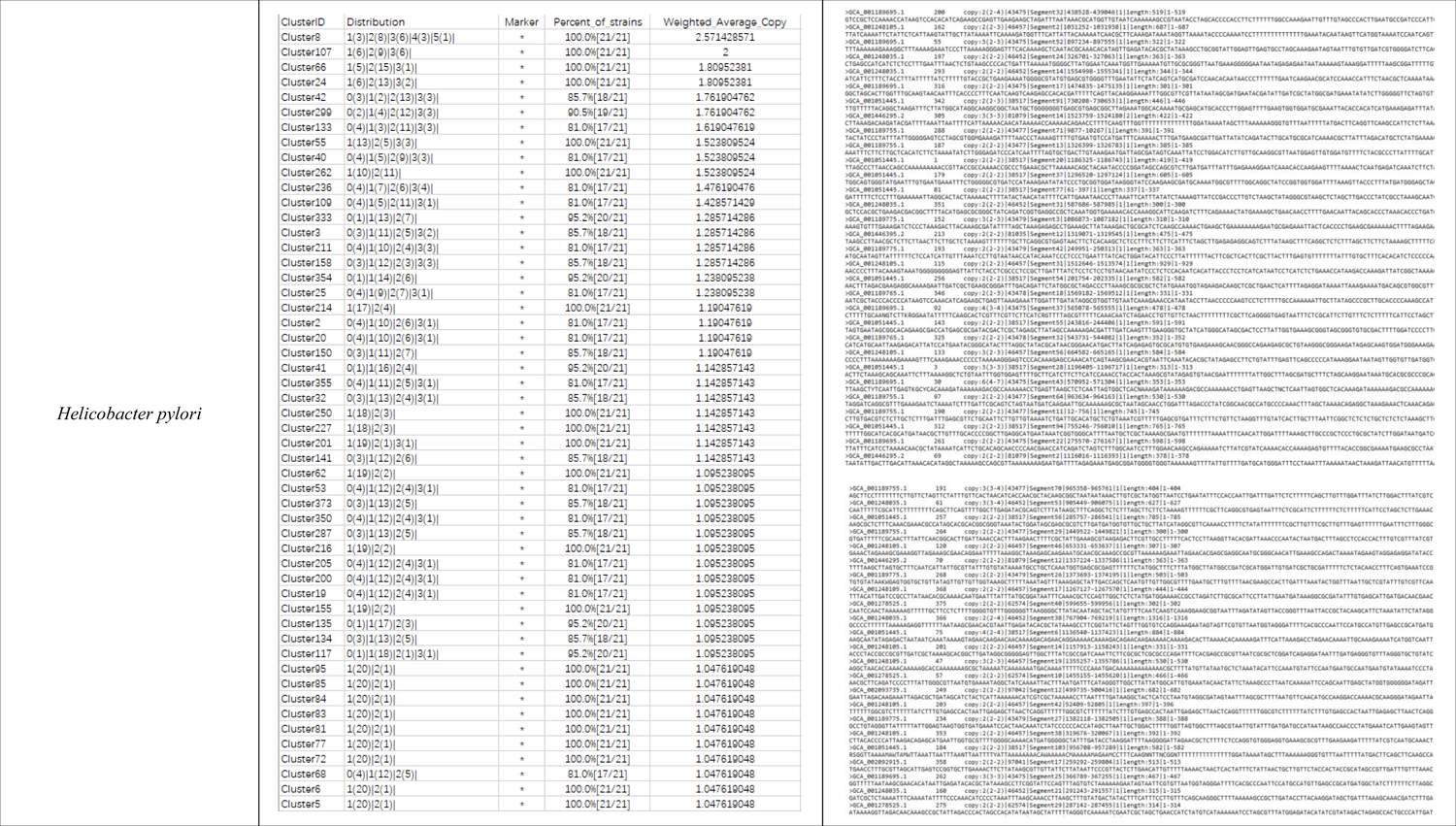
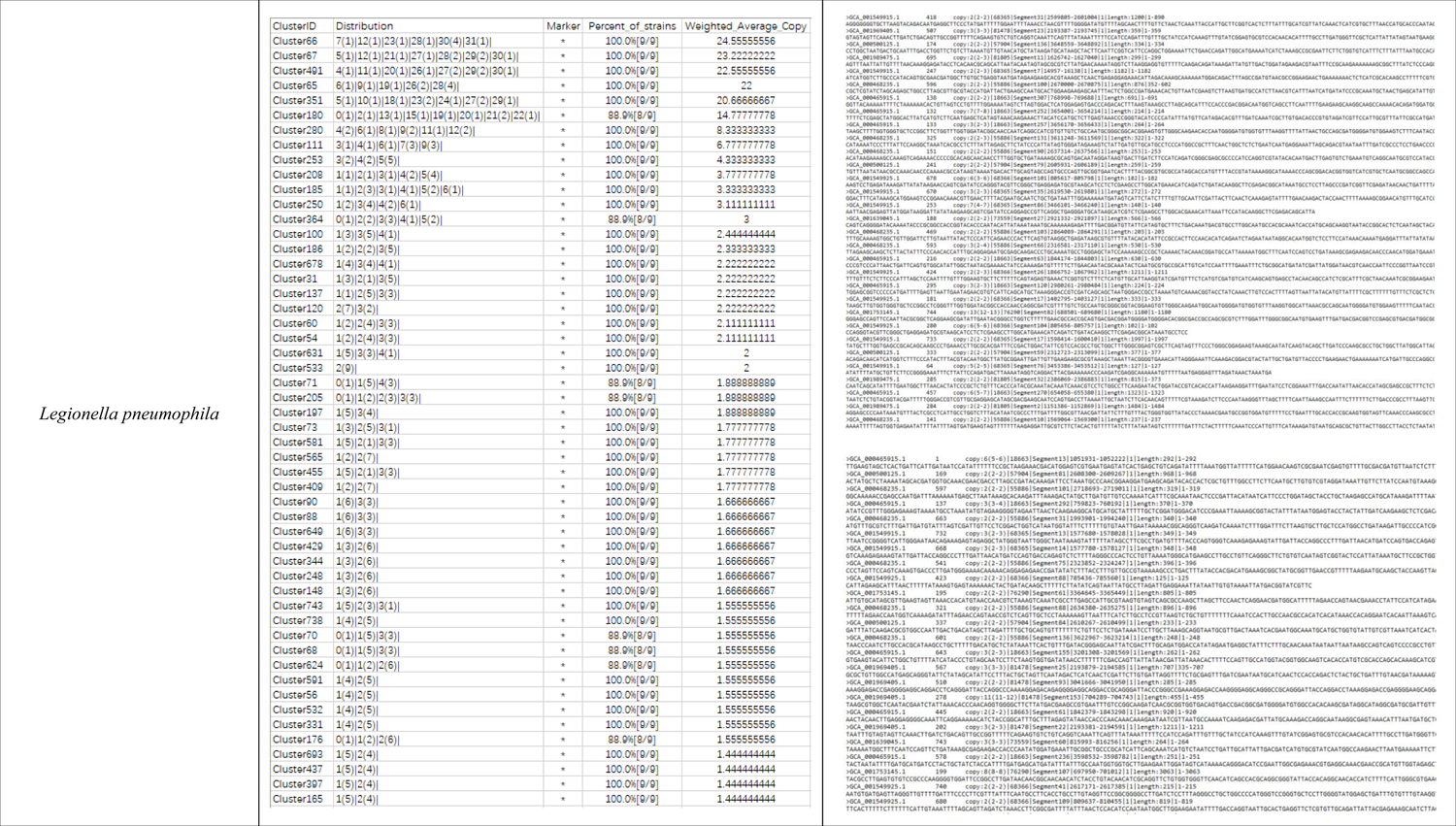
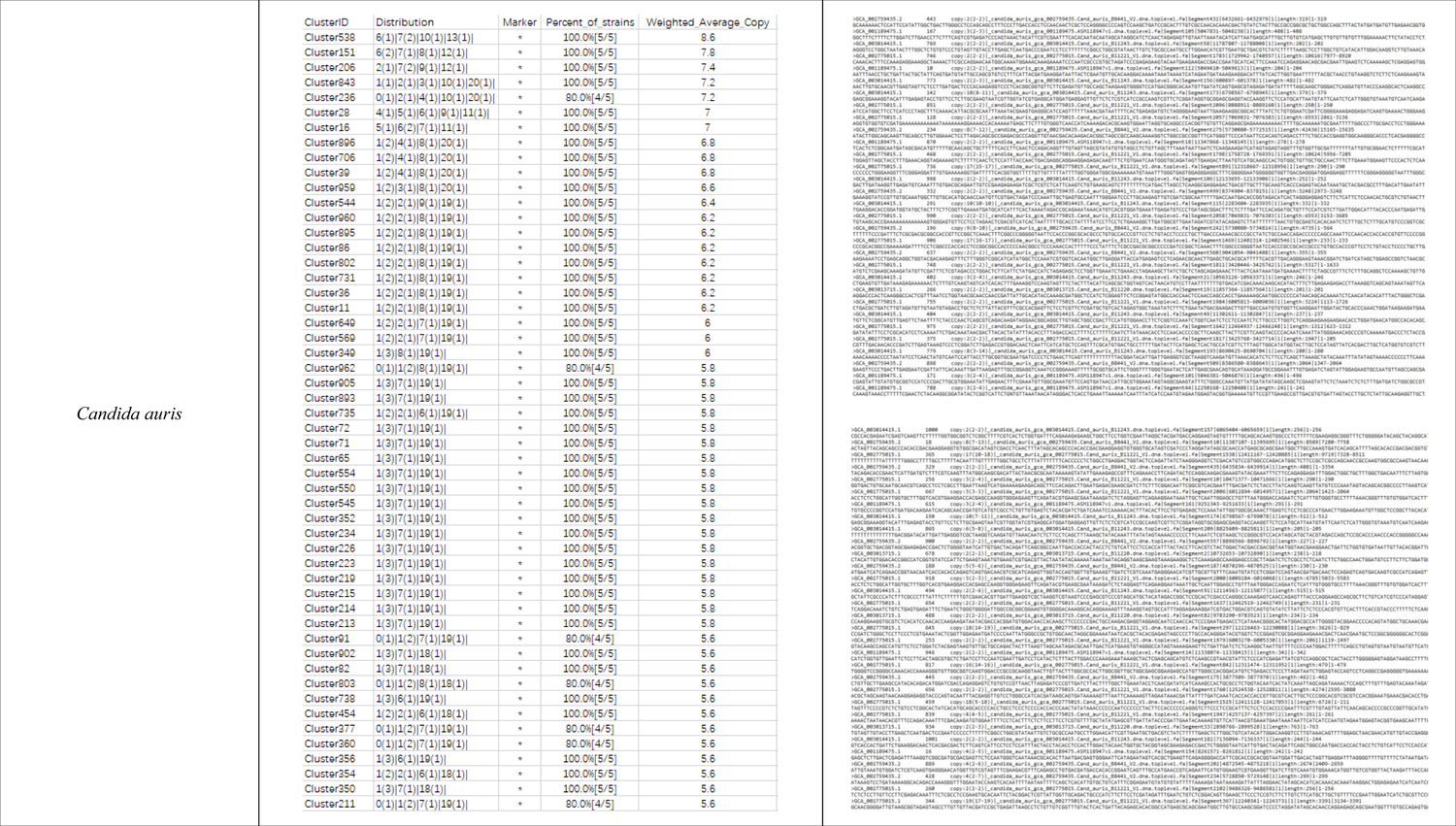
The performance of our strategy in reducing the false positive rate for sample sets of newly discovered species- or subspecies-specific multicopy regions. Here the distribution of copy numbers of de novo biomarkers separately shows the cluster ID; distribution; copy number corresponding to the i-th candidate species-specific consensus sequences (the number of strains with the i-th candidate species-specific consensus sequences); markers; percentage of strains [summation of numbers of covered strains/total number of target strains]; and weighted average copy numbers.

### Advantages of quantitative systematic and automated detection

Since 16S rRNA genes are not limited to whole-genome sequences that are not always multicopy, some rRNA genes in closely related species could not be distinguished from each other. Also, it is likely that not all plasmids have specificity and universality and are unevenly distributed across prokaryotic taxa. However, our method is highly accurate and sensitive and could be capable of detecting newly discovered multicopy universal species-specific and even subspecies-specific target fragments, covering all identified epidemic pathogenic microorganisms. 16s rRNA genes such as IS481, RepMP 2/3, and GBSil could not cover all pathogenic target microbial genomes to avoid producing false negative results and lowering the quality of the nucleic acid detection reagents presented in Table 2. However, we tried to explore a series of newly discovered specific, sensitive and conserved biomarkers from the database to improve the performance of clinical kits and reduce the false positive rates in Table 3. Overall, our study was suitable for identifying shared universal phylogenetic biomarkers with few false positive or false negative errors and automating the design of minimal primers and probes to detect pathogens in the community with cost-effective predictive power. In particular, this method was more comprehensive than limited selection of plasmids or 16S rRNA genes as template regions, as repetitive, specific and universal target fragments could be found even in incompletely assembled motifs in any case.

The detection device developed based on our strategy, Shine, was designed with quantitative PCR primers and probes for systematic and automated detection of pathogenic microorganisms in biological samples. Additionally, our strategy was more flexible with customized settings, which allowed the identification of the most conserved biomarkers, primers and probes via continuous updating of massive microbial genomic datasets. Users may submit the latest sequence dataset through a user-friendly interface. The sequence update coverage rate modules may reintegrate the latest sequence dataset into the database to calculate the coverage rates by recomparing the sequences of the original probes and primers to the updated sequences. This result could reflect whether the sequence of the original probes and primers could cover the newer strains.

The availability of genome annotation is not a limiting factor, and accurate detection with specific settings may cover all pathogenic microorganisms, including bacteria, viruses, fungi, amoebas, cryptosporidia, flagellates, microsporidia, piroplasma, plasmodia, toxoplasma, trichomonas and kinetoplastids. This method is also available for whole-genome sequencing data generated by new technologies such as third-generation sequencing.

## Discussion

To explore specific, sensitive and conserved biomarkers from massive microbial genomic data within populations to improve detection sensitivity and accuracy, several clinical projects have been carried out by using devices based on Shine. Unfortunately, it should be noted that this study examined only limited public genomic data, and we are still looking forward to promoting collaboration with more organizations for open sharing of data and with respect for all rights and interests[28]. Assuming that the more comprehensive genomic data are, the more effective the detection biomarkers. The importance of identifying specific regions in microbial target fragments cannot be overestimated. The biodiversity and evolution of vertebrate RNA viruses has expanded dramatically since the beginning of the millennium, and it has been reported that more expensive and better sampling worldwide and more powerful approaches for virus characterization are needed to help us find these divergent viruses, such as chuviruses and jingmenviruses[29], which will help fill the gaps in the knowledge of RNA virus evolution[30]. With the development of methods for detecting more than 100 different nucleic acid targets simultaneously, FilmArray made the system well suited for the molecular detection of infectious agents, and the automated identification of pathogens from their corresponding target amplicons could be accomplished by analysis of the DNA melting curve of the amplicon[31]. Additionally, several studies have reported multiplex real-time PCR assays for detecting four microorganisms relevant to community-acquired pneumonia (CAP) infections[32] in Asia; CAP is one of the most common infectious diseases and a significant cause of mortality and morbidity globally. The availability of tests with improved diagnostic capabilities will potentially lead to the ability to make informed choices regarding antibiotic usage and appropriate management of patients to achieve better treatment outcomes and financial savings[32].

Herein, we generated a more significant biomarker dataset, which was validated by several clinical experiments, as described in Table 1-3. All the results indicate that our strategy is robust for detecting effective biomarkers, which seems to indicate that the specificity could account for this performance. Hypothesized that the specific biomarkers from different pathogens are identified, later they may be sensed by graphene arrays. Interestingly, graphene is a lightweight, chemically stable and conductive material that can be successfully utilized for the detection of various virus strains. The current state-of-the-art applications of graphene-based systems for sensing a variety of viruses, e.g., SARS coronavirus 2 (SARS-CoV-2), influenza, dengue fever, hepatitis C virus, human immunodeficiency virus (HIV), rotavirus and Zika virus, have been summarized previously[33, 34]. That was to say, graphene-based biosensor technology with high sensitivity and specificity could be particularly useful in the life sciences and medicine since it can significantly enhance patient care, early disease diagnosis and pathogen detection in clinical practice[35, 36]. Furthermore, CRISPR-Cas systems, in particular the recently discovered DNA-targeting Cas12 and RNA-targeting Cas13 systems, both possessing unique trans-cleavage activity, are being harnessed for viral diagnostics and therapies[37]. In addition, specific high-sensitivity enzymatic reporter unlocking (SHERLOCK) testing in one pot (STOP) is a streamlined assay combining simplified extraction of viral RNA with isothermal amplification and clustered regularly interspaced short palindromic repeats (CRISPR)-mediated detection, which could be performed at a single temperature in less than one hour with minimal equipment[38]. As a consequence, we tentatively propose cooperating with related institutes to combine the Shine strategy with graphene-based biosensor technology or CRISPR-Cas systems for application in pathogen sensing.

The method ensured coverage of all pathogenic microbial genomes and effectively identify the specific species or subspecies to avoid lowering the quality of nucleic acid detection reagents, but of course, there were exceptions, i.e., highly divergent viruses, such as Sapovirus and human astrovirus, which have limited consensus biomarkers with high performance. If none of the strain coverage rates of the candidate consensus sequences reached the preset value, we had to prioritize specificity and/or sensitivity and combine the candidate consensus sequences to improve conservation, although this may not be cost-effective and could cause several experimental errors. The recommended process was in turn performed by screening the combinations with the strain coverage rate reaching the preset values and having the fewest consensus sequences, taking the screened combinations as the candidate consensus sequences, and then verifying/obtaining the primary-screened species-specific consensus sequences. The combination could be screened according to the number of consensus sequences from low to high, for selection. Unless a single consensus sequence covered all the current strains, it was possible to find two consensus sequences for which the sum of the strain coverage rates of the two consensus sequences was greater than or equal to the pre-set value of the strain coverage rate. If so, two consensus sequences were recorded in the results; if not, three consensus sequences were combined. That is, unless there was a single consensus sequence or two consensus sequences that could meet the preset value of the strain coverage rate, it was possible to find three consensus sequences, where the sum of the strain coverage rates of the three consensus sequences was greater than or equal to the preset value of the strain coverage rate. If so, the three consensus sequences were recorded in the results; if not, four consensus sequences were combined. Based on this analogy, infinite numbers of consensus sequences should not be combined until the consensus sequence combination that can meet the preset value of the total strain coverage rate is found and recorded in the result. Our strategy has been successfully applied to a number of clinical projects and has the overwhelming advantages of quantitative systematic and automated detection of all pathogenic microorganisms without limits of genome annotation and incompletely assembled motifs described above.

Based on the incomplete motifs, there was no specific restriction on the order in which the motifs were connected together, i.e., the motifs could be connected to the chain in random order. If the region of similarity met the preset value contained different motifs, it was divided based on the original motif connection points into different subregions to determine whether the subregions were candidate multicopy regions. Compared with 16s rRNA, de novo biomarkers based on weighted average copy numbers and a series of sequences of newly discovered species- or subspecies-specific multicopy regions could be identified as potential design templates to avoid false positive results for sequenced pathogenic organisms.

In addition to the above characteristics, this approach involves obtaining species-specific consensus sequences for microorganisms. Were these different assignments due to the fundamental nature of the approach or the result of different approaches to species demarcation by the respective specialized study groups (SGs)? For instance, HIV-1 and HIV-2 were assigned to two different species, while SARS-CoV and SARS-CoV-2 were assigned to two strains of a single species. That is, how can the position of the viral entity in the natural world be defined? In practical terms, recognizing virus species as the principal subjects of virology would also expand the scale of the spatiotemporal framework connecting studies of natural virus variation, cross-host transmission, and pathogenicity and thus contribute to the understanding and control of virus infections[39].

## Materials and Methods

### Acquisition of raw data

The pathogenic genomic data were derived from public databases, such as the National Center for Biotechnology Information (NCBI) Assembly database[40], Global Initiative on Sharing All Influenza Data (GISAID)[28, 41], ChunLab’s public data and analytics portal (EzBioCloud)[42], Eukaryotic Pathogen Database Project (EuPathDB)[43], Giardia Genomics Resources (GiardiaDB)[44], Trichomonas Informatics Resources (TrichDB)[44], Fungal & Oomycete Informatics Resources (FungiDB)[45], and Pathosystems Resource Integration Center database (PATRIC) [46], which contained either completely assembled pathogenic genomes or incompletely assembled motifs. The defined populations were specific species or subspecies, and the control group was all the other species or subspecies of the same classification excluding the defined populations. The method further involved comparing selected adjacent microorganism target fragments one to one; if the similarity after comparison was lower than the preset values, an alert was issued, and screening conditions corresponding to the target strains were displayed. Abnormal data and redundant data introduced by human errors could be filtered. The target fragments of microorganisms could be the whole genomes of microorganisms or their gene fragments.

### Searching for the specific regions

As shown in Figure 2, to identify the specific regions in the microbial target fragments, 1) the microbial target fragments were compared with the whole-genome sequences of one or more comparison strains one to one, and fragments for which the similarity exceeded the preset value were removed to obtain several residual fragments as the first-round cut fragments T1-Tn, wherein n is an integer greater than or equal to 1; 2) then, the first-round cut fragments T1-Tn were compared with whole-genome sequences of the remaining comparison strains, and fragments for which the similarity exceeded the preset values were removed to obtain the collection of residual cut fragments as the candidate specific regions of the microbial target fragments; and 3) the specific regions were then verified and obtained to determine whether the candidate specific regions met the criteria for the following steps: a) searching in public databases[47] to find whether there were other species for which the similarity to the candidate specific region was greater than the preset value; b) comparing the candidate specific regions of the whole-genome sequences with those of the comparison strains to determine whether there were fragments with similarity greater than the preset values. If the candidate specific regions met the above criteria, the candidate specific regions were considered the specific regions of the microbial target fragments.

### Searching for the multicopy regions

The target fragments of microorganisms may be chains or multiple incomplete motifs. When identifying multicopy regions in microbial target fragments, the motifs were connected together before searching for candidate multicopy regions, in which the microbial target fragments often have multiple incomplete motifs. These motifs are formed by incomplete splicing of short reads under existing second-generation sequencing conditions. To identify the multicopy regions in the microbial target fragments illustrated in Figure 3, 1) for searching candidate multicopy regions, internal alignments were performed on the microbial target fragments, and searching for the regions corresponding to the to-be-detected sequences for which the similarity met the preset values as candidate multicopy regions, and the similarity was the product of the coverage rates and matching rates of the to-be-detected sequence; 2) for verifying and obtaining the multicopy regions, the median values of the copy numbers of the candidate multicopy regions were obtained, including by a) determining the positions of each candidate multicopy region on the microorganism target fragments; b) obtaining the numbers of other candidate multicopy regions covering the positions of each base of the to-be-verified candidate multicopy regions; and c) calculating the median values of the copy numbers of the to-be-verified candidate multicopy regions.

The other candidate multicopy regions mentioned above refer to candidate multicopy regions other than the candidate multicopy regions to be verified. If the median copy numbers of the candidate multicopy regions were greater than 1, the candidate multicopy regions were recorded as multicopy regions. The preset value of the similarity could be determined as needed. The recommended preset value of similarity had to exceed 80%. If the region where the similarity met the preset value contained different motifs, the region was divided based on the original motif connection points and divided into different subregions to determine whether the subregions were candidate multicopy regions. The coverage rate = (length of similar sequence/(end value of the to-be-detected sequence – starting value of the to-be-detected sequence +1)) %. The matching rates referred to the identity values when the to-be-detected sequences were aligned with themselves. The identity values of the two aligned sequences could be obtained by software such as needle[48], water[49] or blat[50]. The length of similar sequences referred to the number of bases in which the matched fragments occupied the to-be-detected sequences when the to-be-detected sequence was aligned with other sequences, that is, the length of the matched fragments. In the preferred embodiment, the 95% confidence interval of the copy numbers of the candidate multicopy regions was calculated. The confidence interval refers to the estimated interval of the overall parameters constructed by the sample statistics, that is, the interval estimation of the overall copy numbers of the target regions. The confidence interval reflected the degree to which the true values of the copy numbers of the target regions were close to the measurement result, which indicates the credibility of the measured values of the measured parameters.

### Searching for the conserved regions

To obtain species-specific consensus sequences of microorganisms as presented in Figure 1, 1) for searching for candidate consensus sequences, specific sequences of target strains belonging to the same species were clustered based on the clustering algorithm[51] to obtain several candidate species-specific consensus sequences; and 2) for verifying and obtaining primary-screened species-specific consensus sequences, whether the candidate species-specific consensus sequences met the following conditions remapped by mafft was determined[52]. Herein, the strain coverage rates met the preset values, and the effective copy numbers met the preset values. If the candidate species-specific consensus sequences met all the above conditions, it was determined that the candidate species-specific consensus sequences were species-specific consensus sequences; the percentage of strain = (number of target strains with the candidate species-specific consensus sequence/total number of target strains) * 100%. The effective copy numbers, i.e., weighted average copy numbers were calculated according to formula (I), where n was the total number of copy number gradients of the candidate species-specific consensus sequences; Ci was the copy number corresponding to the i-th candidate species-specific consensus sequence; Si was the number of strains with the i-th candidate species-specific consensus sequence; and Sall was the total number of target strains. Formula (I) refers to the summation of Ci (Si/Sall), where i ranges from Cmin to Cmax, and the number of i is n. Cmin is the minimum copy number of all candidate species-specific consensus sequences. Cmax is the maximum copy number of all candidate species-specific consensus sequences.

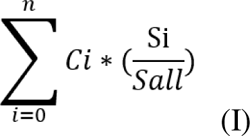

### Designing the best sets from the appropriate templates above

Based on the above various combinations of different submodules, the final candidate species-specific consensus sequences could be compared to the whole genomes of all target strains to calculate the percentage of strains and effective copy numbers of the candidate species-specific consensus sequences. Designing the templates of the primary-screened species-specific consensus sequence and achieving the best sets of primers and probes were performed as follows: 1) we obtained the candidate probes and primers by Primer3[53] or Beacon Designer™; 2) the sequences of the candidate probes and primers were aligned to the whole-genome sequences of all the target strains; 3) the strain coverage rates corresponding to the sequences of each probe and primer were calculated; and 4) the candidate probes and primers for which the strain coverage rates met the preset values were screened, and the primary-screened species-specific consensus sequences corresponding to the screened candidate probes and primers were chosen as the final species-specific consensus sequences.

## Conclusions

Overall, the Shine strategy was presented to identify specific, sensitive and conserved biomarkers from massive microbial genomic data within populations. We have proposed a design strategy to improve the quality of nucleic acid detection reagents, which has been validated by several clinical projects. Our method was highly accurate and sensitive and could detect newly discovered multicopy universal species-specific and even subspecies-specific target fragments, covering all identified epidemic pathogenic microorganisms. Therefore, the method was suitable for identifying shared universal phylogenetic biomarkers with few false positive or false negative results and automating the design of minimal primers and probes to detect pathogens with cost-effective predictive power.

## Declarations

### Ethics approval and consent to participate

All protocols were approved by the Liferiver Science and Technology Institute, Shanghai ZJ Bio-Tech Co., Ltd and Use Committee (Shanghai, China).

### Consent to publish

Not applicable.

### Availability of data and materials

All shared universal phylogenetic markers from massive microbiological genomics are available at the website https://bioinfo.liferiver.com.cn/#/home.

### Author information

Affiliations

### Liferiver Science and Technology Institute, Shanghai ZJ Bio-Tech Co., Ltd. Shanghai, China

Cong Ji, Junbin (Jack) Shao

Authors’ Contributions

The authors’ responsibilities were as follows: JBS and CJ designed and conducted the research; CJ and JBS analysed the data and performed the analysis; CJ wrote the paper; JBS and CJ revised the manuscript; and CJ had primary responsibility for the final content. The authors read and approved the final manuscript.

Corresponding author

Correspondence to Cong Ji and Junbin (Jack) Shao.

## Acknowledgments

Appreciation goes to Zhang Hanyan, Xiong Lei, and Pan Daxia for their experimental validation in this study. The authors deeply thank Liu Yan, Zhang Jie, and Li Qiang for their valuable suggestions and comments on this work. Many facets of the user interface design benefited from assistance provided by Niu Xingsheng, Lu Wang and Pan Yajie. We wish to express our thanks for the valuable modifications to the paper made by Wang Guangzhong, Shen Yilin, Zhu Lingjiao, Guo Jingjing, and Zhou Miaomiao, who helped us greatly revise this paper. All the other data supporting the findings of this study and the computational code used in this study are available from the corresponding authors upon reasonable request. Cong Ji and Junbin (Jack) Shao are named inventors on the pending PCT Patent Applications PCT/CN2020/090180, PCT/CN2020/090175, and PCT/CN2020/090177 filed by the Liferiver Science and Technology Institute of Shanghai ZJ Bio-Tech Co., Ltd., which separately describe the method and device for identifying multicopy, species-specific consensus sequences in microbial target fragments and use thereof. The other authors declare no competing interests.

## Abbreviations

PCR: Polymerase Chain Reaction

LDT: Laboratory-developed Test

DNA: Deoxyribo Nucleic Acid

RNA: Ribonucleic Acid

16S rRNA: 16S Ribosomal RNA

MEGA: Molecular Evolutionary Genetics Analysis

PAML: Phylogenetic Analysis by Maximum Likelihood

NCBI: National Center for Biotechnology Information

GISAID: Global Initiative on Sharing All Influenza Data

EzBioCloud: ChunLab’s public data and analytics portal

EuPathDB: Eukaryotic Pathogen Database Project

GiardiaDB: Giardia Genomics Resources

TrichDB: Trichomonas Informatics Resources

FungiDB: Fungal & Oomycete Informatics Resources

HKU1: Human coronavirus HKU1

OC43: Human coronavirus OC43

NL63: Human coronavirus NL63

229E: Human coronavirus 229E

MERS: Middle East respiratory syndrome-related coronavirus

SARS-CoV: Severe acute respiratory syndrome coronavirus

SARS-CoV-2: Severe acute respiratory syndrome coronavirus 2

MTBC: Mycobacterium tuberculosis complex

BP: Bordetella pertussis

BPP: Bordetella parapertussis

CAP: community-acquired pneumonia

HIV: human immunodeficiency virus

CRISPR: Clustered Regularly Interspaced Short Palindromic Repeats

Cas: CRISPR-associated (Cas) genes

SHERLOCK: specific high-sensitivity enzymatic reporter unlocking

STOP: SHERLOCK testing in one pot

SGs: study groups

## Figures, Tables and Supplementary Information

**Supplementary Table1. Cluster sets of identified undiscovered multicopy regions** from *Mycobacterium tuberculosis*, *Mycobacterium africanum*, *Mycobacterium bovis*, and the *Mycobacterium tuberculosis* complex.

**Supplementary Table2. Cluster sets of identified undiscovered multicopy regions from** *B. pertussis*, *B. parapertussis*, and *B. holmesii*.

**Supplementary Table3. Cluster sets of identified undiscovered multicopy regions from** *B. parapertussis* and *B. bronchiseptica*.

**Supplementary Table4. Cluster sets of identified undiscovered multicopy regions from** *M. pneumoniae* strain M129.

**Supplementary Table5. Cluster sets of identified undiscovered multicopy regions from** *Streptococcus agalactiae*.

**Supplementary Table6. Cluster sets of identified undiscovered multicopy regions from** *H. pylori* UA802, *H. pylori* strain PMSS1, and *H. pylori* strain 7.13.

**Supplementary Table7. Cluster sets of identified undiscovered multicopy regions from** *Legionella pneumophila*.

**Supplementary Table8. Cluster sets of identified undiscovered multicopy regions from** *Candida auris*.

