## Supplementary Tables for "Shine: A novel strategy to extract specific, sensitive and well-conserved biomarkers from massive microbial genomic datasets"

**Supplementary Table1.**

**Cluster sets of identified undiscovered multicopy regions from Mycobacterium tuberculosis, Mycobacterium Africanum, Mycobacterium bovis, Tuberculosis complex.**

| ClusterID | Distribution | Marker | Percent_of_strains | Weighted_Average_Copy |
|---|---|---|---|---|
| Cluster195 | 0(3)\|1(51)\|2(40)\|3(35)\|4(21)\|5(4)\|6(3)\|7(5)\|8(5)\|9(9)\|10(7)\|11(11)\|12(19)\|13(21)\|14(19)\|15(20)\|16(24)\|17(13)\|18(6)\|19(3)\|20(2)\|21(7)\|22(3)\|23(2)\|24(2)\|25(2)\|27(1)\|28(1)\| | * | 99.1%[336/339] | 8.831858407 |
| Cluster102 | 0(3)\|1(51)\|2(40)\|3(35)\|4(21)\|5(4)\|6(3)\|7(5)\|8(5)\|9(9)\|10(7)\|11(11)\|12(19)\|13(21)\|14(19)\|15(21)\|16(23)\|17(13)\|18(6)\|19(3)\|20(2)\|21(7)\|22(4)\|23(1)\|24(2)\|25(2)\|27(1)\|28(1)\| | * | 99.1%[336/339] | 8.825958702 |
| Cluster454 | 0(3)\|1(51)\|2(40)\|3(35)\|4(21)\|5(4)\|6(3)\|7(6)\|8(5)\|9(8)\|10(7)\|11(14)\|12(16)\|13(21)\|14(19)\|15(21)\|16(23)\|17(12)\|18(7)\|19(3)\|20(2)\|21(7)\|22(3)\|23(2)\|24(2)\|25(2)\|27(1)\|28(1)\| | * | 99.1%[336/339] | 8.817109145 |
| Cluster170 | 0(3)\|1(51)\|2(41)\|3(34)\|4(21)\|5(4)\|6(3)\|7(6)\|8(5)\|9(8)\|10(7)\|11(14)\|12(17)\|13(20)\|14(19)\|15(21)\|16(24)\|17(11)\|18(7)\|19(3)\|20(2)\|21(7)\|22(3)\|23(2)\|24(2)\|25(2)\|27(1)\|28(1)\| | * | 99.1%[336/339] | 8.808259587 |
| Cluster439 | 0(3)\|1(52)\|2(43)\|3(33)\|4(20)\|5(4)\|6(3)\|7(7)\|8(6)\|9(6)\|10(9)\|11(11)\|12(17)\|13(20)\|14(19)\|15(20)\|16(25)\|17(11)\|18(6)\|19(4)\|20(2)\|21(7)\|22(3)\|23(1)\|24(3)\|25(2)\|27(1)\|28(1)\| | * | 99.1%[336/339] | 8.764011799 |
| Cluster128 | 0(3)\|1(51)\|2(40)\|3(36)\|4(20)\|5(4)\|6(3)\|7(7)\|8(5)\|9(8)\|10(9)\|11(13)\|12(16)\|13(22)\|14(18)\|15(22)\|16(20)\|17(14)\|18(5)\|19(3)\|20(2)\|21(8)\|22(3)\|23(1)\|24(2)\|25(2)\|27(1)\|28(1)\| | * | 99.1%[336/339] | 8.749262537 |
| Cluster363 | 0(3)\|1(54)\|2(41)\|3(33)\|4(20)\|5(4)\|6(4)\|7(6)\|8(6)\|9(6)\|10(9)\|11(11)\|12(18)\|13(20)\|14(18)\|15(22)\|16(23)\|17(12)\|18(5)\|19(4)\|20(2)\|21(7)\|22(3)\|23(1)\|24(4)\|25(1)\|27(1)\|28(1)\| | * | 99.1%[336/339] | 8.737463127 |
| Cluster289 | 0(3)\|1(54)\|2(41)\|3(32)\|4(21)\|5(4)\|6(4)\|7(6)\|8(6)\|9(6)\|10(9)\|11(12)\|12(16)\|13(22)\|14(17)\|15(22)\|16(23)\|17(12)\|18(5)\|19(4)\|20(2)\|21(7)\|22(3)\|23(2)\|24(3)\|25(1)\|27(1)\|28(1)\| | * | 99.1%[336/339] | 8.734513274 |
| Cluster76 | 0(3)\|1(52)\|2(43)\|3(33)\|4(20)\|5(4)\|6(3)\|7(7)\|8(6)\|9(6)\|10(10)\|11(11)\|12(17)\|13(21)\|14(19)\|15(23)\|16(20)\|17(12)\|18(6)\|19(3)\|20(2)\|21(7)\|22(4)\|23(1)\|24(3)\|25(1)\|27(1)\|28(1)\| | * | 99.1%[336/339] | 8.713864307 |
| Cluster599 | 0(3)\|1(58)\|2(42)\|3(32)\|4(22)\|5(4)\|6(3)\|7(8)\|8(3)\|9(7)\|10(8)\|11(11)\|12(14)\|13(19)\|14(17)\|15(23)\|16(24)\|17(11)\|18(5)\|19(4)\|20(2)\|21(8)\|22(2)\|23(1)\|24(4)\|25(1)\|26(1)\|27(1)\|28(1)\| | * | 99.1%[336/339] | 8.613569322 |

| | | | | |
|---|---|---|---|---|
| Cluster532 | 0(3)\|1(58)\|2(49)\|3(22)\|4(25)\|5(7)\|6(1)\|7(5)\|8(3)\|9(14)\|10(5)\|11(11)\|12(16)\|13(17)\|14(14)\|15(24)\|16(17)\|17(13)\|18(9)\|19(4)\|20(1)\|21(9)\|22(3)\|23(2)\|24(3)\|25(3)\|26(1)\| | * | 99.1%[336/339] | 8.592920354 |
| Cluster160 | 0(3)\|1(58)\|2(49)\|3(22)\|4(26)\|5(6)\|6(1)\|7(5)\|8(4)\|9(13)\|10(5)\|11(11)\|12(16)\|13(17)\|14(15)\|15(23)\|16(17)\|17(14)\|18(8)\|19(4)\|20(1)\|21(10)\|22(2)\|23(2)\|24(3)\|25(3)\|26(1)\| | * | 99.1%[336/339] | 8.578171091 |
| Cluster606 | 0(3)\|1(58)\|2(49)\|3(22)\|4(26)\|5(6)\|6(1)\|7(5)\|8(4)\|9(13)\|10(5)\|11(11)\|12(16)\|13(17)\|14(14)\|15(24)\|16(17)\|17(15)\|18(8)\|19(3)\|20(2)\|21(9)\|22(2)\|23(2)\|24(3)\|25(3)\|26(1)\| | * | 99.1%[336/339] | 8.572271386 |
| Cluster537 | 0(3)\|1(58)\|2(49)\|3(22)\|4(27)\|5(5)\|6(1)\|7(5)\|8(3)\|9(14)\|10(6)\|11(11)\|12(16)\|13(18)\|14(13)\|15(23)\|16(17)\|17(13)\|18(9)\|19(4)\|20(1)\|21(9)\|22(3)\|23(2)\|24(3)\|25(3)\|26(1)\| | * | 99.1%[336/339] | 8.569321534 |
| Cluster101 | 0(3)\|1(58)\|2(49)\|3(22)\|4(27)\|5(5)\|6(1)\|7(5)\|8(4)\|9(13)\|10(6)\|11(11)\|12(16)\|13(18)\|14(13)\|15(23)\|16(17)\|17(13)\|18(9)\|19(4)\|20(1)\|21(9)\|22(3)\|23(2)\|24(3)\|25(3)\|26(1)\| | * | 99.1%[336/339] | 8.566371681 |
| Cluster615 | 0(4)\|1(57)\|2(49)\|3(23)\|4(25)\|5(6)\|6(2)\|7(5)\|8(5)\|9(12)\|10(5)\|11(12)\|12(14)\|13(17)\|14(14)\|15(24)\|16(17)\|17(15)\|18(8)\|19(3)\|20(2)\|21(9)\|22(2)\|23(3)\|24(3)\|25(2)\|26(1)\| | * | 98.8%[335/339] | 8.536873156 |
| Cluster2 | 0(4)\|1(57)\|2(42)\|3(33)\|4(21)\|5(4)\|6(3)\|7(9)\|8(3)\|9(8)\|10(6)\|11(11)\|12(15)\|13(19)\|14(18)\|15(25)\|16(21)\|17(12)\|18(5)\|19(3)\|20(3)\|21(7)\|22(3)\|23(1)\|24(3)\|25(1)\|26(1)\|28(1)\| | * | 98.8%[335/339] | 8.525073746 |
| Cluster535 | 0(3)\|1(59)\|2(47)\|3(31)\|4(22)\|5(4)\|6(2)\|7(9)\|8(2)\|9(9)\|10(6)\|11(11)\|12(11)\|13(20)\|14(16)\|15(23)\|16(20)\|17(14)\|18(5)\|19(4)\|20(3)\|21(6)\|22(4)\|23(1)\|24(3)\|25(1)\|26(2)\|28(1)\| | * | 99.1%[336/339] | 8.454277286 |
| Cluster397 | 0(3)\|1(59)\|2(47)\|3(31)\|4(22)\|5(4)\|6(2)\|7(9)\|8(2)\|9(9)\|10(6)\|11(11)\|12(11)\|13(20)\|14(16)\|15(23)\|16(20)\|17(14)\|18(5)\|19(4)\|20(3)\|21(6)\|22(4)\|23(1)\|24(3)\|25(1)\|26(2)\|28(1)\| | * | 99.1%[336/339] | 8.454277286 |
| Cluster396 | 0(3)\|1(59)\|2(47)\|3(31)\|4(22)\|5(4)\|6(2)\|7(9)\|8(2)\|9(9)\|10(6)\|11(11)\|12(11)\|13(20)\|14(16)\|15(23)\|16(20)\|17(14)\|18(5)\|19(4)\|20(3)\|21(6)\|22(4)\|23(1)\|24(3)\|25(1)\|26(2)\|28(1)\| | * | 99.1%[336/339] | 8.454277286 |
| Cluster182 | 0(3)\|1(60)\|2(46)\|3(32)\|4(21)\|5(4)\|6(2)\|7(9)\|8(2)\|9(11)\|10(5)\|11(11)\|12(10)\|13(20)\|14(17)\|15(22)\|16(23)\|17(11)\|18(5)\|19(4)\|20(3)\|21(6)\|22(4)\|23(1)\|24(3)\|25(1)\|26(2)\|28(1)\| | * | 99.1%[336/339] | 8.424778761 |
| Cluster605 | 0(2)\|1(65)\|2(45)\|3(28)\|4(24)\|5(5)\|6(1)\|7(5)\|8(4)\|9(15)\|10(4)\|11(6)\|12(13)\|13(19)\|14(16)\|15(23)\|16(19)\|17(14)\|18(6)\|19(4)\|20(3)\|21(6)\|22(3)\|23(2)\|24(3)\|25(1)\|26(3)\| | * | 99.4%[337/339] | 8.392330383 |
| Cluster290 | 0(4)\|1(57)\|2(44)\|3(35)\|4(21)\|5(2)\|6(6)\|7(6)\|8(4)\|9(8)\|10(7)\|11(10)\|12(14)\|13(21)\|14(17)\|15(26)\|16(18)\|17(12)\|18(5)\|19(2)\|20(4)\|21(6)\|22(4)\|23(1)\|24(2)\|25(1)\|26(1)\|28(1)\| | * | 98.8%[335/339] | 8.374631268 |

| | | | | |
|---|---|---|---|---|
| Cluster246 | 0(7)\|1(63)\|2(43)\|3(23)\|4(25)\|5(4)\|6(3)\|7(6)\|8(5)\|9(11)\|10(7)\|11(11)\|12(14)\|13(17)\|14(14)\|15(23)\|16(17)\|17(13)\|18(8)\|19(4)\|20(1)\|21(9)\|22(2)\|23(3)\|24(4)\|25(2)\| | * | 97.9%[332/339] | 8.374631268 |
| Cluster315 | 0(4)\|1(64)\|2(47)\|3(28)\|4(24)\|5(3)\|6(2)\|7(4)\|8(7)\|9(11)\|10(5)\|11(8)\|12(12)\|13(16)\|14(15)\|15(23)\|16(16)\|17(20)\|18(5)\|19(3)\|21(10)\|22(2)\|23(3)\|24(3)\|25(2)\|26(2)\| | * | 98.8%[335/339] | 8.356932153 |
| Cluster67 | 0(3)\|1(66)\|2(46)\|3(27)\|4(24)\|5(4)\|6(1)\|7(6)\|8(4)\|9(13)\|10(5)\|11(6)\|12(14)\|13(18)\|14(15)\|15(23)\|16(18)\|17(16)\|18(5)\|19(3)\|20(2)\|21(8)\|22(2)\|23(3)\|24(3)\|25(2)\|26(2)\| | * | 99.1%[336/339] | 8.339233038 |
| Cluster395 | 0(3)\|1(66)\|2(46)\|3(27)\|4(24)\|5(4)\|6(1)\|7(6)\|8(4)\|9(13)\|10(5)\|11(6)\|12(14)\|13(18)\|14(15)\|15(23)\|16(18)\|17(16)\|18(5)\|19(3)\|20(2)\|21(8)\|22(2)\|23(3)\|24(3)\|25(2)\|26(2)\| | * | 99.1%[336/339] | 8.339233038 |
| Cluster317 | 0(3)\|1(66)\|2(47)\|3(27)\|4(24)\|5(3)\|6(2)\|7(5)\|8(6)\|9(12)\|10(6)\|11(7)\|12(11)\|13(17)\|14(14)\|15(25)\|16(15)\|17(19)\|18(5)\|19(3)\|21(10)\|22(2)\|23(3)\|24(3)\|25(2)\|26(2)\| | * | 99.1%[336/339] | 8.327433628 |
| Cluster212 | 0(5)\|1(62)\|2(45)\|3(28)\|4(24)\|5(5)\|6(1)\|7(5)\|8(4)\|9(15)\|10(4)\|11(6)\|12(14)\|13(18)\|14(16)\|15(25)\|16(17)\|17(16)\|18(6)\|19(4)\|20(6)\|21(2)\|22(3)\|23(1)\|24(4)\|25(3)\| | * | 98.5%[334/339] | 8.324483776 |
| Cluster514 | 0(3)\|1(66)\|2(47)\|3(27)\|4(23)\|5(4)\|6(2)\|7(5)\|8(6)\|9(12)\|10(5)\|11(6)\|12(13)\|13(18)\|14(15)\|15(23)\|16(18)\|17(16)\|18(5)\|19(3)\|20(1)\|21(9)\|22(2)\|23(3)\|24(3)\|25(2)\|26(2)\| | * | 99.1%[336/339] | 8.318584071 |
| Cluster88 | 0(5)\|1(62)\|2(46)\|3(27)\|4(24)\|5(5)\|6(2)\|7(5)\|8(4)\|9(14)\|10(4)\|11(7)\|12(13)\|13(17)\|14(16)\|15(26)\|16(17)\|17(16)\|18(6)\|19(4)\|20(6)\|21(2)\|22(3)\|23(1)\|24(4)\|25(3)\| | * | 98.5%[334/339] | 8.315634218 |
| Cluster667 | 0(3)\|1(66)\|2(47)\|3(27)\|4(24)\|5(3)\|6(2)\|7(5)\|8(6)\|9(12)\|10(5)\|11(7)\|12(12)\|13(17)\|14(16)\|15(23)\|16(18)\|17(16)\|18(5)\|19(3)\|20(1)\|21(9)\|22(2)\|23(3)\|24(3)\|25(2)\|26(2)\| | * | 99.1%[336/339] | 8.315634218 |
| Cluster581 | 0(5)\|1(67)\|2(46)\|3(27)\|4(25)\|5(3)\|6(1)\|7(4)\|8(8)\|9(10)\|10(5)\|11(8)\|12(12)\|13(14)\|14(15)\|15(23)\|16(17)\|17(19)\|18(5)\|19(3)\|21(11)\|22(1)\|23(3)\|24(3)\|25(2)\|26(2)\| | * | 98.5%[334/339] | 8.259587021 |
| Cluster556 | 0(4)\|1(64)\|2(45)\|3(33)\|4(19)\|5(3)\|6(6)\|7(5)\|8(4)\|9(13)\|10(7)\|11(8)\|12(10)\|13(20)\|14(16)\|15(25)\|16(19)\|17(11)\|18(6)\|19(1)\|20(5)\|21(6)\|22(3)\|23(1)\|24(2)\|25(1)\|26(1)\|27(1)\| | * | 98.8%[335/339] | 8.14159292 |
| Cluster84 | 0(3)\|1(7)\|2(2)\|3(25)\|4(3)\|5(20)\|6(263)\|7(15)\|9(1)\| | * | 99.1%[336/339] | 5.575221239 |
| Cluster83 | 0(3)\|1(7)\|2(2)\|3(25)\|4(3)\|5(20)\|6(263)\|7(15)\|9(1)\| | * | 99.1%[336/339] | 5.575221239 |
| Cluster82 | 0(3)\|1(7)\|2(2)\|3(25)\|4(3)\|5(20)\|6(263)\|7(15)\|9(1)\| | * | 99.1%[336/339] | 5.575221239 |
| Cluster294 | 0(3)\|1(7)\|2(2)\|3(25)\|4(3)\|5(20)\|6(263)\|7(15)\|9(1)\| | * | 99.1%[336/339] | 5.575221239 |

| | | | | |
|---|---|---|---|---|
| Cluster460 | 0(19)\|1(23)\|2(36)\|3(31)\|4(13)\|5(37)\|6(159)\|7(8)\|8(1)\|9(1)\|10(1)\|11(10)\| | * | 94.4%[320/339] | 4.637168142 |
| Cluster410 | 0(1)\|1(2)\|2(6)\|3(28)\|4(65)\|5(236)\|7(1)\| | * | 99.7%[338/339] | 4.557522124 |
| Cluster459 | 0(2)\|1(6)\|2(27)\|3(290)\|4(14)\| | * | 99.4%[337/339] | 2.908554572 |
| Cluster523 | 0(2)\|1(8)\|2(39)\|3(272)\|4(18)\| | * | 99.4%[337/339] | 2.873156342 |
| Cluster652 | 0(2)\|1(8)\|2(40)\|3(272)\|4(17)\| | * | 99.4%[337/339] | 2.867256637 |
| Cluster586 | 0(1)\|1(8)\|2(40)\|3(283)\|4(6)\|5(1)\| | * | 99.7%[338/339] | 2.849557522 |
| Cluster634 | 0(2)\|1(8)\|2(51)\|3(261)\|4(17)\| | * | 99.4%[337/339] | 2.83480826 |
| Cluster16 | 1(7)\|2(45)\|3(286)\|4(1)\| | * | 100.0%[339/339] | 2.828908555 |
| Cluster187 | 0(2)\|1(9)\|2(53)\|3(262)\|4(13)\| | * | 99.4%[337/339] | 2.81120944 |
| Cluster245 | 0(2)\|1(39)\|2(69)\|3(178)\|4(39)\|5(12)\| | * | 99.4%[337/339] | 2.734513274 |
| Cluster372 | 0(8)\|1(15)\|2(62)\|3(241)\|4(9)\|5(2)\|6(2)\| | * | 97.6%[331/339] | 2.713864307 |
| Cluster474 | 0(4)\|1(21)\|2(60)\|3(239)\|4(14)\|5(1)\| | * | 98.8%[335/339] | 2.710914454 |
| Cluster604 | 0(3)\|1(25)\|2(101)\|3(156)\|4(54)\| | * | 99.1%[336/339] | 2.687315634 |
| Cluster339 | 0(5)\|1(39)\|2(81)\|3(179)\|4(23)\|5(11)\|6(1)\| | * | 98.5%[334/339] | 2.628318584 |
| Cluster510 | 0(5)\|1(41)\|2(83)\|3(178)\|4(19)\|5(12)\|6(1)\| | * | 98.5%[334/339] | 2.604719764 |
| Cluster117 | 0(1)\|1(20)\|2(97)\|3(220)\|4(1)\| | * | 99.7%[338/339] | 2.589970501 |
| Cluster295 | 0(13)\|1(51)\|2(76)\|3(177)\|4(11)\|5(10)\|6(1)\| | * | 96.2%[326/339] | 2.460176991 |
| Cluster32 | 0(11)\|1(58)\|2(81)\|3(166)\|4(13)\|5(9)\|6(1)\| | * | 96.8%[328/339] | 2.421828909 |
| Cluster98 | 0(4)\|1(25)\|2(224)\|3(3)\|4(83)\| | * | 98.8%[335/339] | 2.401179941 |
| Cluster56 | 0(10)\|1(60)\|2(86)\|3(163)\|4(10)\|5(9)\|6(1)\| | * | 97.1%[329/339] | 2.395280236 |
| Cluster139 | 0(4)\|1(31)\|2(217)\|3(4)\|4(83)\| | * | 98.8%[335/339] | 2.386430678 |
| Cluster248 | 0(4)\|1(31)\|2(218)\|3(3)\|4(83)\| | * | 98.8%[335/339] | 2.383480826 |
| Cluster258 | 0(1)\|1(29)\|2(225)\|3(24)\|4(60)\| | * | 99.7%[338/339] | 2.333333333 |
| Cluster526 | 0(1)\|1(22)\|2(233)\|3(35)\|4(48)\| | * | 99.7%[338/339] | 2.315634218 |
| Cluster354 | 0(1)\|1(22)\|2(233)\|3(35)\|4(48)\| | * | 99.7%[338/339] | 2.315634218 |

| | | | | |
|---|---|---|---|---|
| Cluster564 | 0(2)\|1(35)\|2(221)\|3(21)\|4(60)\| | * | 99.4%[337/339] | 2.300884956 |
| Cluster654 | 0(1)\|1(15)\|2(255)\|3(18)\|4(50)\| | * | 99.7%[338/339] | 2.297935103 |
| Cluster231 | 1(20)\|2(255)\|3(13)\|4(51)\| | * | 100.0%[339/339] | 2.280235988 |
| Cluster49 | 1(20)\|2(258)\|3(10)\|4(51)\| | * | 100.0%[339/339] | 2.271386431 |
| Cluster573 | 0(2)\|1(42)\|2(158)\|3(137)\| | * | 99.4%[337/339] | 2.268436578 |
| Cluster681 | 0(1)\|1(25)\|2(246)\|3(17)\|4(50)\| | * | 99.7%[338/339] | 2.265486726 |
| Cluster651 | 0(2)\|1(60)\|2(196)\|3(8)\|4(73)\| | * | 99.4%[337/339] | 2.265486726 |
| Cluster255 | 1(24)\|2(251)\|3(14)\|4(50)\| | * | 100.0%[339/339] | 2.265486726 |
| Cluster87 | 0(13)\|1(29)\|2(216)\|3(21)\|4(60)\| | * | 96.2%[326/339] | 2.253687316 |
| Cluster105 | 0(13)\|1(31)\|2(214)\|3(21)\|4(60)\| | * | 96.2%[326/339] | 2.247787611 |
| Cluster261 | 0(2)\|1(30)\|2(243)\|3(14)\|4(50)\| | * | 99.4%[337/339] | 2.235988201 |
| Cluster80 | 1(32)\|2(249)\|3(7)\|4(51)\| | * | 100.0%[339/339] | 2.227138643 |
| Cluster21 | 1(31)\|2(251)\|3(6)\|4(51)\| | * | 100.0%[339/339] | 2.227138643 |
| Cluster17 | 0(5)\|1(28)\|2(242)\|3(14)\|4(50)\| | * | 98.5%[334/339] | 2.224188791 |
| Cluster1 | 0(3)\|1(32)\|2(240)\|3(14)\|4(50)\| | * | 99.1%[336/339] | 2.224188791 |
| Cluster125 | 0(2)\|1(31)\|2(247)\|3(8)\|4(51)\| | * | 99.4%[337/339] | 2.221238938 |
| Cluster59 | 0(3)\|1(31)\|2(245)\|3(9)\|4(51)\| | * | 99.1%[336/339] | 2.218289086 |
| Cluster345 | 0(3)\|1(31)\|2(245)\|3(9)\|4(51)\| | * | 99.1%[336/339] | 2.218289086 |
| Cluster650 | 0(4)\|1(33)\|2(238)\|3(14)\|4(50)\| | * | 98.8%[335/339] | 2.215339233 |
| Cluster557 | 0(3)\|1(31)\|2(247)\|3(7)\|4(51)\| | * | 99.1%[336/339] | 2.212389381 |
| Cluster547 | 0(5)\|1(32)\|2(238)\|3(14)\|4(50)\| | * | 98.5%[334/339] | 2.212389381 |
| Cluster269 | 0(5)\|1(32)\|2(238)\|3(14)\|4(50)\| | * | 98.5%[334/339] | 2.212389381 |
| Cluster570 | 0(8)\|1(27)\|2(241)\|3(13)\|4(50)\| | * | 97.6%[331/339] | 2.206489676 |
| Cluster583 | 0(1)\|1(32)\|2(253)\|3(3)\|4(50)\| | * | 99.7%[338/339] | 2.203539823 |
| Cluster453 | 0(23)\|1(29)\|2(206)\|3(23)\|4(58)\| | * | 93.2%[316/339] | 2.18879056 |

| | | | | |
|---|---|---|---|---|
| Cluster50 | 0(11)\|1(28)\|2(237)\|3(13)\|4(50)\| | * | 96.8%[328/339] | 2.185840708 |
| Cluster292 | 0(11)\|1(28)\|2(237)\|3(13)\|4(50)\| | * | 96.8%[328/339] | 2.185840708 |
| Cluster70 | 0(3)\|1(38)\|2(246)\|3(2)\|4(50)\| | * | 99.1%[336/339] | 2.171091445 |
| Cluster657 | 0(24)\|1(29)\|2(227)\|3(8)\|4(51)\| | * | 92.9%[315/339] | 2.097345133 |
| Cluster633 | 0(3)\|1(61)\|2(183)\|3(90)\|4(2)\| | * | 99.1%[336/339] | 2.079646018 |
| Cluster366 | 0(29)\|1(25)\|2(226)\|3(8)\|4(51)\| | * | 91.4%[310/339] | 2.079646018 |
| Cluster356 | 0(2)\|1(71)\|2(172)\|3(87)\|4(7)\| | * | 99.4%[337/339] | 2.076696165 |
| Cluster399 | 0(1)\|1(130)\|2(53)\|3(154)\|4(1)\| | * | 99.7%[338/339] | 2.07079646 |
| Cluster48 | 0(32)\|1(25)\|2(223)\|3(8)\|4(51)\| | * | 90.6%[307/339] | 2.061946903 |
| Cluster447 | 0(32)\|1(25)\|2(223)\|3(8)\|4(51)\| | * | 90.6%[307/339] | 2.061946903 |
| Cluster282 | 0(32)\|1(25)\|2(223)\|3(8)\|4(51)\| | * | 90.6%[307/339] | 2.061946903 |
| Cluster179 | 0(32)\|1(26)\|2(222)\|3(8)\|4(51)\| | * | 90.6%[307/339] | 2.05899705 |
| Cluster361 | 0(4)\|1(63)\|2(184)\|3(86)\|4(2)\| | * | 98.8%[335/339] | 2.056047198 |
| Cluster408 | 1(45)\|2(248)\|3(30)\|4(16)\| | * | 100.0%[339/339] | 2.050147493 |
| Cluster458 | 0(3)\|1(42)\|2(247)\|3(31)\|4(16)\| | * | 99.1%[336/339] | 2.044247788 |
| Cluster52 | 0(19)\|1(44)\|2(228)\|3(2)\|4(46)\| | * | 94.4%[320/339] | 2.03539823 |
| Cluster608 | 0(1)\|1(49)\|2(244)\|3(29)\|4(16)\| | * | 99.7%[338/339] | 2.029498525 |
| Cluster436 | 0(1)\|1(50)\|2(243)\|3(29)\|4(16)\| | * | 99.7%[338/339] | 2.026548673 |
| Cluster369 | 0(10)\|1(71)\|2(165)\|3(86)\|4(7)\| | * | 97.1%[329/339] | 2.026548673 |
| Cluster402 | 0(3)\|1(47)\|2(245)\|3(28)\|4(16)\| | * | 99.1%[336/339] | 2.020648968 |
| Cluster73 | 0(2)\|1(54)\|2(239)\|3(28)\|4(16)\| | * | 99.4%[337/339] | 2.005899705 |
| Cluster69 | 1(4)\|2(332)\|3(3)\| | * | 100.0%[339/339] | 1.997050147 |
| Cluster466 | 1(4)\|2(332)\|3(3)\| | * | 100.0%[339/339] | 1.997050147 |
| Cluster528 | 0(1)\|1(1)\|2(336)\|3(1)\| | * | 99.7%[338/339] | 1.994100295 |
| Cluster613 | 0(6)\|1(76)\|2(173)\|3(83)\|4(1)\| | * | 98.2%[333/339] | 1.991150442 |

| | | | | |
|---|---|---|---|---|
| Cluster94 | 0(1)\|1(139)\|2(70)\|3(124)\|4(4)\|5(1)\| | * | 99.7%[338/339] | 1.982300885 |
| Cluster448 | 0(10)\|1(77)\|2(163)\|3(88)\|4(1)\| | * | 97.1%[329/339] | 1.979351032 |
| Cluster120 | 0(2)\|1(8)\|2(328)\|4(1)\| | * | 99.4%[337/339] | 1.970501475 |
| Cluster565 | 0(10)\|1(75)\|2(171)\|3(82)\|4(1)\| | * | 97.1%[329/339] | 1.967551622 |
| Cluster401 | 0(2)\|1(8)\|2(329)\| | * | 99.4%[337/339] | 1.96460177 |
| Cluster631 | 0(3)\|1(8)\|2(327)\|3(1)\| | * | 99.1%[336/339] | 1.961651917 |
| Cluster230 | 0(4)\|1(15)\|2(316)\|3(4)\| | * | 98.8%[335/339] | 1.943952802 |
| Cluster412 | 0(1)\|1(20)\|2(316)\|3(2)\| | * | 99.7%[338/339] | 1.94100295 |
| Cluster538 | 0(2)\|1(133)\|2(91)\|3(110)\|4(3)\| | * | 99.4%[337/339] | 1.938053097 |
| Cluster374 | 1(24)\|2(315)\| | * | 100.0%[339/339] | 1.92920354 |
| Cluster334 | 0(4)\|1(17)\|2(317)\|3(1)\| | * | 98.8%[335/339] | 1.92920354 |
| Cluster497 | 0(1)\|1(23)\|2(315)\| | * | 99.7%[338/339] | 1.926253687 |
| Cluster664 | 0(1)\|1(24)\|2(314)\| | * | 99.7%[338/339] | 1.923303835 |
| Cluster39 | 0(2)\|1(141)\|2(86)\|3(105)\|4(5)\| | * | 99.4%[337/339] | 1.911504425 |
| Cluster267 | 0(8)\|1(15)\|2(315)\|3(1)\| | * | 97.6%[331/339] | 1.911504425 |
| Cluster656 | 0(4)\|1(30)\|2(301)\|3(4)\| | * | 98.8%[335/339] | 1.899705015 |
| Cluster542 | 0(9)\|1(32)\|2(285)\|3(12)\|4(1)\| | * | 97.3%[330/339] | 1.89380531 |
| Cluster227 | 0(9)\|1(32)\|2(284)\|3(14)\| | * | 97.3%[330/339] | 1.89380531 |
| Cluster472 | 0(6)\|1(41)\|2(281)\|3(10)\|4(1)\| | * | 98.2%[333/339] | 1.879056047 |
| Cluster367 | 0(11)\|1(21)\|2(305)\|3(2)\| | * | 96.8%[328/339] | 1.879056047 |
| Cluster569 | 0(3)\|1(108)\|2(186)\|3(12)\|4(30)\| | * | 99.1%[336/339] | 1.876106195 |
| Cluster333 | 0(3)\|1(108)\|2(186)\|3(12)\|4(30)\| | * | 99.1%[336/339] | 1.876106195 |
| Cluster679 | 0(7)\|1(38)\|2(286)\|3(7)\|4(1)\| | * | 97.9%[332/339] | 1.873156342 |
| Cluster207 | 0(7)\|1(42)\|2(278)\|3(12)\| | * | 97.9%[332/339] | 1.87020649 |
| Cluster232 | 0(6)\|1(35)\|2(296)\|3(2)\| | * | 98.2%[333/339] | 1.867256637 |

| | | | | |
|---|---|---|---|---|
| Cluster443 | 0(6)\|1(44)\|2(280)\|3(8)\|4(1)\| | * | 98.2%[333/339] | 1.864306785 |
| Cluster166 | 0(3)\|1(111)\|2(184)\|3(11)\|4(30)\| | * | 99.1%[336/339] | 1.864306785 |
| Cluster438 | 0(30)\|1(100)\|2(110)\|3(85)\|4(14)\| | * | 91.2%[309/339] | 1.861356932 |
| Cluster389 | 0(3)\|1(56)\|2(267)\|3(12)\|4(1)\| | * | 99.1%[336/339] | 1.85840708 |
| Cluster568 | 0(14)\|1(49)\|2(261)\|3(2)\|4(13)\| | * | 95.9%[325/339] | 1.855457227 |
| Cluster165 | 0(4)\|1(112)\|2(183)\|3(10)\|4(30)\| | * | 98.8%[335/339] | 1.852507375 |
| Cluster159 | 0(4)\|1(52)\|2(275)\|3(7)\|4(1)\| | * | 98.8%[335/339] | 1.849557522 |
| Cluster321 | 0(8)\|1(41)\|2(288)\|3(2)\| | * | 97.6%[331/339] | 1.837758112 |
| Cluster663 | 0(5)\|1(113)\|2(182)\|3(11)\|4(28)\| | * | 98.5%[334/339] | 1.83480826 |
| Cluster461 | 0(5)\|1(112)\|2(184)\|3(10)\|4(28)\| | * | 98.5%[334/339] | 1.83480826 |
| Cluster673 | 0(5)\|1(63)\|2(260)\|3(11)\| | * | 98.5%[334/339] | 1.817109145 |
| Cluster414 | 0(7)\|1(49)\|2(283)\| | * | 97.9%[332/339] | 1.814159292 |
| Cluster670 | 0(8)\|1(61)\|2(257)\|3(13)\| | * | 97.6%[331/339] | 1.81120944 |
| Cluster624 | 0(10)\|1(59)\|2(256)\|3(14)\| | * | 97.1%[329/339] | 1.808259587 |
| Cluster585 | 0(11)\|1(56)\|2(260)\|3(12)\| | * | 96.8%[328/339] | 1.805309735 |
| Cluster578 | 0(7)\|1(120)\|2(175)\|3(9)\|4(28)\| | * | 97.9%[332/339] | 1.796460177 |
| Cluster590 | 0(14)\|1(56)\|2(257)\|3(12)\| | * | 95.9%[325/339] | 1.787610619 |
| Cluster609 | 0(7)\|1(89)\|2(216)\|3(27)\| | * | 97.9%[332/339] | 1.775811209 |
| Cluster601 | 0(13)\|1(63)\|2(252)\|3(11)\| | * | 96.2%[326/339] | 1.769911504 |
| Cluster299 | 0(7)\|1(91)\|2(215)\|3(26)\| | * | 97.9%[332/339] | 1.766961652 |
| Cluster118 | 0(11)\|1(62)\|2(264)\|3(2)\| | * | 96.8%[328/339] | 1.758112094 |
| Cluster473 | 0(3)\|1(94)\|2(232)\|3(8)\|4(2)\| | * | 99.1%[336/339] | 1.740412979 |
| Cluster218 | 0(31)\|1(109)\|2(123)\|3(70)\|4(5)\|5(1)\| | * | 90.9%[308/339] | 1.740412979 |
| Cluster303 | 0(3)\|1(95)\|2(231)\|3(8)\|4(2)\| | * | 99.1%[336/339] | 1.737463127 |
| Cluster384 | 1(93)\|2(245)\|3(1)\| | * | 100.0%[339/339] | 1.728613569 |

| | | | | |
|---|---|---|---|---|
| Cluster268 | 0(21)\|1(110)\|2(156)\|3(45)\|4(6)\|5(1)\| | * | 93.8%[318/339] | 1.728613569 |
| Cluster558 | 0(41)\|1(30)\|2(256)\|3(12)\| | * | 87.9%[298/339] | 1.705014749 |
| Cluster119 | 0(40)\|1(31)\|2(257)\|3(11)\| | * | 88.2%[299/339] | 1.705014749 |
| Cluster632 | 0(16)\|1(82)\|2(240)\|3(1)\| | * | 95.3%[323/339] | 1.666666667 |
| Cluster446 | 0(10)\|1(129)\|2(186)\|3(3)\|4(11)\| | * | 97.1%[329/339] | 1.634218289 |
| Cluster40 | 0(13)\|1(131)\|2(182)\|3(2)\|4(11)\| | * | 96.2%[326/339] | 1.607669617 |
| Cluster91 | 0(34)\|1(102)\|2(175)\|3(27)\|4(1)\| | * | 90.0%[305/339] | 1.584070796 |
| Cluster359 | 0(14)\|1(117)\|2(208)\| | * | 95.9%[325/339] | 1.572271386 |
| Cluster310 | 0(9)\|1(231)\|2(13)\|3(79)\|4(3)\|5(4)\| | * | 97.3%[330/339] | 1.551622419 |
| Cluster44 | 0(23)\|1(137)\|2(151)\|3(27)\|4(1)\| | * | 93.2%[316/339] | 1.545722714 |
| Cluster320 | 0(14)\|1(139)\|2(182)\|3(2)\|4(2)\| | * | 95.9%[325/339] | 1.525073746 |
| Cluster416 | 0(13)\|1(138)\|2(187)\|3(1)\| | * | 96.2%[326/339] | 1.519174041 |
| Cluster636 | 0(30)\|1(142)\|2(139)\|3(27)\|4(1)\| | * | 91.2%[309/339] | 1.489675516 |
| Cluster612 | 0(4)\|1(189)\|2(133)\|3(2)\|4(11)\| | * | 98.8%[335/339] | 1.489675516 |
| Cluster431 | 0(3)\|1(171)\|2(163)\|3(2)\| | * | 99.1%[336/339] | 1.483775811 |
| Cluster440 | 0(50)\|1(103)\|2(160)\|3(25)\|4(1)\| | * | 85.3%[289/339] | 1.480825959 |
| Cluster683 | 0(36)\|1(133)\|2(143)\|3(26)\|4(1)\| | * | 89.4%[303/339] | 1.477876106 |
| Cluster592 | 0(5)\|1(183)\|2(145)\|3(4)\|4(2)\| | * | 98.5%[334/339] | 1.454277286 |
| Cluster72 | 0(5)\|1(191)\|2(137)\|3(4)\|4(2)\| | * | 98.5%[334/339] | 1.430678466 |
| Cluster539 | 0(42)\|1(137)\|2(152)\|3(8)\| | * | 87.6%[297/339] | 1.371681416 |
| Cluster296 | 0(12)\|1(194)\|2(129)\|3(3)\|4(1)\| | * | 96.5%[327/339] | 1.371681416 |
| Cluster193 | 0(20)\|1(196)\|2(121)\|3(1)\|4(1)\| | * | 94.1%[319/339] | 1.312684366 |
| Cluster403 | 0(12)\|1(216)\|2(108)\|3(3)\| | * | 96.5%[327/339] | 1.300884956 |
| Cluster36 | 0(6)\|1(232)\|2(97)\|3(4)\| | * | 98.2%[333/339] | 1.292035398 |
| Cluster192 | 0(20)\|1(210)\|2(105)\|3(4)\| | * | 94.1%[319/339] | 1.274336283 |

| | | | | |
|---|---|---|---|---|
| Cluster279 | 0(36)\|1(216)\|2(52)\|3(31)\|4(3)\|5(1)\| | * | 89.4%[303/339] | 1.268436578 |
| Cluster580 | 0(3)\|1(249)\|2(84)\|4(3)\| | * | 99.1%[336/339] | 1.265486726 |
| Cluster167 | 0(3)\|1(249)\|2(84)\|4(3)\| | * | 99.1%[336/339] | 1.265486726 |
| Cluster441 | 0(19)\|1(215)\|2(104)\|3(1)\| | * | 94.4%[320/339] | 1.256637168 |
| Cluster336 | 0(9)\|1(239)\|2(86)\|3(5)\| | * | 97.3%[330/339] | 1.256637168 |
| Cluster276 | 0(46)\|1(165)\|2(124)\|3(4)\| | * | 86.4%[293/339] | 1.253687316 |
| Cluster260 | 0(1)\|1(260)\|2(76)\|3(2)\| | * | 99.7%[338/339] | 1.233038348 |
| Cluster548 | 0(9)\|1(251)\|2(76)\|3(3)\| | * | 97.3%[330/339] | 1.215339233 |
| Cluster387 | 0(11)\|1(247)\|2(78)\|3(3)\| | * | 96.8%[328/339] | 1.215339233 |
| Cluster291 | 0(13)\|1(244)\|2(78)\|3(4)\| | * | 96.2%[326/339] | 1.215339233 |
| Cluster71 | 0(58)\|1(155)\|2(124)\|3(2)\| | * | 82.9%[281/339] | 1.206489676 |
| Cluster545 | 0(25)\|1(222)\|2(90)\|3(2)\| | * | 92.6%[314/339] | 1.203539823 |
| Cluster53 | 0(23)\|1(229)\|2(84)\|3(1)\|4(2)\| | * | 93.2%[316/339] | 1.203539823 |
| Cluster168 | 0(39)\|1(232)\|2(37)\|3(28)\|4(1)\|5(2)\| | * | 88.5%[300/339] | 1.191740413 |
| Cluster467 | 1(297)\|2(37)\|3(5)\| | * | 100.0%[339/339] | 1.138643068 |
| Cluster191 | 0(3)\|1(286)\|2(50)\| | * | 99.1%[336/339] | 1.138643068 |
| Cluster587 | 1(300)\|2(39)\| | * | 100.0%[339/339] | 1.115044248 |
| Cluster172 | 1(300)\|2(39)\| | * | 100.0%[339/339] | 1.115044248 |
| Cluster455 | 0(41)\|1(225)\|2(73)\| | * | 87.9%[298/339] | 1.09439528 |
| Cluster531 | 0(9)\|1(303)\|2(21)\|4(6)\| | * | 97.3%[330/339] | 1.088495575 |
| Cluster562 | 1(317)\|2(22)\| | * | 100.0%[339/339] | 1.064896755 |
| Cluster371 | 0(17)\|1(290)\|2(30)\|3(1)\|4(1)\| | * | 95.0%[322/339] | 1.053097345 |
| Cluster206 | 0(9)\|1(309)\|2(18)\|4(3)\| | * | 97.3%[330/339] | 1.053097345 |
| Cluster20 | 0(10)\|1(308)\|2(18)\|4(3)\| | * | 97.1%[329/339] | 1.050147493 |
| Cluster465 | 0(51)\|1(232)\|2(45)\|3(11)\| | * | 85.0%[288/339] | 1.04719764 |

| | | | | |
|---|---|---|---|---|
| Cluster158 | 0(51)\|1(232)\|2(45)\|3(11)\| | * | 85.0%[288/339] | 1.04719764 |
| Cluster490 | 0(42)\|1(240)\|2(57)\| | * | 87.6%[297/339] | 1.044247788 |
| Cluster198 | 0(11)\|1(308)\|2(17)\|4(3)\| | * | 96.8%[328/339] | 1.044247788 |
| Cluster157 | 0(52)\|1(233)\|2(43)\|3(11)\| | * | 84.7%[287/339] | 1.038348083 |
| Cluster541 | 1(331)\|2(8)\| | * | 100.0%[339/339] | 1.02359882 |
| Cluster420 | 0(1)\|1(336)\|3(1)\|7(1)\| | * | 99.7%[338/339] | 1.020648968 |
| Cluster591 | 0(19)\|1(295)\|2(25)\| | * | 94.4%[320/339] | 1.017699115 |
| Cluster506 | 0(1)\|1(332)\|2(5)\|3(1)\| | * | 99.7%[338/339] | 1.017699115 |
| Cluster346 | 0(4)\|1(326)\|2(9)\| | * | 98.8%[335/339] | 1.014749263 |
| Cluster121 | 1(337)\|2(1)\|5(1)\| | * | 100.0%[339/339] | 1.014749263 |
| Cluster677 | 1(338)\|4(1)\| | * | 100.0%[339/339] | 1.008849558 |
| Cluster674 | 0(2)\|1(332)\|2(5)\| | * | 99.4%[337/339] | 1.008849558 |
| Cluster661 | 1(336)\|2(3)\| | * | 100.0%[339/339] | 1.008849558 |
| Cluster35 | 0(2)\|1(333)\|2(3)\|3(1)\| | * | 99.4%[337/339] | 1.008849558 |
| Cluster671 | 1(337)\|2(2)\| | * | 100.0%[339/339] | 1.005899705 |
| Cluster653 | 1(337)\|2(2)\| | * | 100.0%[339/339] | 1.005899705 |
| Cluster561 | 1(337)\|2(2)\| | * | 100.0%[339/339] | 1.005899705 |
| Cluster525 | 1(337)\|2(2)\| | * | 100.0%[339/339] | 1.005899705 |
| Cluster46 | 0(4)\|1(329)\|2(6)\| | * | 98.8%[335/339] | 1.005899705 |
| Cluster383 | 1(337)\|2(2)\| | * | 100.0%[339/339] | 1.005899705 |
| Cluster358 | 1(337)\|2(2)\| | * | 100.0%[339/339] | 1.005899705 |
| Cluster264 | 1(337)\|2(2)\| | * | 100.0%[339/339] | 1.005899705 |
| Cluster194 | 0(1)\|1(335)\|2(3)\| | * | 99.7%[338/339] | 1.005899705 |
| Cluster162 | 0(1)\|1(335)\|2(3)\| | * | 99.7%[338/339] | 1.005899705 |
| Cluster13 | 1(337)\|2(2)\| | * | 100.0%[339/339] | 1.005899705 |

| | | | | |
|---|---|---|---|---|
| Cluster112 | 0(1)\|1(335)\|2(3)\| | * | 99.7%[338/339] | 1.005899705 |
| Cluster9 | 0(1)\|1(337)\|3(1)\| | * | 99.7%[338/339] | 1.002949853 |
| Cluster643 | 1(338)\|2(1)\| | * | 100.0%[339/339] | 1.002949853 |
| Cluster639 | 1(338)\|2(1)\| | * | 100.0%[339/339] | 1.002949853 |
| Cluster627 | 1(338)\|2(1)\| | * | 100.0%[339/339] | 1.002949853 |
| Cluster598 | 0(2)\|1(334)\|2(3)\| | * | 99.4%[337/339] | 1.002949853 |
| Cluster501 | 1(338)\|2(1)\| | * | 100.0%[339/339] | 1.002949853 |
| Cluster499 | 1(338)\|2(1)\| | * | 100.0%[339/339] | 1.002949853 |
| Cluster486 | 1(338)\|2(1)\| | * | 100.0%[339/339] | 1.002949853 |
| Cluster468 | 1(338)\|2(1)\| | * | 100.0%[339/339] | 1.002949853 |
| Cluster463 | 1(338)\|2(1)\| | * | 100.0%[339/339] | 1.002949853 |
| Cluster38 | 0(1)\|1(336)\|2(2)\| | * | 99.7%[338/339] | 1.002949853 |
| Cluster343 | 1(338)\|2(1)\| | * | 100.0%[339/339] | 1.002949853 |
| Cluster34 | 1(338)\|2(1)\| | * | 100.0%[339/339] | 1.002949853 |
| Cluster33 | 1(338)\|2(1)\| | * | 100.0%[339/339] | 1.002949853 |
| Cluster325 | 1(338)\|2(1)\| | * | 100.0%[339/339] | 1.002949853 |
| Cluster265 | 1(338)\|2(1)\| | * | 100.0%[339/339] | 1.002949853 |
| Cluster174 | 1(338)\|2(1)\| | * | 100.0%[339/339] | 1.002949853 |
| Cluster146 | 0(1)\|1(336)\|2(2)\| | * | 99.7%[338/339] | 1.002949853 |
| Cluster133 | 1(338)\|2(1)\| | * | 100.0%[339/339] | 1.002949853 |
| Cluster645 | 1(339)\| | * | 100.0%[339/339] | 1 |
| Cluster641 | 0(2)\|1(335)\|2(2)\| | * | 99.4%[337/339] | 1 |
| Cluster524 | 0(51)\|1(237)\|2(51)\| | * | 85.0%[288/339] | 1 |
| Cluster502 | 0(2)\|1(335)\|2(2)\| | * | 99.4%[337/339] | 1 |
| Cluster498 | 1(339)\| | * | 100.0%[339/339] | 1 |

| | | | | |
|---|---|---|---|---|
| Cluster487 | 0(1)\|1(337)\|2(1)\| | * | 99.7%[338/339] | 1 |
| Cluster430 | 1(339)\| | * | 100.0%[339/339] | 1 |
| Cluster423 | 1(339)\| | * | 100.0%[339/339] | 1 |
| Cluster286 | 0(1)\|1(337)\|2(1)\| | * | 99.7%[338/339] | 1 |
| Cluster285 | 0(1)\|1(337)\|2(1)\| | * | 99.7%[338/339] | 1 |
| Cluster254 | 1(339)\| | * | 100.0%[339/339] | 1 |
| Cluster23 | 0(1)\|1(337)\|2(1)\| | * | 99.7%[338/339] | 1 |
| Cluster228 | 0(2)\|1(335)\|2(2)\| | * | 99.4%[337/339] | 1 |
| Cluster199 | 0(3)\|1(333)\|2(3)\| | * | 99.1%[336/339] | 1 |
| Cluster156 | 0(4)\|1(331)\|2(4)\| | * | 98.8%[335/339] | 1 |
| Cluster142 | 1(339)\| | * | 100.0%[339/339] | 1 |
| Cluster141 | 1(339)\| | * | 100.0%[339/339] | 1 |
| Cluster124 | 0(1)\|1(337)\|2(1)\| | * | 99.7%[338/339] | 1 |
| Cluster11 | 0(1)\|1(337)\|2(1)\| | * | 99.7%[338/339] | 1 |
| Cluster669 | 0(2)\|1(336)\|2(1)\| | * | 99.4%[337/339] | 0.997050147 |
| Cluster644 | 0(2)\|1(336)\|2(1)\| | * | 99.4%[337/339] | 0.997050147 |
| Cluster610 | 0(3)\|1(334)\|2(2)\| | * | 99.1%[336/339] | 0.997050147 |
| Cluster504 | 0(2)\|1(336)\|2(1)\| | * | 99.4%[337/339] | 0.997050147 |
| Cluster188 | 0(2)\|1(336)\|2(1)\| | * | 99.4%[337/339] | 0.997050147 |
| Cluster186 | 0(3)\|1(334)\|2(2)\| | * | 99.1%[336/339] | 0.997050147 |
| Cluster540 | 0(3)\|1(335)\|2(1)\| | * | 99.1%[336/339] | 0.994100295 |
| Cluster533 | 0(3)\|1(335)\|2(1)\| | * | 99.1%[336/339] | 0.994100295 |
| Cluster527 | 0(2)\|1(337)\| | * | 99.4%[337/339] | 0.994100295 |
| Cluster505 | 0(3)\|1(335)\|2(1)\| | * | 99.1%[336/339] | 0.994100295 |
| Cluster476 | 0(2)\|1(337)\| | * | 99.4%[337/339] | 0.994100295 |

| | | | | |
|---|---|---|---|---|
| Cluster421 | 0(2)\|1(337)\| | * | 99.4%[337/339] | 0.994100295 |
| Cluster409 | 0(4)\|1(333)\|2(2)\| | * | 98.8%[335/339] | 0.994100295 |
| Cluster293 | 0(2)\|1(337)\| | * | 99.4%[337/339] | 0.994100295 |
| Cluster151 | 0(2)\|1(337)\| | * | 99.4%[337/339] | 0.994100295 |
| Cluster617 | 0(4)\|1(334)\|2(1)\| | * | 98.8%[335/339] | 0.991150442 |
| Cluster362 | 0(5)\|1(332)\|2(2)\| | * | 98.5%[334/339] | 0.991150442 |
| Cluster332 | 0(3)\|1(336)\| | * | 99.1%[336/339] | 0.991150442 |
| Cluster259 | 0(25)\|1(292)\|2(22)\| | * | 92.6%[314/339] | 0.991150442 |
| Cluster655 | 0(4)\|1(335)\| | * | 98.8%[335/339] | 0.98820059 |
| Cluster534 | 0(4)\|1(335)\| | * | 98.8%[335/339] | 0.98820059 |
| Cluster427 | 0(4)\|1(335)\| | * | 98.8%[335/339] | 0.98820059 |
| Cluster185 | 0(19)\|1(305)\|2(15)\| | * | 94.4%[320/339] | 0.98820059 |
| Cluster626 | 0(5)\|1(334)\| | * | 98.5%[334/339] | 0.985250737 |
| Cluster434 | 0(6)\|1(332)\|2(1)\| | * | 98.2%[333/339] | 0.985250737 |
| Cluster386 | 0(5)\|1(334)\| | * | 98.5%[334/339] | 0.985250737 |
| Cluster340 | 0(5)\|1(334)\| | * | 98.5%[334/339] | 0.985250737 |
| Cluster225 | 0(5)\|1(334)\| | * | 98.5%[334/339] | 0.985250737 |
| Cluster19 | 0(5)\|1(334)\| | * | 98.5%[334/339] | 0.985250737 |
| Cluster428 | 0(6)\|1(333)\| | * | 98.2%[333/339] | 0.982300885 |
| Cluster477 | 0(8)\|1(330)\|2(1)\| | * | 97.6%[331/339] | 0.979351032 |
| Cluster432 | 0(8)\|1(330)\|2(1)\| | * | 97.6%[331/339] | 0.979351032 |
| Cluster425 | 0(8)\|1(330)\|2(1)\| | * | 97.6%[331/339] | 0.979351032 |
| Cluster597 | 0(8)\|1(331)\| | * | 97.6%[331/339] | 0.97640118 |
| Cluster422 | 0(19)\|1(313)\|2(5)\|4(2)\| | * | 94.4%[320/339] | 0.97640118 |
| Cluster236 | 0(11)\|1(326)\|2(1)\|3(1)\| | * | 96.8%[328/339] | 0.97640118 |

| | | | | |
|---|---|---|---|---|
| Cluster183 | 0(9)\|1(329)\|2(1)\| | * | 97.3%[330/339] | 0.97640118 |
| Cluster687 | 0(9)\|1(330)\| | * | 97.3%[330/339] | 0.973451327 |
| Cluster216 | 0(11)\|1(326)\|2(2)\| | * | 96.8%[328/339] | 0.973451327 |
| Cluster579 | 0(49)\|1(253)\|2(35)\|3(2)\| | * | 85.5%[290/339] | 0.970501475 |
| Cluster110 | 0(12)\|1(325)\|2(2)\| | * | 96.5%[327/339] | 0.970501475 |
| Cluster549 | 0(15)\|1(320)\|2(4)\| | * | 95.6%[324/339] | 0.967551622 |
| Cluster147 | 0(17)\|1(316)\|2(6)\| | * | 95.0%[322/339] | 0.967551622 |
| Cluster308 | 0(14)\|1(323)\|2(2)\| | * | 95.9%[325/339] | 0.96460177 |
| Cluster407 | 0(13)\|1(326)\| | * | 96.2%[326/339] | 0.961651917 |
| Cluster364 | 0(14)\|1(324)\|2(1)\| | * | 95.9%[325/339] | 0.961651917 |
| Cluster355 | 0(16)\|1(320)\|2(3)\| | * | 95.3%[323/339] | 0.961651917 |
| Cluster415 | 0(16)\|1(321)\|2(2)\| | * | 95.3%[323/339] | 0.958702065 |
| Cluster92 | 0(17)\|1(321)\|2(1)\| | * | 95.0%[322/339] | 0.95280236 |
| Cluster3 | 0(26)\|1(303)\|2(10)\| | * | 92.3%[313/339] | 0.95280236 |
| Cluster244 | 0(16)\|1(323)\| | * | 95.3%[323/339] | 0.95280236 |
| Cluster114 | 0(21)\|1(313)\|2(5)\| | * | 93.8%[318/339] | 0.95280236 |
| Cluster31 | 0(51)\|1(256)\|2(30)\|3(2)\| | * | 85.0%[288/339] | 0.949852507 |
| Cluster323 | 0(19)\|1(319)\|2(1)\| | * | 94.4%[320/339] | 0.946902655 |
| Cluster256 | 0(40)\|1(277)\|2(22)\| | * | 88.2%[299/339] | 0.946902655 |
| Cluster419 | 0(20)\|1(318)\|2(1)\| | * | 94.1%[319/339] | 0.943952802 |
| Cluster234 | 0(28)\|1(302)\|2(9)\| | * | 91.7%[311/339] | 0.943952802 |
| Cluster220 | 0(30)\|1(299)\|2(10)\| | * | 91.2%[309/339] | 0.94100295 |
| Cluster616 | 0(23)\|1(315)\|3(1)\| | * | 93.2%[316/339] | 0.938053097 |
| Cluster418 | 0(22)\|1(316)\|2(1)\| | * | 93.5%[317/339] | 0.938053097 |
| Cluster113 | 0(27)\|1(306)\|2(6)\| | * | 92.0%[312/339] | 0.938053097 |

| | | | | |
|---|---|---|---|---|
| Cluster237 | 0(22)\|1(317)\| | * | 93.5%[317/339] | 0.935103245 |
| Cluster215 | 0(26)\|1(309)\|2(4)\| | * | 92.3%[313/339] | 0.935103245 |
| Cluster57 | 0(29)\|1(304)\|2(6)\| | * | 91.4%[310/339] | 0.932153392 |
| Cluster509 | 0(24)\|1(314)\|2(1)\| | * | 92.9%[315/339] | 0.932153392 |
| Cluster375 | 0(30)\|1(303)\|2(6)\| | * | 91.2%[309/339] | 0.92920354 |
| Cluster123 | 0(27)\|1(309)\|2(3)\| | * | 92.0%[312/339] | 0.92920354 |
| Cluster278 | 0(27)\|1(310)\|2(2)\| | * | 92.0%[312/339] | 0.926253687 |
| Cluster224 | 0(27)\|1(310)\|2(2)\| | * | 92.0%[312/339] | 0.926253687 |
| Cluster43 | 0(36)\|1(296)\|2(6)\|5(1)\| | * | 89.4%[303/339] | 0.923303835 |
| Cluster214 | 0(27)\|1(311)\|2(1)\| | * | 92.0%[312/339] | 0.923303835 |
| Cluster14 | 0(27)\|1(311)\|2(1)\| | * | 92.0%[312/339] | 0.923303835 |
| Cluster456 | 0(29)\|1(308)\|2(2)\| | * | 91.4%[310/339] | 0.920353982 |
| Cluster106 | 0(28)\|1(310)\|2(1)\| | * | 91.7%[311/339] | 0.920353982 |
| Cluster481 | 0(33)\|1(301)\|2(5)\| | * | 90.3%[306/339] | 0.91740413 |
| Cluster108 | 0(38)\|1(295)\|2(5)\|4(1)\| | * | 88.8%[301/339] | 0.911504425 |
| Cluster378 | 0(32)\|1(306)\|2(1)\| | * | 90.6%[307/339] | 0.908554572 |
| Cluster127 | 0(37)\|1(297)\|2(4)\|3(1)\| | * | 89.1%[302/339] | 0.908554572 |
| Cluster536 | 0(39)\|1(295)\|2(4)\|4(1)\| | * | 88.5%[300/339] | 0.90560472 |
| Cluster479 | 0(37)\|1(298)\|2(4)\| | * | 89.1%[302/339] | 0.902654867 |
| Cluster342 | 0(55)\|1(264)\|2(19)\|3(1)\| | * | 83.8%[284/339] | 0.899705015 |
| Cluster470 | 0(37)\|1(301)\|2(1)\| | * | 89.1%[302/339] | 0.89380531 |
| Cluster603 | 0(47)\|1(283)\|2(9)\| | * | 86.1%[292/339] | 0.887905605 |
| Cluster270 | 0(40)\|1(297)\|2(2)\| | * | 88.2%[299/339] | 0.887905605 |
| Cluster503 | 0(50)\|1(280)\|2(9)\| | * | 85.3%[289/339] | 0.879056047 |
| Cluster660 | 0(51)\|1(281)\|2(7)\| | * | 85.0%[288/339] | 0.87020649 |

| | | | | |
|---|---|---|---|---|
| Cluster445 | 0(49)\|1(285)\|2(5)\| | * | 85.5%[290/339] | 0.87020649 |
| Cluster444 | 0(49)\|1(285)\|2(5)\| | * | 85.5%[290/339] | 0.87020649 |
| Cluster219 | 0(45)\|1(293)\|2(1)\| | * | 86.7%[294/339] | 0.87020649 |
| Cluster301 | 0(47)\|1(291)\|2(1)\| | * | 86.1%[292/339] | 0.864306785 |
| Cluster614 | 0(48)\|1(290)\|2(1)\| | * | 85.8%[291/339] | 0.861356932 |
| Cluster566 | 0(51)\|1(286)\|2(2)\| | * | 85.0%[288/339] | 0.855457227 |
| Cluster331 | 0(53)\|1(284)\|2(1)\|4(1)\| | * | 84.4%[286/339] | 0.855457227 |
| Cluster388 | 0(52)\|1(285)\|2(2)\| | * | 84.7%[287/339] | 0.852507375 |
| Cluster529 | 0(54)\|1(283)\|2(2)\| | * | 84.1%[285/339] | 0.84660767 |
| Cluster607 | 0(64)\|1(268)\|2(7)\| | * | 81.1%[275/339] | 0.831858407 |
| Cluster341 | 0(66)\|1(265)\|2(8)\| | * | 80.5%[273/339] | 0.8289085545 |

**Supplementary Table2.**

**Cluster sets of identified undiscovered multicopy regions from B. pertussis, B. parapertussis, B. holmesii.**

| ClusterID | Distribution | Marker | Percent_of_strains | Weighted_Average_Copy |
|---|---|---|---|---|
| Cluster43 | 32(84)\| | * | 100.0%[84/84] | 32 |
| Cluster38 | 32(84)\| | * | 100.0%[84/84] | 32 |
| Cluster33 | 31(1)\|32(83)\| | * | 100.0%[84/84] | 31.98809524 |
| Cluster159 | 31(1)\|32(83)\| | * | 100.0%[84/84] | 31.98809524 |
| Cluster198 | 19(1)\|32(83)\| | * | 100.0%[84/84] | 31.8452381 |
| Cluster59 | 18(1)\|32(83)\| | * | 100.0%[84/84] | 31.83333333 |
| Cluster208 | 18(1)\|32(83)\| | * | 100.0%[84/84] | 31.83333333 |
| Cluster90 | 14(1)\|32(83)\| | * | 100.0%[84/84] | 31.78571429 |
| Cluster57 | 8(1)\|32(83)\| | * | 100.0%[84/84] | 31.71428571 |
| Cluster73 | 4(1)\|32(83)\| | * | 100.0%[84/84] | 31.66666667 |
| Cluster25 | 4(1)\|32(83)\| | * | 100.0%[84/84] | 31.66666667 |
| Cluster160 | 2(1)\|32(83)\| | * | 100.0%[84/84] | 31.64285714 |
| Cluster37 | 1(1)\|32(83)\| | * | 100.0%[84/84] | 31.63095238 |
| Cluster24 | 1(1)\|32(83)\| | * | 100.0%[84/84] | 31.63095238 |
| Cluster64 | 0(1)\|32(83)\| | * | 98.8%[83/84] | 31.61904762 |
| Cluster63 | 0(1)\|32(83)\| | * | 98.8%[83/84] | 31.61904762 |
| Cluster62 | 0(1)\|32(83)\| | * | 98.8%[83/84] | 31.61904762 |
| Cluster60 | 0(1)\|32(83)\| | * | 98.8%[83/84] | 31.61904762 |
| Cluster58 | 0(1)\|32(83)\| | * | 98.8%[83/84] | 31.61904762 |
| Cluster49 | 0(1)\|32(83)\| | * | 98.8%[83/84] | 31.61904762 |

| | | | | |
|---|---|---|---|---|
| Cluster48 | 0(1)\|32(83)\| | * | 98.8%[83/84] | 31.61904762 |
| Cluster45 | 0(1)\|32(83)\| | * | 98.8%[83/84] | 31.61904762 |
| Cluster44 | 0(1)\|32(83)\| | * | 98.8%[83/84] | 31.61904762 |
| Cluster40 | 0(1)\|32(83)\| | * | 98.8%[83/84] | 31.61904762 |
| Cluster39 | 0(1)\|32(83)\| | * | 98.8%[83/84] | 31.61904762 |
| Cluster35 | 0(1)\|32(83)\| | * | 98.8%[83/84] | 31.61904762 |
| Cluster34 | 0(1)\|32(83)\| | * | 98.8%[83/84] | 31.61904762 |
| Cluster29 | 0(1)\|32(83)\| | * | 98.8%[83/84] | 31.61904762 |
| Cluster28 | 0(1)\|32(83)\| | * | 98.8%[83/84] | 31.61904762 |
| Cluster26 | 0(1)\|32(83)\| | * | 98.8%[83/84] | 31.61904762 |
| Cluster22 | 0(1)\|32(83)\| | * | 98.8%[83/84] | 31.61904762 |
| Cluster21 | 0(1)\|32(83)\| | * | 98.8%[83/84] | 31.61904762 |
| Cluster207 | 0(1)\|32(83)\| | * | 98.8%[83/84] | 31.61904762 |
| Cluster199 | 0(1)\|32(83)\| | * | 98.8%[83/84] | 31.61904762 |
| Cluster167 | 0(1)\|32(83)\| | * | 98.8%[83/84] | 31.61904762 |
| Cluster164 | 0(1)\|32(83)\| | * | 98.8%[83/84] | 31.61904762 |
| Cluster161 | 0(1)\|32(83)\| | * | 98.8%[83/84] | 31.61904762 |
| Cluster16 | 0(1)\|32(83)\| | * | 98.8%[83/84] | 31.61904762 |
| Cluster15 | 0(1)\|32(83)\| | * | 98.8%[83/84] | 31.61904762 |
| Cluster14 | 0(1)\|32(83)\| | * | 98.8%[83/84] | 31.61904762 |
| Cluster13 | 0(1)\|32(83)\| | * | 98.8%[83/84] | 31.61904762 |
| Cluster12 | 0(1)\|32(83)\| | * | 98.8%[83/84] | 31.61904762 |
| Cluster120 | 0(1)\|32(83)\| | * | 98.8%[83/84] | 31.61904762 |
| Cluster113 | 0(1)\|32(83)\| | * | 98.8%[83/84] | 31.61904762 |
| Cluster53 | 0(1)\|31(4)\|32(79)\| | * | 98.8%[83/84] | 31.57142857 |

| | | | | |
|---|---|---|---|---|
| Cluster27 | 0(1)\|30(2)\|31(71)\|32(10)\| | * | 98.8%[83/84] | 30.72619048 |
| Cluster80 | 2(1)\|17(83)\| | * | 100.0%[84/84] | 16.82142857 |
| Cluster77 | 2(1)\|17(83)\| | * | 100.0%[84/84] | 16.82142857 |
| Cluster119 | 2(1)\|17(83)\| | * | 100.0%[84/84] | 16.82142857 |
| Cluster215 | 15(1)\|16(83)\| | * | 100.0%[84/84] | 15.98809524 |
| Cluster181 | 15(2)\|16(82)\| | * | 100.0%[84/84] | 15.97619048 |
| Cluster1 | 15(2)\|16(82)\| | * | 100.0%[84/84] | 15.97619048 |
| Cluster10 | 15(2)\|16(82)\| | * | 100.0%[84/84] | 15.97619048 |
| Cluster23 | 0(1)\|15(1)\|16(82)\| | * | 98.8%[83/84] | 15.79761905 |
| Cluster17 | 0(1)\|15(1)\|16(82)\| | * | 98.8%[83/84] | 15.79761905 |
| Cluster20 | 0(1)\|8(1)\|9(1)\|10(4)\|11(9)\|12(18)\|13(16)\|14(20)\|15(7)\|16(5)\|18(2)\| | * | 98.8%[83/84] | 12.86904762 |
| Cluster19 | 0(1)\|8(1)\|9(2)\|10(7)\|11(14)\|12(22)\|13(21)\|14(10)\|15(3)\|16(2)\|18(1)\| | * | 98.8%[83/84] | 12.16666667 |
| Cluster79 | 4(1)\|5(2)\|6(81)\| | * | 100.0%[84/84] | 5.952380952 |
| Cluster122 | 4(1)\|5(2)\|6(81)\| | * | 100.0%[84/84] | 5.952380952 |
| Cluster95 | 2(1)\|5(2)\|6(81)\| | * | 100.0%[84/84] | 5.928571429 |
| Cluster74 | 2(1)\|5(2)\|6(81)\| | * | 100.0%[84/84] | 5.928571429 |
| Cluster41 | 2(1)\|5(2)\|6(81)\| | * | 100.0%[84/84] | 5.928571429 |
| Cluster200 | 2(1)\|5(2)\|6(81)\| | * | 100.0%[84/84] | 5.928571429 |
| Cluster156 | 2(1)\|5(2)\|6(81)\| | * | 100.0%[84/84] | 5.928571429 |
| Cluster36 | 2(1)\|4(2)\|5(81)\| | * | 100.0%[84/84] | 4.94047619 |
| Cluster91 | 3(84)\| | * | 100.0%[84/84] | 3 |
| Cluster46 | 3(84)\| | * | 100.0%[84/84] | 3 |
| Cluster30 | 3(84)\| | * | 100.0%[84/84] | 3 |
| Cluster219 | 3(84)\| | * | 100.0%[84/84] | 3 |
| Cluster194 | 3(84)\| | * | 100.0%[84/84] | 3 |

| | | | | |
|---|---|---|---|---|
| Cluster193 | 3(84)\| | * | 100.0%[84/84] | 3 |
| Cluster192 | 3(84)\| | * | 100.0%[84/84] | 3 |
| Cluster191 | 3(84)\| | * | 100.0%[84/84] | 3 |
| Cluster176 | 3(84)\| | * | 100.0%[84/84] | 3 |
| Cluster155 | 3(84)\| | * | 100.0%[84/84] | 3 |
| Cluster154 | 3(84)\| | * | 100.0%[84/84] | 3 |
| Cluster140 | 3(84)\| | * | 100.0%[84/84] | 3 |
| Cluster137 | 3(84)\| | * | 100.0%[84/84] | 3 |
| Cluster135 | 3(84)\| | * | 100.0%[84/84] | 3 |
| Cluster131 | 3(84)\| | * | 100.0%[84/84] | 3 |
| Cluster118 | 3(84)\| | * | 100.0%[84/84] | 3 |
| Cluster110 | 3(84)\| | * | 100.0%[84/84] | 3 |
| Cluster108 | 3(84)\| | * | 100.0%[84/84] | 3 |
| Cluster123 | 0(1)\|1(15)\|2(38)\|3(27)\|4(3)\| | * | 98.8%[83/84] | 2.19047619 |
| Cluster98 | 2(83)\|4(1)\| | * | 100.0%[84/84] | 2.023809524 |
| Cluster97 | 2(83)\|4(1)\| | * | 100.0%[84/84] | 2.023809524 |
| Cluster96 | 2(83)\|4(1)\| | * | 100.0%[84/84] | 2.023809524 |
| Cluster3 | 2(83)\|4(1)\| | * | 100.0%[84/84] | 2.023809524 |
| Cluster2 | 2(83)\|4(1)\| | * | 100.0%[84/84] | 2.023809524 |
| Cluster210 | 2(83)\|4(1)\| | * | 100.0%[84/84] | 2.023809524 |
| Cluster201 | 2(83)\|4(1)\| | * | 100.0%[84/84] | 2.023809524 |
| Cluster169 | 2(83)\|4(1)\| | * | 100.0%[84/84] | 2.023809524 |
| Cluster158 | 2(83)\|4(1)\| | * | 100.0%[84/84] | 2.023809524 |
| Cluster127 | 2(83)\|4(1)\| | * | 100.0%[84/84] | 2.023809524 |
| Cluster124 | 2(83)\|4(1)\| | * | 100.0%[84/84] | 2.023809524 |

| | | | | |
|---|---|---|---|---|
| Cluster115 | 2(83)\|4(1)\| | * | 100.0%[84/84] | 2.023809524 |
| Cluster100 | 2(83)\|4(1)\| | * | 100.0%[84/84] | 2.023809524 |
| Cluster190 | 2(83)\|3(1)\| | * | 100.0%[84/84] | 2.011904762 |
| Cluster153 | 2(83)\|3(1)\| | * | 100.0%[84/84] | 2.011904762 |
| Cluster84 | 2(84)\| | * | 100.0%[84/84] | 2 |
| Cluster71 | 2(84)\| | * | 100.0%[84/84] | 2 |
| Cluster52 | 2(84)\| | * | 100.0%[84/84] | 2 |
| Cluster31 | 1(1)\|2(82)\|3(1)\| | * | 100.0%[84/84] | 2 |
| Cluster217 | 2(84)\| | * | 100.0%[84/84] | 2 |
| Cluster195 | 2(84)\| | * | 100.0%[84/84] | 2 |
| Cluster177 | 2(84)\| | * | 100.0%[84/84] | 2 |
| Cluster175 | 2(84)\| | * | 100.0%[84/84] | 2 |
| Cluster168 | 2(84)\| | * | 100.0%[84/84] | 2 |
| Cluster163 | 2(84)\| | * | 100.0%[84/84] | 2 |
| Cluster126 | 2(84)\| | * | 100.0%[84/84] | 2 |
| Cluster117 | 2(84)\| | * | 100.0%[84/84] | 2 |
| Cluster109 | 2(84)\| | * | 100.0%[84/84] | 2 |
| Cluster72 | 1(1)\|2(83)\| | * | 100.0%[84/84] | 1.988095238 |
| Cluster54 | 0(1)\|1(1)\|2(80)\|3(2)\| | * | 98.8%[83/84] | 1.988095238 |
| Cluster47 | 1(1)\|2(83)\| | * | 100.0%[84/84] | 1.988095238 |
| Cluster216 | 1(1)\|2(83)\| | * | 100.0%[84/84] | 1.988095238 |
| Cluster18 | 1(1)\|2(83)\| | * | 100.0%[84/84] | 1.988095238 |
| Cluster178 | 1(1)\|2(83)\| | * | 100.0%[84/84] | 1.988095238 |
| Cluster142 | 1(1)\|2(83)\| | * | 100.0%[84/84] | 1.988095238 |
| Cluster141 | 0(1)\|1(1)\|2(80)\|3(2)\| | * | 98.8%[83/84] | 1.988095238 |

| | | | | |
|---|---|---|---|---|
| Cluster138 | 1(1)\|2(83)\| | * | 100.0%[84/84] | 1.988095238 |
| Cluster136 | 1(1)\|2(83)\| | * | 100.0%[84/84] | 1.988095238 |
| Cluster114 | 1(1)\|2(83)\| | * | 100.0%[84/84] | 1.988095238 |
| Cluster111 | 1(1)\|2(83)\| | * | 100.0%[84/84] | 1.988095238 |
| Cluster81 | 0(1)\|2(83)\| | * | 98.8%[83/84] | 1.976190476 |
| Cluster42 | 0(1)\|2(83)\| | * | 98.8%[83/84] | 1.976190476 |
| Cluster89 | 0(1)\|1(1)\|2(82)\| | * | 98.8%[83/84] | 1.964285714 |
| Cluster88 | 0(1)\|1(1)\|2(82)\| | * | 98.8%[83/84] | 1.964285714 |
| Cluster197 | 0(1)\|1(1)\|2(82)\| | * | 98.8%[83/84] | 1.964285714 |
| Cluster87 | 1(5)\|2(79)\| | * | 100.0%[84/84] | 1.94047619 |
| Cluster206 | 1(6)\|2(78)\| | * | 100.0%[84/84] | 1.928571429 |
| Cluster174 | 1(9)\|2(75)\| | * | 100.0%[84/84] | 1.892857143 |
| Cluster61 | 1(22)\|2(62)\| | * | 100.0%[84/84] | 1.738095238 |
| Cluster92 | 1(61)\|2(23)\| | * | 100.0%[84/84] | 1.273809524 |
| Cluster56 | 1(65)\|2(19)\| | * | 100.0%[84/84] | 1.226190476 |
| Cluster182 | 1(70)\|2(14)\| | * | 100.0%[84/84] | 1.166666667 |
| Cluster205 | 1(80)\|2(4)\| | * | 100.0%[84/84] | 1.047619048 |
| Cluster86 | 1(82)\|2(2)\| | * | 100.0%[84/84] | 1.023809524 |
| Cluster75 | 1(83)\|2(1)\| | * | 100.0%[84/84] | 1.011904762 |
| Cluster165 | 1(83)\|2(1)\| | * | 100.0%[84/84] | 1.011904762 |
| Cluster139 | 1(83)\|2(1)\| | * | 100.0%[84/84] | 1.011904762 |
| Cluster112 | 1(83)\|2(1)\| | * | 100.0%[84/84] | 1.011904762 |
| Cluster99 | 1(84)\| | * | 100.0%[84/84] | 1 |
| Cluster94 | 1(84)\| | * | 100.0%[84/84] | 1 |
| Cluster93 | 1(84)\| | * | 100.0%[84/84] | 1 |

| | | | | |
|---|---|---|---|---|
| Cluster9 | 1(84)\| | * | 100.0%[84/84] | 1 |
| Cluster85 | 1(84)\| | * | 100.0%[84/84] | 1 |
| Cluster83 | 1(84)\| | * | 100.0%[84/84] | 1 |
| Cluster82 | 1(84)\| | * | 100.0%[84/84] | 1 |
| Cluster8 | 1(84)\| | * | 100.0%[84/84] | 1 |
| Cluster78 | 1(84)\| | * | 100.0%[84/84] | 1 |
| Cluster76 | 1(84)\| | * | 100.0%[84/84] | 1 |
| Cluster70 | 1(84)\| | * | 100.0%[84/84] | 1 |
| Cluster69 | 1(84)\| | * | 100.0%[84/84] | 1 |
| Cluster67 | 1(84)\| | * | 100.0%[84/84] | 1 |
| Cluster66 | 1(84)\| | * | 100.0%[84/84] | 1 |
| Cluster65 | 1(84)\| | * | 100.0%[84/84] | 1 |
| Cluster6 | 1(84)\| | * | 100.0%[84/84] | 1 |
| Cluster55 | 1(84)\| | * | 100.0%[84/84] | 1 |
| Cluster5 | 1(84)\| | * | 100.0%[84/84] | 1 |
| Cluster51 | 1(84)\| | * | 100.0%[84/84] | 1 |
| Cluster50 | 0(1)\|1(82)\|2(1)\| | * | 98.8%[83/84] | 1 |
| Cluster4 | 1(84)\| | * | 100.0%[84/84] | 1 |
| Cluster32 | 1(84)\| | * | 100.0%[84/84] | 1 |
| Cluster222 | 1(84)\| | * | 100.0%[84/84] | 1 |
| Cluster221 | 1(84)\| | * | 100.0%[84/84] | 1 |
| Cluster220 | 1(84)\| | * | 100.0%[84/84] | 1 |
| Cluster218 | 1(84)\| | * | 100.0%[84/84] | 1 |
| Cluster213 | 1(84)\| | * | 100.0%[84/84] | 1 |
| Cluster212 | 1(84)\| | * | 100.0%[84/84] | 1 |

| | | | | |
|---|---|---|---|---|
| Cluster211 | 1(84)\| | * | 100.0%[84/84] | 1 |
| Cluster209 | 1(84)\| | * | 100.0%[84/84] | 1 |
| Cluster204 | 1(84)\| | * | 100.0%[84/84] | 1 |
| Cluster203 | 1(84)\| | * | 100.0%[84/84] | 1 |
| Cluster202 | 1(84)\| | * | 100.0%[84/84] | 1 |
| Cluster196 | 1(84)\| | * | 100.0%[84/84] | 1 |
| Cluster189 | 1(84)\| | * | 100.0%[84/84] | 1 |
| Cluster188 | 1(84)\| | * | 100.0%[84/84] | 1 |
| Cluster187 | 1(84)\| | * | 100.0%[84/84] | 1 |
| Cluster186 | 1(84)\| | * | 100.0%[84/84] | 1 |
| Cluster185 | 1(84)\| | * | 100.0%[84/84] | 1 |
| Cluster184 | 1(84)\| | * | 100.0%[84/84] | 1 |
| Cluster183 | 1(84)\| | * | 100.0%[84/84] | 1 |
| Cluster179 | 1(84)\| | * | 100.0%[84/84] | 1 |
| Cluster173 | 1(84)\| | * | 100.0%[84/84] | 1 |
| Cluster172 | 1(84)\| | * | 100.0%[84/84] | 1 |
| Cluster171 | 1(84)\| | * | 100.0%[84/84] | 1 |
| Cluster170 | 1(84)\| | * | 100.0%[84/84] | 1 |
| Cluster166 | 0(1)\|1(82)\|2(1)\| | * | 98.8%[83/84] | 1 |
| Cluster162 | 1(84)\| | * | 100.0%[84/84] | 1 |
| Cluster157 | 1(84)\| | * | 100.0%[84/84] | 1 |
| Cluster152 | 1(84)\| | * | 100.0%[84/84] | 1 |
| Cluster151 | 1(84)\| | * | 100.0%[84/84] | 1 |
| Cluster150 | 1(84)\| | * | 100.0%[84/84] | 1 |
| Cluster149 | 1(84)\| | * | 100.0%[84/84] | 1 |

| | | | | |
|---|---|---|---|---|
| Cluster148 | 1(84)\| | * | 100.0%[84/84] | 1 |
| Cluster147 | 1(84)\| | * | 100.0%[84/84] | 1 |
| Cluster146 | 1(84)\| | * | 100.0%[84/84] | 1 |
| Cluster145 | 1(84)\| | * | 100.0%[84/84] | 1 |
| Cluster144 | 1(84)\| | * | 100.0%[84/84] | 1 |
| Cluster143 | 1(84)\| | * | 100.0%[84/84] | 1 |
| Cluster134 | 1(84)\| | * | 100.0%[84/84] | 1 |
| Cluster133 | 1(84)\| | * | 100.0%[84/84] | 1 |
| Cluster132 | 1(84)\| | * | 100.0%[84/84] | 1 |
| Cluster130 | 1(84)\| | * | 100.0%[84/84] | 1 |
| Cluster129 | 1(84)\| | * | 100.0%[84/84] | 1 |
| Cluster128 | 1(84)\| | * | 100.0%[84/84] | 1 |
| Cluster125 | 1(84)\| | * | 100.0%[84/84] | 1 |
| Cluster116 | 1(84)\| | * | 100.0%[84/84] | 1 |
| Cluster11 | 1(84)\| | * | 100.0%[84/84] | 1 |
| Cluster107 | 1(84)\| | * | 100.0%[84/84] | 1 |
| Cluster106 | 1(84)\| | * | 100.0%[84/84] | 1 |
| Cluster105 | 1(84)\| | * | 100.0%[84/84] | 1 |
| Cluster104 | 1(84)\| | * | 100.0%[84/84] | 1 |
| Cluster103 | 1(84)\| | * | 100.0%[84/84] | 1 |
| Cluster102 | 1(84)\| | * | 100.0%[84/84] | 1 |
| Cluster101 | 1(84)\| | * | 100.0%[84/84] | 1 |
| Cluster0 | 1(84)\| | * | 100.0%[84/84] | 1 |
| Cluster7 | 0(7)\|1(77)\| | * | 91.7%[77/84] | 0.916666667 |
| Cluster68 | 0(7)\|1(77)\| | * | 91.7%[77/84] | 0.916666667 |

| | | | | |
|---|---|---|---|---|
| Cluster214 | 0(39)\|1(45)\| | | | 53.6%[45/84] |
| Cluster180 | 0(73)\|1(9)\|2(2)\| | | | 13.1%[11/84] |
| Cluster121 | 0(69)\|1(13)\|2(2)\| | | | 17.9%[15/84] |

**Supplementary Table3.**

**Cluster sets of identified undiscovered multicopy regions from B. parapertussis, B. bronchiseptica.**

| ClusterID | Distribution | Marker | Percent_of_strains | Weighted_Average_Copy |
|---|---|---|---|---|
| Cluster9 | 22(16)\|24(1)\|25(1)\| | * | 100.0%[18/18] | 22.27777778 |
| Cluster8 | 22(16)\|24(1)\|25(1)\| | * | 100.0%[18/18] | 22.27777778 |
| Cluster25 | 22(16)\|24(1)\|25(1)\| | * | 100.0%[18/18] | 22.27777778 |
| Cluster10 | 22(16)\|24(1)\|25(1)\| | * | 100.0%[18/18] | 22.27777778 |
| Cluster19 | 22(16)\|24(2)\| | * | 100.0%[18/18] | 22.22222222 |
| Cluster18 | 22(16)\|24(2)\| | * | 100.0%[18/18] | 22.22222222 |
| Cluster1 | 22(16)\|23(1)\|24(1)\| | * | 100.0%[18/18] | 22.16666667 |
| Cluster0 | 22(16)\|23(1)\|24(1)\| | * | 100.0%[18/18] | 22.16666667 |
| Cluster40 | 2(1)\|3(17)\| | * | 100.0%[18/18] | 2.944444444 |
| Cluster3 | 2(1)\|3(17)\| | * | 100.0%[18/18] | 2.944444444 |
| Cluster28 | 2(1)\|3(17)\| | * | 100.0%[18/18] | 2.944444444 |
| Cluster27 | 2(1)\|3(17)\| | * | 100.0%[18/18] | 2.944444444 |
| Cluster4 | 1(1)\|2(5)\|3(12)\| | * | 100.0%[18/18] | 2.611111111 |
| Cluster37 | 2(11)\|3(7)\| | * | 100.0%[18/18] | 2.388888889 |
| Cluster32 | 2(15)\|3(3)\| | * | 100.0%[18/18] | 2.166666667 |
| Cluster26 | 2(15)\|3(3)\| | * | 100.0%[18/18] | 2.166666667 |
| Cluster63 | 2(18)\| | * | 100.0%[18/18] | 2 |

**Supplementary Table4.**

**Cluster sets of identified undiscovered multicopy regions from M. pneumoniae strain M129.**

| ClusterID | Distribution | Marker | Percent_of_strains | Weighted_Average_Copy |
|---|---|---|---|---|
| Cluster22 | 11(10)\| | * | 100.0%[10/10] | 11 |
| Cluster164 | 11(10)\| | * | 100.0%[10/10] | 11 |
| Cluster141 | 11(10)\| | * | 100.0%[10/10] | 11 |
| Cluster9 | 10(2)\|11(8)\| | * | 100.0%[10/10] | 10.8 |
| Cluster472 | 10(2)\|11(8)\| | * | 100.0%[10/10] | 10.8 |
| Cluster468 | 10(2)\|11(8)\| | * | 100.0%[10/10] | 10.8 |
| Cluster188 | 10(2)\|11(8)\| | * | 100.0%[10/10] | 10.8 |
| Cluster187 | 10(2)\|11(8)\| | * | 100.0%[10/10] | 10.8 |
| Cluster182 | 10(2)\|11(8)\| | * | 100.0%[10/10] | 10.8 |
| Cluster0 | 10(2)\|11(8)\| | * | 100.0%[10/10] | 10.8 |
| Cluster66 | 10(10)\| | * | 100.0%[10/10] | 10 |
| Cluster16 | 10(10)\| | * | 100.0%[10/10] | 10 |
| Cluster92 | 9(2)\|10(8)\| | * | 100.0%[10/10] | 9.8 |
| Cluster512 | 9(2)\|10(8)\| | * | 100.0%[10/10] | 9.8 |
| Cluster43 | 8(1)\|10(9)\| | * | 100.0%[10/10] | 9.8 |
| Cluster446 | 8(2)\|10(8)\| | * | 100.0%[10/10] | 9.6 |
| Cluster421 | 9(10)\| | * | 100.0%[10/10] | 9 |
| Cluster34 | 9(10)\| | * | 100.0%[10/10] | 9 |
| Cluster6 | 8(10)\| | * | 100.0%[10/10] | 8 |
| Cluster51 | 8(10)\| | * | 100.0%[10/10] | 8 |

| | | | | |
|---|---|---|---|---|
| Cluster457 | 8(10)\| | * | 100.0%[10/10] | 8 |
| Cluster413 | 8(10)\| | * | 100.0%[10/10] | 8 |
| Cluster35 | 8(10)\| | * | 100.0%[10/10] | 8 |
| Cluster299 | 8(10)\| | * | 100.0%[10/10] | 8 |
| Cluster198 | 8(10)\| | * | 100.0%[10/10] | 8 |
| Cluster19 | 8(10)\| | * | 100.0%[10/10] | 8 |
| Cluster18 | 8(10)\| | * | 100.0%[10/10] | 8 |
| Cluster13 | 8(10)\| | * | 100.0%[10/10] | 8 |
| Cluster241 | 7(1)\|8(9)\| | * | 100.0%[10/10] | 7.9 |
| Cluster156 | 7(1)\|8(9)\| | * | 100.0%[10/10] | 7.9 |
| Cluster420 | 7(2)\|8(8)\| | * | 100.0%[10/10] | 7.8 |
| Cluster189 | 7(2)\|8(8)\| | * | 100.0%[10/10] | 7.8 |
| Cluster149 | 7(2)\|8(8)\| | * | 100.0%[10/10] | 7.8 |
| Cluster134 | 7(2)\|8(8)\| | * | 100.0%[10/10] | 7.8 |
| Cluster21 | 6(1)\|7(1)\|8(8)\| | * | 100.0%[10/10] | 7.7 |
| Cluster27 | 7(8)\|8(2)\| | * | 100.0%[10/10] | 7.2 |
| Cluster82 | 7(10)\| | * | 100.0%[10/10] | 7 |
| Cluster67 | 7(10)\| | * | 100.0%[10/10] | 7 |
| Cluster54 | 7(10)\| | * | 100.0%[10/10] | 7 |
| Cluster532 | 7(10)\| | * | 100.0%[10/10] | 7 |
| Cluster509 | 7(10)\| | * | 100.0%[10/10] | 7 |
| Cluster388 | 7(10)\| | * | 100.0%[10/10] | 7 |
| Cluster257 | 7(10)\| | * | 100.0%[10/10] | 7 |
| Cluster247 | 7(10)\| | * | 100.0%[10/10] | 7 |
| Cluster239 | 7(10)\| | * | 100.0%[10/10] | 7 |

| | | | | |
|---|---|---|---|---|
| Cluster232 | 7(10)\| | * | 100.0%[10/10] | 7 |
| Cluster230 | 7(10)\| | * | 100.0%[10/10] | 7 |
| Cluster204 | 7(10)\| | * | 100.0%[10/10] | 7 |
| Cluster173 | 7(10)\| | * | 100.0%[10/10] | 7 |
| Cluster125 | 7(10)\| | * | 100.0%[10/10] | 7 |
| Cluster102 | 7(10)\| | * | 100.0%[10/10] | 7 |
| Cluster248 | 6(1)\|7(9)\| | * | 100.0%[10/10] | 6.9 |
| Cluster343 | 6(2)\|7(8)\| | * | 100.0%[10/10] | 6.8 |
| Cluster100 | 6(2)\|7(8)\| | * | 100.0%[10/10] | 6.8 |
| Cluster491 | 5(1)\|6(1)\|7(8)\| | * | 100.0%[10/10] | 6.7 |
| Cluster205 | 5(2)\|7(8)\| | * | 100.0%[10/10] | 6.6 |
| Cluster351 | 5(1)\|6(5)\|7(4)\| | * | 100.0%[10/10] | 6.3 |
| Cluster131 | 6(8)\|7(2)\| | * | 100.0%[10/10] | 6.2 |
| Cluster77 | 6(10)\| | * | 100.0%[10/10] | 6 |
| Cluster531 | 6(10)\| | * | 100.0%[10/10] | 6 |
| Cluster497 | 6(10)\| | * | 100.0%[10/10] | 6 |
| Cluster426 | 6(10)\| | * | 100.0%[10/10] | 6 |
| Cluster416 | 6(10)\| | * | 100.0%[10/10] | 6 |
| Cluster377 | 6(10)\| | * | 100.0%[10/10] | 6 |
| Cluster328 | 6(10)\| | * | 100.0%[10/10] | 6 |
| Cluster226 | 6(10)\| | * | 100.0%[10/10] | 6 |
| Cluster20 | 6(10)\| | * | 100.0%[10/10] | 6 |
| Cluster120 | 6(10)\| | * | 100.0%[10/10] | 6 |
| Cluster119 | 6(10)\| | * | 100.0%[10/10] | 6 |
| Cluster53 | 5(1)\|6(9)\| | * | 100.0%[10/10] | 5.9 |

| | | | | |
|---|---|---|---|---|
| Cluster319 | 5(2)\|6(8)\| | * | 100.0%[10/10] | 5.8 |
| Cluster317 | 5(2)\|6(8)\| | * | 100.0%[10/10] | 5.8 |
| Cluster305 | 5(2)\|6(8)\| | * | 100.0%[10/10] | 5.8 |
| Cluster126 | 5(2)\|6(8)\| | * | 100.0%[10/10] | 5.8 |
| Cluster11 | 5(2)\|6(8)\| | * | 100.0%[10/10] | 5.8 |
| Cluster144 | 5(3)\|6(7)\| | * | 100.0%[10/10] | 5.7 |
| Cluster65 | 5(4)\|6(6)\| | * | 100.0%[10/10] | 5.6 |
| Cluster2 | 4(2)\|6(8)\| | * | 100.0%[10/10] | 5.6 |
| Cluster190 | 5(4)\|6(6)\| | * | 100.0%[10/10] | 5.6 |
| Cluster60 | 5(8)\|6(2)\| | * | 100.0%[10/10] | 5.2 |
| Cluster5 | 5(8)\|6(2)\| | * | 100.0%[10/10] | 5.2 |
| Cluster264 | 5(8)\|6(2)\| | * | 100.0%[10/10] | 5.2 |
| Cluster124 | 5(8)\|6(2)\| | * | 100.0%[10/10] | 5.2 |
| Cluster52 | 5(10)\| | * | 100.0%[10/10] | 5 |
| Cluster506 | 5(10)\| | * | 100.0%[10/10] | 5 |
| Cluster499 | 5(10)\| | * | 100.0%[10/10] | 5 |
| Cluster46 | 4(3)\|5(4)\|6(3)\| | * | 100.0%[10/10] | 5 |
| Cluster453 | 5(10)\| | * | 100.0%[10/10] | 5 |
| Cluster414 | 5(10)\| | * | 100.0%[10/10] | 5 |
| Cluster40 | 5(10)\| | * | 100.0%[10/10] | 5 |
| Cluster384 | 5(10)\| | * | 100.0%[10/10] | 5 |
| Cluster357 | 5(10)\| | * | 100.0%[10/10] | 5 |
| Cluster352 | 5(10)\| | * | 100.0%[10/10] | 5 |
| Cluster315 | 5(10)\| | * | 100.0%[10/10] | 5 |
| Cluster306 | 5(10)\| | * | 100.0%[10/10] | 5 |

| | | | | |
|---|---|---|---|---|
| Cluster289 | 5(10)\| | * | 100.0%[10/10] | 5 |
| Cluster246 | 5(10)\| | * | 100.0%[10/10] | 5 |
| Cluster24 | 5(10)\| | * | 100.0%[10/10] | 5 |
| Cluster176 | 5(10)\| | * | 100.0%[10/10] | 5 |
| Cluster167 | 5(10)\| | * | 100.0%[10/10] | 5 |
| Cluster138 | 5(10)\| | * | 100.0%[10/10] | 5 |
| Cluster123 | 4(1)\|5(8)\|6(1)\| | * | 100.0%[10/10] | 5 |
| Cluster293 | 4(1)\|5(9)\| | * | 100.0%[10/10] | 4.9 |
| Cluster174 | 4(1)\|5(9)\| | * | 100.0%[10/10] | 4.9 |
| Cluster386 | 4(2)\|5(8)\| | * | 100.0%[10/10] | 4.8 |
| Cluster345 | 4(2)\|5(8)\| | * | 100.0%[10/10] | 4.8 |
| Cluster298 | 4(2)\|5(8)\| | * | 100.0%[10/10] | 4.8 |
| Cluster282 | 4(2)\|5(8)\| | * | 100.0%[10/10] | 4.8 |
| Cluster8 | 3(2)\|5(8)\| | * | 100.0%[10/10] | 4.6 |
| Cluster56 | 4(5)\|5(5)\| | * | 100.0%[10/10] | 4.5 |
| Cluster543 | 4(6)\|5(4)\| | * | 100.0%[10/10] | 4.4 |
| Cluster522 | 4(8)\|5(1)\|6(1)\| | * | 100.0%[10/10] | 4.3 |
| Cluster489 | 4(8)\|5(2)\| | * | 100.0%[10/10] | 4.2 |
| Cluster395 | 4(8)\|5(2)\| | * | 100.0%[10/10] | 4.2 |
| Cluster387 | 4(8)\|5(2)\| | * | 100.0%[10/10] | 4.2 |
| Cluster346 | 4(8)\|5(2)\| | * | 100.0%[10/10] | 4.2 |
| Cluster312 | 4(8)\|5(2)\| | * | 100.0%[10/10] | 4.2 |
| Cluster287 | 4(8)\|5(2)\| | * | 100.0%[10/10] | 4.2 |
| Cluster197 | 4(8)\|5(2)\| | * | 100.0%[10/10] | 4.2 |
| Cluster191 | 4(8)\|5(2)\| | * | 100.0%[10/10] | 4.2 |

| | | | | |
|---|---|---|---|---|
| Cluster97 | 4(9)\|5(1)\| | * | 100.0%[10/10] | 4.1 |
| Cluster549 | 4(9)\|5(1)\| | * | 100.0%[10/10] | 4.1 |
| Cluster208 | 4(9)\|5(1)\| | * | 100.0%[10/10] | 4.1 |
| Cluster166 | 4(9)\|5(1)\| | * | 100.0%[10/10] | 4.1 |
| Cluster153 | 4(9)\|5(1)\| | * | 100.0%[10/10] | 4.1 |
| Cluster148 | 4(9)\|5(1)\| | * | 100.0%[10/10] | 4.1 |
| Cluster71 | 4(10)\| | * | 100.0%[10/10] | 4 |
| Cluster57 | 4(10)\| | * | 100.0%[10/10] | 4 |
| Cluster544 | 4(10)\| | * | 100.0%[10/10] | 4 |
| Cluster537 | 4(10)\| | * | 100.0%[10/10] | 4 |
| Cluster533 | 4(10)\| | * | 100.0%[10/10] | 4 |
| Cluster527 | 4(10)\| | * | 100.0%[10/10] | 4 |
| Cluster473 | 4(10)\| | * | 100.0%[10/10] | 4 |
| Cluster465 | 4(10)\| | * | 100.0%[10/10] | 4 |
| Cluster463 | 4(10)\| | * | 100.0%[10/10] | 4 |
| Cluster450 | 4(10)\| | * | 100.0%[10/10] | 4 |
| Cluster443 | 4(10)\| | * | 100.0%[10/10] | 4 |
| Cluster438 | 4(10)\| | * | 100.0%[10/10] | 4 |
| Cluster424 | 4(10)\| | * | 100.0%[10/10] | 4 |
| Cluster417 | 4(10)\| | * | 100.0%[10/10] | 4 |
| Cluster391 | 4(10)\| | * | 100.0%[10/10] | 4 |
| Cluster390 | 4(10)\| | * | 100.0%[10/10] | 4 |
| Cluster385 | 4(10)\| | * | 100.0%[10/10] | 4 |
| Cluster37 | 4(10)\| | * | 100.0%[10/10] | 4 |
| Cluster370 | 4(10)\| | * | 100.0%[10/10] | 4 |

| | | | | |
|---|---|---|---|---|
| Cluster367 | 4(10)\| | * | 100.0%[10/10] | 4 |
| Cluster365 | 4(10)\| | * | 100.0%[10/10] | 4 |
| Cluster360 | 4(10)\| | * | 100.0%[10/10] | 4 |
| Cluster355 | 4(10)\| | * | 100.0%[10/10] | 4 |
| Cluster326 | 4(10)\| | * | 100.0%[10/10] | 4 |
| Cluster32 | 4(10)\| | * | 100.0%[10/10] | 4 |
| Cluster316 | 4(10)\| | * | 100.0%[10/10] | 4 |
| Cluster296 | 4(10)\| | * | 100.0%[10/10] | 4 |
| Cluster284 | 4(10)\| | * | 100.0%[10/10] | 4 |
| Cluster271 | 4(10)\| | * | 100.0%[10/10] | 4 |
| Cluster262 | 4(10)\| | * | 100.0%[10/10] | 4 |
| Cluster240 | 4(10)\| | * | 100.0%[10/10] | 4 |
| Cluster236 | 4(10)\| | * | 100.0%[10/10] | 4 |
| Cluster228 | 4(10)\| | * | 100.0%[10/10] | 4 |
| Cluster217 | 4(10)\| | * | 100.0%[10/10] | 4 |
| Cluster215 | 4(10)\| | * | 100.0%[10/10] | 4 |
| Cluster210 | 4(10)\| | * | 100.0%[10/10] | 4 |
| Cluster206 | 4(10)\| | * | 100.0%[10/10] | 4 |
| Cluster186 | 4(10)\| | * | 100.0%[10/10] | 4 |
| Cluster184 | 4(10)\| | * | 100.0%[10/10] | 4 |
| Cluster118 | 4(10)\| | * | 100.0%[10/10] | 4 |
| Cluster112 | 4(10)\| | * | 100.0%[10/10] | 4 |
| Cluster104 | 4(10)\| | * | 100.0%[10/10] | 4 |
| Cluster72 | 3(2)\|4(8)\| | * | 100.0%[10/10] | 3.8 |
| Cluster64 | 3(2)\|4(8)\| | * | 100.0%[10/10] | 3.8 |

| | | | | |
|---|---|---|---|---|
| Cluster492 | 3(2)\|4(8)\| | * | 100.0%[10/10] | 3.8 |
| Cluster474 | 3(2)\|4(8)\| | * | 100.0%[10/10] | 3.8 |
| Cluster448 | 3(2)\|4(8)\| | * | 100.0%[10/10] | 3.8 |
| Cluster33 | 3(2)\|4(8)\| | * | 100.0%[10/10] | 3.8 |
| Cluster324 | 3(4)\|4(4)\|5(2)\| | * | 100.0%[10/10] | 3.8 |
| Cluster295 | 3(2)\|4(8)\| | * | 100.0%[10/10] | 3.8 |
| Cluster14 | 3(2)\|4(8)\| | * | 100.0%[10/10] | 3.8 |
| Cluster12 | 3(2)\|4(8)\| | * | 100.0%[10/10] | 3.8 |
| Cluster242 | 3(3)\|4(7)\| | * | 100.0%[10/10] | 3.7 |
| Cluster29 | 2(2)\|4(8)\| | * | 100.0%[10/10] | 3.6 |
| Cluster216 | 2(2)\|4(8)\| | * | 100.0%[10/10] | 3.6 |
| Cluster76 | 3(8)\|5(2)\| | * | 100.0%[10/10] | 3.4 |
| Cluster336 | 3(8)\|5(2)\| | * | 100.0%[10/10] | 3.4 |
| Cluster23 | 3(8)\|4(1)\|6(1)\| | * | 100.0%[10/10] | 3.4 |
| Cluster441 | 0(2)\|1(1)\|2(1)\|5(6)\| | * | 80.0%[8/10] | 3.3 |
| Cluster95 | 3(8)\|4(2)\| | * | 100.0%[10/10] | 3.2 |
| Cluster91 | 3(8)\|4(2)\| | * | 100.0%[10/10] | 3.2 |
| Cluster513 | 3(8)\|4(2)\| | * | 100.0%[10/10] | 3.2 |
| Cluster3 | 3(8)\|4(2)\| | * | 100.0%[10/10] | 3.2 |
| Cluster151 | 3(8)\|4(2)\| | * | 100.0%[10/10] | 3.2 |
| Cluster111 | 3(8)\|4(2)\| | * | 100.0%[10/10] | 3.2 |
| Cluster78 | 3(9)\|4(1)\| | * | 100.0%[10/10] | 3.1 |
| Cluster529 | 2(2)\|3(5)\|4(3)\| | * | 100.0%[10/10] | 3.1 |
| Cluster393 | 3(9)\|4(1)\| | * | 100.0%[10/10] | 3.1 |
| Cluster196 | 3(9)\|4(1)\| | * | 100.0%[10/10] | 3.1 |

| | | | | |
|---|---|---|---|---|
| Cluster165 | 3(9)\|4(1)\| | * | 100.0%[10/10] | 3.1 |
| Cluster96 | 3(10)\| | * | 100.0%[10/10] | 3 |
| Cluster63 | 3(10)\| | * | 100.0%[10/10] | 3 |
| Cluster58 | 3(10)\| | * | 100.0%[10/10] | 3 |
| Cluster540 | 3(10)\| | * | 100.0%[10/10] | 3 |
| Cluster539 | 3(10)\| | * | 100.0%[10/10] | 3 |
| Cluster535 | 2(1)\|3(8)\|4(1)\| | * | 100.0%[10/10] | 3 |
| Cluster482 | 3(10)\| | * | 100.0%[10/10] | 3 |
| Cluster47 | 3(10)\| | * | 100.0%[10/10] | 3 |
| Cluster456 | 3(10)\| | * | 100.0%[10/10] | 3 |
| Cluster455 | 3(10)\| | * | 100.0%[10/10] | 3 |
| Cluster451 | 3(10)\| | * | 100.0%[10/10] | 3 |
| Cluster449 | 3(10)\| | * | 100.0%[10/10] | 3 |
| Cluster439 | 3(10)\| | * | 100.0%[10/10] | 3 |
| Cluster42 | 3(10)\| | * | 100.0%[10/10] | 3 |
| Cluster415 | 3(10)\| | * | 100.0%[10/10] | 3 |
| Cluster404 | 3(10)\| | * | 100.0%[10/10] | 3 |
| Cluster39 | 3(10)\| | * | 100.0%[10/10] | 3 |
| Cluster383 | 3(10)\| | * | 100.0%[10/10] | 3 |
| Cluster382 | 3(10)\| | * | 100.0%[10/10] | 3 |
| Cluster376 | 3(10)\| | * | 100.0%[10/10] | 3 |
| Cluster361 | 3(10)\| | * | 100.0%[10/10] | 3 |
| Cluster358 | 3(10)\| | * | 100.0%[10/10] | 3 |
| Cluster348 | 3(10)\| | * | 100.0%[10/10] | 3 |
| Cluster335 | 3(10)\| | * | 100.0%[10/10] | 3 |

| | | | | |
|---|---|---|---|---|
| Cluster333 | 3(10)\| | * | 100.0%[10/10] | 3 |
| Cluster332 | 3(10)\| | * | 100.0%[10/10] | 3 |
| Cluster329 | 3(10)\| | * | 100.0%[10/10] | 3 |
| Cluster265 | 2(2)\|3(6)\|4(2)\| | * | 100.0%[10/10] | 3 |
| Cluster260 | 3(10)\| | * | 100.0%[10/10] | 3 |
| Cluster243 | 3(10)\| | * | 100.0%[10/10] | 3 |
| Cluster227 | 3(10)\| | * | 100.0%[10/10] | 3 |
| Cluster223 | 3(10)\| | * | 100.0%[10/10] | 3 |
| Cluster214 | 3(10)\| | * | 100.0%[10/10] | 3 |
| Cluster212 | 3(10)\| | * | 100.0%[10/10] | 3 |
| Cluster207 | 3(10)\| | * | 100.0%[10/10] | 3 |
| Cluster193 | 3(10)\| | * | 100.0%[10/10] | 3 |
| Cluster170 | 3(10)\| | * | 100.0%[10/10] | 3 |
| Cluster163 | 3(10)\| | * | 100.0%[10/10] | 3 |
| Cluster145 | 3(10)\| | * | 100.0%[10/10] | 3" |

**Supplementary Table5.**

**Cluster sets of identified undiscovered multicopy regions from Streptococcus agalactiae.**

| ClusterID | Distribution | Marker | Percent_of_strains | Weighted_Average_Copy |
|---|---|---|---|---|
| Cluster99 | 0(1)\|1(2)\|3(2)\|4(1)\|7(1)\| | * | 85.7%[6/7] | 2.714285714 |
| Cluster89 | 0(1)\|1(2)\|3(2)\|4(1)\|7(1)\| | * | 85.7%[6/7] | 2.714285714 |
| Cluster87 | 0(1)\|1(2)\|3(2)\|4(1)\|7(1)\| | * | 85.7%[6/7] | 2.714285714 |
| Cluster86 | 0(1)\|1(2)\|3(2)\|4(1)\|7(1)\| | * | 85.7%[6/7] | 2.714285714 |
| Cluster81 | 0(1)\|1(2)\|3(2)\|4(1)\|7(1)\| | * | 85.7%[6/7] | 2.714285714 |
| Cluster80 | 0(1)\|1(2)\|3(2)\|4(1)\|7(1)\| | * | 85.7%[6/7] | 2.714285714 |
| Cluster74 | 0(1)\|1(2)\|3(2)\|4(1)\|7(1)\| | * | 85.7%[6/7] | 2.714285714 |
| Cluster7 | 0(1)\|1(2)\|3(2)\|4(1)\|7(1)\| | * | 85.7%[6/7] | 2.714285714 |
| Cluster68 | 0(1)\|1(2)\|3(2)\|4(1)\|7(1)\| | * | 85.7%[6/7] | 2.714285714 |
| Cluster19 | 0(1)\|1(2)\|3(2)\|4(1)\|7(1)\| | * | 85.7%[6/7] | 2.714285714 |
| Cluster67 | 0(1)\|1(2)\|3(3)\|4(1)\| | * | 85.7%[6/7] | 2.142857143 |
| Cluster53 | 0(1)\|1(2)\|3(3)\|4(1)\| | * | 85.7%[6/7] | 2.142857143 |

**Supplementary Table6.**

**Cluster sets of identified undiscovered multicopy regions from H. pylori UA802, H. pylori strain PMSS1, H. pylori strain 7.13.**

| ClusterID | Distribution | Marker | Percent_of_strains | Weighted_Average_Copy |
|---|---|---|---|---|
| Cluster8 | 1(3)\|2(8)\|3(6)\|4(3)\|5(1)\| | * | 100.0%[21/21] | 2.571428571 |
| Cluster107 | 1(6)\|2(9)\|3(6)\| | * | 100.0%[21/21] | 2 |
| Cluster66 | 1(5)\|2(15)\|3(1)\| | * | 100.0%[21/21] | 1.80952381 |
| Cluster24 | 1(6)\|2(13)\|3(2)\| | * | 100.0%[21/21] | 1.80952381 |
| Cluster42 | 0(3)\|1(2)\|2(13)\|3(3)\| | * | 85.7%[18/21] | 1.761904762 |
| Cluster299 | 0(2)\|1(4)\|2(12)\|3(3)\| | * | 90.5%[19/21] | 1.761904762 |
| Cluster133 | 0(4)\|1(3)\|2(11)\|3(3)\| | * | 81.0%[17/21] | 1.619047619 |
| Cluster55 | 1(13)\|2(5)\|3(3)\| | * | 100.0%[21/21] | 1.523809524 |
| Cluster40 | 0(4)\|1(5)\|2(9)\|3(3)\| | * | 81.0%[17/21] | 1.523809524 |
| Cluster262 | 1(10)\|2(11)\| | * | 100.0%[21/21] | 1.523809524 |
| Cluster236 | 0(4)\|1(7)\|2(6)\|3(4)\| | * | 81.0%[17/21] | 1.476190476 |
| Cluster109 | 0(4)\|1(5)\|2(11)\|3(1)\| | * | 81.0%[17/21] | 1.428571429 |
| Cluster333 | 0(1)\|1(13)\|2(7)\| | * | 95.2%[20/21] | 1.285714286 |
| Cluster3 | 0(3)\|1(11)\|2(5)\|3(2)\| | * | 85.7%[18/21] | 1.285714286 |
| Cluster211 | 0(4)\|1(10)\|2(4)\|3(3)\| | * | 81.0%[17/21] | 1.285714286 |
| Cluster158 | 0(3)\|1(12)\|2(3)\|3(3)\| | * | 85.7%[18/21] | 1.285714286 |
| Cluster354 | 0(1)\|1(14)\|2(6)\| | * | 95.2%[20/21] | 1.238095238 |
| Cluster25 | 0(4)\|1(9)\|2(7)\|3(1)\| | * | 81.0%[17/21] | 1.238095238 |
| Cluster214 | 1(17)\|2(4)\| | * | 100.0%[21/21] | 1.19047619 |
| Cluster2 | 0(4)\|1(10)\|2(6)\|3(1)\| | * | 81.0%[17/21] | 1.19047619 |

| | | | | |
|---|---|---|---|---|
| Cluster20 | 0(4)\|1(10)\|2(6)\|3(1)\| | * | 81.0%[17/21] | 1.19047619 |
| Cluster150 | 0(3)\|1(11)\|2(7)\| | * | 85.7%[18/21] | 1.19047619 |
| Cluster41 | 0(1)\|1(16)\|2(4)\| | * | 95.2%[20/21] | 1.142857143 |
| Cluster355 | 0(4)\|1(11)\|2(5)\|3(1)\| | * | 81.0%[17/21] | 1.142857143 |
| Cluster32 | 0(3)\|1(13)\|2(4)\|3(1)\| | * | 85.7%[18/21] | 1.142857143 |
| Cluster250 | 1(18)\|2(3)\| | * | 100.0%[21/21] | 1.142857143 |
| Cluster227 | 1(18)\|2(3)\| | * | 100.0%[21/21] | 1.142857143 |
| Cluster201 | 1(19)\|2(1)\|3(1)\| | * | 100.0%[21/21] | 1.142857143 |
| Cluster141 | 0(3)\|1(12)\|2(6)\| | * | 85.7%[18/21] | 1.142857143 |
| Cluster62 | 1(19)\|2(2)\| | * | 100.0%[21/21] | 1.095238095 |
| Cluster53 | 0(4)\|1(12)\|2(4)\|3(1)\| | * | 81.0%[17/21] | 1.095238095 |
| Cluster373 | 0(3)\|1(13)\|2(5)\| | * | 85.7%[18/21] | 1.095238095 |
| Cluster350 | 0(4)\|1(12)\|2(4)\|3(1)\| | * | 81.0%[17/21] | 1.095238095 |
| Cluster287 | 0(3)\|1(13)\|2(5)\| | * | 85.7%[18/21] | 1.095238095 |
| Cluster216 | 1(19)\|2(2)\| | * | 100.0%[21/21] | 1.095238095 |
| Cluster205 | 0(4)\|1(12)\|2(4)\|3(1)\| | * | 81.0%[17/21] | 1.095238095 |
| Cluster200 | 0(4)\|1(12)\|2(4)\|3(1)\| | * | 81.0%[17/21] | 1.095238095 |
| Cluster19 | 0(4)\|1(12)\|2(4)\|3(1)\| | * | 81.0%[17/21] | 1.095238095 |
| Cluster155 | 1(19)\|2(2)\| | * | 100.0%[21/21] | 1.095238095 |
| Cluster135 | 0(1)\|1(17)\|2(3)\| | * | 95.2%[20/21] | 1.095238095 |
| Cluster134 | 0(3)\|1(13)\|2(5)\| | * | 85.7%[18/21] | 1.095238095 |
| Cluster117 | 0(1)\|1(18)\|2(1)\|3(1)\| | * | 95.2%[20/21] | 1.095238095 |
| Cluster95 | 1(20)\|2(1)\| | * | 100.0%[21/21] | 1.047619048 |
| Cluster85 | 1(20)\|2(1)\| | * | 100.0%[21/21] | 1.047619048 |
| Cluster84 | 1(20)\|2(1)\| | * | 100.0%[21/21] | 1.047619048 |

| | | | | |
|---|---|---|---|---|
| Cluster83 | 1(20)\|2(1)\| | * | 100.0%[21/21] | 1.047619048 |
| Cluster81 | 1(20)\|2(1)\| | * | 100.0%[21/21] | 1.047619048 |
| Cluster77 | 1(20)\|2(1)\| | * | 100.0%[21/21] | 1.047619048 |
| Cluster72 | 1(20)\|2(1)\| | * | 100.0%[21/21] | 1.047619048 |
| Cluster68 | 0(4)\|1(12)\|2(5)\| | * | 81.0%[17/21] | 1.047619048 |
| Cluster6 | 1(20)\|2(1)\| | * | 100.0%[21/21] | 1.047619048 |
| Cluster5 | 1(20)\|2(1)\| | * | 100.0%[21/21] | 1.047619048 |
| Cluster44 | 0(2)\|1(16)\|2(3)\| | * | 90.5%[19/21] | 1.047619048 |
| Cluster4 | 1(20)\|2(1)\| | * | 100.0%[21/21] | 1.047619048 |
| Cluster374 | 1(20)\|2(1)\| | * | 100.0%[21/21] | 1.047619048 |
| Cluster341 | 1(20)\|2(1)\| | * | 100.0%[21/21] | 1.047619048 |
| Cluster338 | 1(20)\|2(1)\| | * | 100.0%[21/21] | 1.047619048 |
| Cluster316 | 0(2)\|1(16)\|2(3)\| | * | 90.5%[19/21] | 1.047619048 |
| Cluster313 | 1(20)\|2(1)\| | * | 100.0%[21/21] | 1.047619048 |
| Cluster310 | 1(20)\|2(1)\| | * | 100.0%[21/21] | 1.047619048 |
| Cluster309 | 1(20)\|2(1)\| | * | 100.0%[21/21] | 1.047619048 |
| Cluster306 | 0(1)\|1(18)\|2(2)\| | * | 95.2%[20/21] | 1.047619048 |
| Cluster298 | 1(20)\|2(1)\| | * | 100.0%[21/21] | 1.047619048 |
| Cluster292 | 1(20)\|2(1)\| | * | 100.0%[21/21] | 1.047619048 |
| Cluster283 | 1(20)\|2(1)\| | * | 100.0%[21/21] | 1.047619048 |
| Cluster258 | 1(20)\|2(1)\| | * | 100.0%[21/21] | 1.047619048 |
| Cluster254 | 1(20)\|2(1)\| | * | 100.0%[21/21] | 1.047619048 |
| Cluster253 | 1(20)\|2(1)\| | * | 100.0%[21/21] | 1.047619048 |
| Cluster204 | 1(20)\|2(1)\| | * | 100.0%[21/21] | 1.047619048 |
| Cluster202 | 1(20)\|2(1)\| | * | 100.0%[21/21] | 1.047619048 |

| | | | | |
|---|---|---|---|---|
| Cluster198 | 1(20)\|2(1)\| | * | 100.0%[21/21] | 1.047619048 |
| Cluster185 | 1(20)\|2(1)\| | * | 100.0%[21/21] | 1.047619048 |
| Cluster179 | 1(20)\|2(1)\| | * | 100.0%[21/21] | 1.047619048 |
| Cluster177 | 0(4)\|1(12)\|2(5)\| | * | 81.0%[17/21] | 1.047619048 |
| Cluster149 | 1(20)\|2(1)\| | * | 100.0%[21/21] | 1.047619048 |
| Cluster143 | 1(20)\|2(1)\| | * | 100.0%[21/21] | 1.047619048 |
| Cluster11 | 1(20)\|2(1)\| | * | 100.0%[21/21] | 1.047619048 |
| Cluster70 | 1(21)\| | * | 100.0%[21/21] | 1 |
| Cluster375 | 1(21)\| | * | 100.0%[21/21] | 1 |
| Cluster301 | 1(21)\| | * | 100.0%[21/21] | 1 |
| Cluster284 | 0(1)\|1(19)\|2(1)\| | * | 95.2%[20/21] | 1 |
| Cluster273 | 1(21)\| | * | 100.0%[21/21] | 1 |
| Cluster256 | 0(1)\|1(19)\|2(1)\| | * | 95.2%[20/21] | 1 |
| Cluster223 | 0(1)\|1(19)\|2(1)\| | * | 95.2%[20/21] | 1 |
| Cluster212 | 0(1)\|1(19)\|2(1)\| | * | 95.2%[20/21] | 1 |
| Cluster146 | 1(21)\| | * | 100.0%[21/21] | 1 |
| Cluster98 | 0(2)\|1(18)\|2(1)\| | * | 90.5%[19/21] | 0.952380952 |
| Cluster45 | 0(4)\|1(15)\|2(1)\|3(1)\| | * | 81.0%[17/21] | 0.952380952 |
| Cluster362 | 0(2)\|1(18)\|2(1)\| | * | 90.5%[19/21] | 0.952380952 |
| Cluster359 | 0(2)\|1(18)\|2(1)\| | * | 90.5%[19/21] | 0.952380952 |
| Cluster344 | 0(1)\|1(20)\| | * | 95.2%[20/21] | 0.952380952 |
| Cluster339 | 0(2)\|1(18)\|2(1)\| | * | 90.5%[19/21] | 0.952380952 |
| Cluster336 | 0(4)\|1(14)\|2(3)\| | * | 81.0%[17/21] | 0.952380952 |
| Cluster286 | 0(1)\|1(20)\| | * | 95.2%[20/21] | 0.952380952 |
| Cluster249 | 0(4)\|1(14)\|2(3)\| | * | 81.0%[17/21] | 0.952380952 |

| | | | | |
|---|---|---|---|---|
| Cluster240 | 0(1)\|1(20)\| | * | 95.2%[20/21] | 0.952380952 |
| Cluster232 | 0(1)\|1(20)\| | * | 95.2%[20/21] | 0.952380952 |
| Cluster187 | 0(1)\|1(20)\| | * | 95.2%[20/21] | 0.952380952 |
| Cluster144 | 0(2)\|1(18)\|2(1)\| | * | 90.5%[19/21] | 0.952380952 |
| Cluster129 | 0(1)\|1(20)\| | * | 95.2%[20/21] | 0.952380952 |
| Cluster100 | 0(2)\|1(18)\|2(1)\| | * | 90.5%[19/21] | 0.952380952 |
| Cluster57 | 0(4)\|1(15)\|2(2)\| | * | 81.0%[17/21] | 0.904761905 |
| Cluster377 | 0(3)\|1(17)\|2(1)\| | * | 85.7%[18/21] | 0.904761905 |
| Cluster308 | 0(3)\|1(17)\|2(1)\| | * | 85.7%[18/21] | 0.904761905 |
| Cluster293 | 0(4)\|1(15)\|2(2)\| | * | 81.0%[17/21] | 0.904761905 |
| Cluster243 | 0(3)\|1(17)\|2(1)\| | * | 85.7%[18/21] | 0.904761905 |
| Cluster191 | 0(4)\|1(16)\|3(1)\| | * | 81.0%[17/21] | 0.904761905 |
| Cluster170 | 0(4)\|1(15)\|2(2)\| | * | 81.0%[17/21] | 0.904761905 |
| Cluster108 | 0(3)\|1(17)\|2(1)\| | * | 85.7%[18/21] | 0.904761905 |
| Cluster378 | 0(3)\|1(18)\| | * | 85.7%[18/21] | 0.857142857 |
| Cluster285 | 0(4)\|1(16)\|2(1)\| | * | 81.0%[17/21] | 0.857142857 |
| Cluster241 | 0(3)\|1(18)\| | * | 85.7%[18/21] | 0.857142857 |
| Cluster221 | 0(4)\|1(16)\|2(1)\| | * | 81.0%[17/21] | 0.857142857 |
| Cluster219 | 0(4)\|1(16)\|2(1)\| | * | 81.0%[17/21] | 0.857142857 |
| Cluster151 | 0(4)\|1(16)\|2(1)\| | * | 81.0%[17/21] | 0.857142857 |
| Cluster99 | 0(16)\|1(2)\|2(3)\| | | | 23.8%[5/21] |
| Cluster97 | 0(17)\|1(3)\|2(1)\| | | | 19.0%[4/21] |
| Cluster96 | 0(20)\|2(1)\| | | | 4.8%[1/21] |
| Cluster94 | 0(17)\|1(3)\|2(1)\| | | | 19.0%[4/21] |
| Cluster93 | 0(20)\|2(1)\| | | | 4.8%[1/21] |

| | | | | |
|---|---|---|---|---|
| Cluster92 | 0(20)\|3(1)\| | | | 4.8%[1/21] |
| Cluster91 | 0(20)\|3(1)\| | | | 4.8%[1/21] |
| Cluster9 | 0(8)\|1(12)\|2(1)\| | | | 61.9%[13/21] |
| Cluster90 | 0(9)\|1(11)\|2(1)\| | | | 57.1%[12/21] |
| Cluster89 | 0(15)\|1(3)\|4(2)\|5(1)\| | | | 28.6%[6/21] |
| Cluster88 | 0(20)\|2(1)\| | | | 4.8%[1/21] |
| Cluster87 | 0(20)\|5(1)\| | | | 4.8%[1/21] |
| Cluster86 | 0(17)\|1(3)\|2(1)\| | | | 19.0%[4/21] |
| Cluster82 | 0(18)\|1(2)\|2(1)\| | | | 14.3%[3/21] |
| Cluster80 | 0(5)\|1(15)\|2(1)\| | | | 76.2%[16/21] |
| Cluster79 | 0(6)\|1(12)\|2(2)\|3(1)\| | | | 71.4%[15/21] |
| Cluster78 | 0(14)\|1(6)\|2(1)\| | | | 33.3%[7/21] |
| Cluster76 | 0(12)\|1(8)\|2(1)\| | | | 42.9%[9/21] |
| Cluster75 | 0(20)\|2(1)\| | | | 4.8%[1/21] |
| Cluster74 | 0(16)\|1(5)\| | | | 23.8%[5/21] |
| Cluster73 | 0(19)\|1(2)\| | | | 9.5%[2/21] |
| Cluster71 | 0(11)\|1(5)\|2(5)\| | | | 47.6%[10/21] |
| Cluster7 | 0(20)\|2(1)\| | | | 4.8%[1/21] |
| Cluster69 | 0(14)\|1(4)\|2(3)\| | | | 33.3%[7/21] |
| Cluster67 | 0(14)\|1(7)\| | | | 33.3%[7/21] |
| Cluster65 | 0(13)\|1(7)\|2(1)\| | | | 38.1%[8/21] |
| Cluster64 | 0(5)\|1(14)\|2(2)\| | | | 76.2%[16/21] |
| Cluster63 | 0(5)\|1(12)\|2(4)\| | | | 76.2%[16/21] |
| Cluster61 | 0(18)\|1(3)\| | | | 14.3%[3/21] |
| Cluster60 | 0(19)\|1(1)\|2(1)\| | | | 9.5%[2/21] |

| | | | | |
|---|---|---|---|---|
| Cluster59 | 0(11)\|1(9)\|2(1)\| | | | 47.6%[10/21] |
| Cluster58 | 0(5)\|1(12)\|2(4)\| | | | 76.2%[16/21] |
| Cluster56 | 0(18)\|1(2)\|3(1)\| | | | 14.3%[3/21] |
| Cluster54 | 0(5)\|1(5)\|2(9)\|3(2)\| | | | 76.2%[16/21] |
| Cluster52 | 0(18)\|1(2)\|2(1)\| | | | 14.3%[3/21] |
| Cluster51 | 0(20)\|2(1)\| | | | 4.8%[1/21] |
| Cluster50 | 0(21)\| | | | 0.0%[0/21] |
| Cluster49 | 0(5)\|1(8)\|2(5)\|3(3)\| | | | 76.2%[16/21] |
| Cluster48 | 0(9)\|1(5)\|2(3)\|3(4)\| | | | 57.1%[12/21] |
| Cluster47 | 0(20)\|1(1)\| | | | 4.8%[1/21] |
| Cluster46 | 0(20)\|3(1)\| | | | 4.8%[1/21] |
| Cluster43 | 0(20)\|1(1)\| | | | 4.8%[1/21] |
| Cluster39 | 0(15)\|1(5)\|2(1)\| | | | 28.6%[6/21] |
| Cluster38 | 0(15)\|1(4)\|2(1)\|3(1)\| | | | 28.6%[6/21] |
| Cluster380 | 0(5)\|1(6)\|2(7)\|3(1)\|4(2)\| | | | 76.2%[16/21] |
| Cluster379 | 0(17)\|1(4)\| | | | 19.0%[4/21] |
| Cluster376 | 0(8)\|1(11)\|2(2)\| | | | 61.9%[13/21] |
| Cluster372 | 0(16)\|1(5)\| | | | 23.8%[5/21] |
| Cluster371 | 0(19)\|1(1)\|3(1)\| | | | 9.5%[2/21] |
| Cluster37 | 0(13)\|1(8)\| | | | 38.1%[8/21] |
| Cluster370 | 0(12)\|1(8)\|2(1)\| | | | 42.9%[9/21] |
| Cluster369 | 0(19)\|1(1)\|2(1)\| | | | 9.5%[2/21] |
| Cluster368 | 0(20)\|2(1)\| | | | 4.8%[1/21] |
| Cluster367 | 0(18)\|1(2)\|2(1)\| | | | 14.3%[3/21] |
| Cluster366 | 0(20)\|2(1)\| | | | 4.8%[1/21] |

| | | | | |
|---|---|---|---|---|
| Cluster365 | 0(16)\|1(5)\| | | | 23.8%[5/21] |
| Cluster364 | 0(18)\|1(1)\|2(2)\| | | | 14.3%[3/21] |
| Cluster363 | 0(5)\|1(16)\| | | | 76.2%[16/21] |
| Cluster361 | 0(12)\|1(1)\|2(6)\|4(2)\| | | | 42.9%[9/21] |
| Cluster36 | 0(20)\|1(1)\| | | | 4.8%[1/21] |
| Cluster360 | 0(20)\|3(1)\| | | | 4.8%[1/21] |
| Cluster358 | 0(19)\|2(2)\| | | | 9.5%[2/21] |
| Cluster357 | 0(19)\|1(2)\| | | | 9.5%[2/21] |
| Cluster356 | 0(12)\|1(8)\|2(1)\| | | | 42.9%[9/21] |
| Cluster353 | 0(20)\|2(1)\| | | | 4.8%[1/21] |
| Cluster352 | 0(18)\|1(2)\|2(1)\| | | | 14.3%[3/21] |
| Cluster351 | 0(9)\|1(9)\|2(3)\| | | | 57.1%[12/21] |
| Cluster35 | 0(20)\|2(1)\| | | | 4.8%[1/21] |
| Cluster349 | 0(20)\|2(1)\| | | | 4.8%[1/21] |
| Cluster348 | 0(5)\|1(15)\|2(1)\| | | | 76.2%[16/21] |
| Cluster347 | 0(19)\|1(1)\|2(1)\| | | | 9.5%[2/21] |
| Cluster346 | 0(10)\|1(10)\|2(1)\| | | | 52.4%[11/21] |
| Cluster345 | 0(20)\|2(1)\| | | | 4.8%[1/21] |
| Cluster343 | 0(8)\|1(12)\|2(1)\| | | | 61.9%[13/21] |
| Cluster342 | 0(8)\|1(12)\|2(1)\| | | | 61.9%[13/21] |
| Cluster34 | 0(8)\|1(6)\|2(7)\| | | | 61.9%[13/21] |
| Cluster340 | 0(20)\|2(1)\| | | | 4.8%[1/21] |
| Cluster337 | 0(12)\|1(7)\|2(2)\| | | | 42.9%[9/21] |
| Cluster335 | 0(17)\|1(3)\|2(1)\| | | | 19.0%[4/21] |
| Cluster334 | 0(20)\|3(1)\| | | | 4.8%[1/21] |

| | | | | |
|---|---|---|---|---|
| Cluster332 | 0(19)\|1(1)\|2(1)\| | | | 9.5%[2/21] |
| Cluster331 | 0(7)\|1(13)\|2(1)\| | | | 66.7%[14/21] |
| Cluster33 | 0(10)\|1(8)\|2(3)\| | | | 52.4%[11/21] |
| Cluster330 | 0(15)\|1(2)\|2(3)\|3(1)\| | | | 28.6%[6/21] |
| Cluster329 | 0(16)\|1(4)\|2(1)\| | | | 23.8%[5/21] |
| Cluster328 | 0(17)\|1(4)\| | | | 19.0%[4/21] |
| Cluster327 | 0(20)\|1(1)\| | | | 4.8%[1/21] |
| Cluster326 | 0(18)\|1(2)\|2(1)\| | | | 14.3%[3/21] |
| Cluster325 | 0(12)\|1(9)\| | | | 42.9%[9/21] |
| Cluster324 | 0(20)\|3(1)\| | | | 4.8%[1/21] |
| Cluster323 | 0(16)\|1(4)\|2(1)\| | | | 23.8%[5/21] |
| Cluster322 | 0(13)\|1(5)\|2(3)\| | | | 38.1%[8/21] |
| Cluster321 | 0(17)\|1(4)\| | | | 19.0%[4/21] |
| Cluster320 | 0(15)\|1(2)\|2(1)\|3(1)\|4(1)\|5(1)\| | | | 28.6%[6/21] |
| Cluster319 | 0(20)\|4(1)\| | | | 4.8%[1/21] |
| Cluster318 | 0(19)\|1(1)\|2(1)\| | | | 9.5%[2/21] |
| Cluster317 | 0(19)\|1(1)\|2(1)\| | | | 9.5%[2/21] |
| Cluster315 | 0(19)\|1(1)\|2(1)\| | | | 9.5%[2/21] |
| Cluster314 | 0(7)\|1(13)\|2(1)\| | | | 66.7%[14/21] |
| Cluster312 | 0(8)\|1(12)\|2(1)\| | | | 61.9%[13/21] |
| Cluster311 | 0(19)\|1(1)\|2(1)\| | | | 9.5%[2/21] |
| Cluster31 | 0(11)\|1(9)\|2(1)\| | | | 47.6%[10/21] |
| Cluster307 | 0(5)\|1(15)\|2(1)\| | | | 76.2%[16/21] |
| Cluster305 | 0(9)\|1(8)\|2(3)\|3(1)\| | | | 57.1%[12/21] |
| Cluster304 | 0(16)\|1(4)\|2(1)\| | | | 23.8%[5/21] |

| | | | | |
|---|---|---|---|---|
| Cluster303 | 0(7)\|1(14)\| | | | 66.7%[14/21] |
| Cluster302 | 0(10)\|1(9)\|2(2)\| | | | 52.4%[11/21] |
| Cluster30 | 0(18)\|3(1)\|4(1)\|5(1)\| | | | 14.3%[3/21] |
| Cluster300 | 0(20)\|1(1)\| | | | 4.8%[1/21] |
| Cluster297 | 0(15)\|1(3)\|3(1)\|4(1)\|5(1)\| | | | 28.6%[6/21] |
| Cluster296 | 0(18)\|1(2)\|2(1)\| | | | 14.3%[3/21] |
| Cluster295 | 0(19)\|2(2)\| | | | 9.5%[2/21] |
| Cluster294 | 0(7)\|1(13)\|2(1)\| | | | 66.7%[14/21] |
| Cluster291 | 0(20)\|3(1)\| | | | 4.8%[1/21] |
| Cluster29 | 0(20)\|2(1)\| | | | 4.8%[1/21] |
| Cluster290 | 0(21)\| | | | 0.0%[0/21] |
| Cluster289 | 0(20)\|3(1)\| | | | 4.8%[1/21] |
| Cluster288 | 0(16)\|1(2)\|2(3)\| | | | 23.8%[5/21] |
| Cluster282 | 0(11)\|1(9)\|2(1)\| | | | 47.6%[10/21] |
| Cluster281 | 0(9)\|1(11)\|2(1)\| | | | 57.1%[12/21] |
| Cluster28 | 0(10)\|1(10)\|2(1)\| | | | 52.4%[11/21] |
| Cluster280 | 0(5)\|1(11)\|2(5)\| | | | 76.2%[16/21] |
| Cluster279 | 0(5)\|1(6)\|2(7)\|3(2)\|4(1)\| | | | 76.2%[16/21] |
| Cluster278 | 0(16)\|1(5)\| | | | 23.8%[5/21] |
| Cluster277 | 0(16)\|1(4)\|2(1)\| | | | 23.8%[5/21] |
| Cluster276 | 0(11)\|1(9)\|2(1)\| | | | 47.6%[10/21] |
| Cluster275 | 0(13)\|1(8)\| | | | 38.1%[8/21] |
| Cluster274 | 0(16)\|1(4)\|2(1)\| | | | 23.8%[5/21] |
| Cluster272 | 0(5)\|1(13)\|2(3)\| | | | 76.2%[16/21] |
| Cluster271 | 0(21)\| | | | 0.0%[0/21] |

| | | | | |
|---|---|---|---|---|
| Cluster27 | 0(5)\|1(11)\|2(4)\|3(1)\| | | | 76.2%[16/21] |
| Cluster270 | 0(14)\|1(4)\|2(3)\| | | | 33.3%[7/21] |
| Cluster269 | 0(11)\|1(9)\|2(1)\| | | | 47.6%[10/21] |
| Cluster268 | 0(16)\|1(4)\|2(1)\| | | | 23.8%[5/21] |
| Cluster267 | 0(16)\|1(5)\| | | | 23.8%[5/21] |
| Cluster266 | 0(18)\|1(2)\|2(1)\| | | | 14.3%[3/21] |
| Cluster265 | 0(20)\|2(1)\| | | | 4.8%[1/21] |
| Cluster264 | 0(13)\|1(3)\|2(5)\| | | | 38.1%[8/21] |
| Cluster263 | 0(12)\|1(7)\|2(2)\| | | | 42.9%[9/21] |
| Cluster261 | 0(5)\|1(14)\|2(2)\| | | | 76.2%[16/21] |
| Cluster26 | 0(12)\|1(4)\|2(3)\|4(2)\| | | | 42.9%[9/21] |
| Cluster260 | 0(20)\|4(1)\| | | | 4.8%[1/21] |
| Cluster259 | 0(20)\|2(1)\| | | | 4.8%[1/21] |
| Cluster257 | 0(6)\|1(14)\|2(1)\| | | | 71.4%[15/21] |
| Cluster255 | 0(18)\|1(2)\|2(1)\| | | | 14.3%[3/21] |
| Cluster252 | 0(20)\|2(1)\| | | | 4.8%[1/21] |
| Cluster251 | 0(10)\|1(6)\|2(4)\|3(1)\| | | | 52.4%[11/21] |
| Cluster248 | 0(11)\|1(10)\| | | | 47.6%[10/21] |
| Cluster247 | 0(6)\|1(13)\|2(2)\| | | | 71.4%[15/21] |
| Cluster246 | 0(9)\|1(9)\|2(3)\| | | | 57.1%[12/21] |
| Cluster245 | 0(14)\|1(5)\|2(2)\| | | | 33.3%[7/21] |
| Cluster244 | 0(19)\|1(1)\|2(1)\| | | | 9.5%[2/21] |
| Cluster242 | 0(20)\|3(1)\| | | | 4.8%[1/21] |
| Cluster239 | 0(18)\|1(2)\|2(1)\| | | | 14.3%[3/21] |
| Cluster238 | 0(18)\|1(2)\|2(1)\| | | | 14.3%[3/21] |

| | | | | |
|---|---|---|---|---|
| Cluster237 | 0(20)\|2(1)\| | | | 4.8%[1/21] |
| Cluster235 | 0(16)\|1(3)\|2(2)\| | | | 23.8%[5/21] |
| Cluster234 | 0(5)\|1(15)\|2(1)\| | | | 76.2%[16/21] |
| Cluster233 | 0(16)\|1(4)\|2(1)\| | | | 23.8%[5/21] |
| Cluster231 | 0(13)\|1(7)\|3(1)\| | | | 38.1%[8/21] |
| Cluster23 | 0(17)\|1(1)\|2(1)\|4(2)\| | | | 19.0%[4/21] |
| Cluster230 | 0(20)\|2(1)\| | | | 4.8%[1/21] |
| Cluster229 | 0(15)\|1(5)\|2(1)\| | | | 28.6%[6/21] |
| Cluster228 | 0(19)\|1(1)\|5(1)\| | | | 9.5%[2/21] |
| Cluster226 | 0(5)\|1(14)\|2(2)\| | | | 76.2%[16/21] |
| Cluster225 | 0(20)\|5(1)\| | | | 4.8%[1/21] |
| Cluster224 | 0(20)\|1(1)\| | | | 4.8%[1/21] |
| Cluster222 | 0(20)\|2(1)\| | | | 4.8%[1/21] |
| Cluster22 | 0(19)\|2(1)\|3(1)\| | | | 9.5%[2/21] |
| Cluster220 | 0(5)\|1(12)\|2(4)\| | | | 76.2%[16/21] |
| Cluster218 | 0(19)\|2(2)\| | | | 9.5%[2/21] |
| Cluster217 | 0(18)\|1(3)\| | | | 14.3%[3/21] |
| Cluster215 | 0(12)\|1(6)\|2(3)\| | | | 42.9%[9/21] |
| Cluster213 | 0(9)\|1(9)\|2(3)\| | | | 57.1%[12/21] |
| Cluster21 | 0(20)\|2(1)\| | | | 4.8%[1/21] |
| Cluster210 | 0(16)\|1(4)\|2(1)\| | | | 23.8%[5/21] |
| Cluster209 | 0(14)\|1(6)\|2(1)\| | | | 33.3%[7/21] |
| Cluster208 | 0(14)\|1(6)\|2(1)\| | | | 33.3%[7/21] |
| Cluster207 | 0(10)\|1(9)\|2(2)\| | | | 52.4%[11/21] |
| Cluster206 | 0(20)\|3(1)\| | | | 4.8%[1/21] |

| | | | | |
|---|---|---|---|---|
| Cluster203 | 0(19)\|1(2)\| | | | 9.5%[2/21] |
| Cluster199 | 0(20)\|2(1)\| | | | 4.8%[1/21] |
| Cluster197 | 0(20)\|2(1)\| | | | 4.8%[1/21] |
| Cluster196 | 0(8)\|1(10)\|2(3)\| | | | 61.9%[13/21] |
| Cluster195 | 0(8)\|1(9)\|2(4)\| | | | 61.9%[13/21] |
| Cluster194 | 0(8)\|1(11)\|2(2)\| | | | 61.9%[13/21] |
| Cluster193 | 0(20)\|2(1)\| | | | 4.8%[1/21] |
| Cluster192 | 0(8)\|1(11)\|2(2)\| | | | 61.9%[13/21] |
| Cluster190 | 0(18)\|1(2)\|2(1)\| | | | 14.3%[3/21] |
| Cluster189 | 0(16)\|1(4)\|2(1)\| | | | 23.8%[5/21] |
| Cluster188 | 0(20)\|2(1)\| | | | 4.8%[1/21] |
| Cluster186 | 0(18)\|2(1)\|4(1)\|5(1)\| | | | 14.3%[3/21] |
| Cluster184 | 0(9)\|1(7)\|2(5)\| | | | 57.1%[12/21] |
| Cluster183 | 0(14)\|1(6)\|2(1)\| | | | 33.3%[7/21] |
| Cluster182 | 0(20)\|2(1)\| | | | 4.8%[1/21] |
| Cluster181 | 0(18)\|1(2)\|2(1)\| | | | 14.3%[3/21] |
| Cluster18 | 0(20)\|2(1)\| | | | 4.8%[1/21] |
| Cluster180 | 0(11)\|1(9)\|2(1)\| | | | 47.6%[10/21] |
| Cluster178 | 0(9)\|1(11)\|2(1)\| | | | 57.1%[12/21] |
| Cluster176 | 0(7)\|1(12)\|2(2)\| | | | 66.7%[14/21] |
| Cluster175 | 0(5)\|1(12)\|2(4)\| | | | 76.2%[16/21] |
| Cluster174 | 0(16)\|1(4)\|2(1)\| | | | 23.8%[5/21] |
| Cluster173 | 0(12)\|1(7)\|2(2)\| | | | 42.9%[9/21] |
| Cluster172 | 0(11)\|1(9)\|2(1)\| | | | 47.6%[10/21] |
| Cluster171 | 0(6)\|1(14)\|4(1)\| | | | 71.4%[15/21] |

| | | | | |
|---|---|---|---|---|
| Cluster17 | 0(15)\|1(3)\|2(1)\|3(1)\|5(1)\| | | | 28.6%[6/21] |
| Cluster169 | 0(7)\|1(13)\|2(1)\| | | | 66.7%[14/21] |
| Cluster168 | 0(20)\|2(1)\| | | | 4.8%[1/21] |
| Cluster167 | 0(15)\|1(5)\|2(1)\| | | | 28.6%[6/21] |
| Cluster166 | 0(20)\|2(1)\| | | | 4.8%[1/21] |
| Cluster165 | 0(8)\|1(8)\|2(4)\|3(1)\| | | | 61.9%[13/21] |
| Cluster164 | 0(8)\|1(8)\|2(5)\| | | | 61.9%[13/21] |
| Cluster163 | 0(13)\|1(6)\|2(2)\| | | | 38.1%[8/21] |
| Cluster162 | 0(19)\|1(1)\|2(1)\| | | | 9.5%[2/21] |
| Cluster161 | 0(7)\|1(11)\|2(3)\| | | | 66.7%[14/21] |
| Cluster16 | 0(9)\|1(11)\|2(1)\| | | | 57.1%[12/21] |
| Cluster160 | 0(19)\|2(2)\| | | | 9.5%[2/21] |
| Cluster159 | 0(16)\|1(4)\|2(1)\| | | | 23.8%[5/21] |
| Cluster157 | 0(20)\|2(1)\| | | | 4.8%[1/21] |
| Cluster156 | 0(8)\|1(9)\|2(4)\| | | | 61.9%[13/21] |
| Cluster154 | 0(16)\|1(3)\|2(2)\| | | | 23.8%[5/21] |
| Cluster153 | 0(5)\|1(3)\|2(11)\|3(2)\| | | | 76.2%[16/21] |
| Cluster152 | 0(11)\|1(9)\|2(1)\| | | | 47.6%[10/21] |
| Cluster15 | 0(12)\|1(3)\|2(4)\|4(2)\| | | | 42.9%[9/21] |
| Cluster148 | 0(18)\|1(3)\| | | | 14.3%[3/21] |
| Cluster147 | 0(18)\|1(2)\|2(1)\| | | | 14.3%[3/21] |
| Cluster145 | 0(20)\|2(1)\| | | | 4.8%[1/21] |
| Cluster142 | 0(19)\|1(1)\|2(1)\| | | | 9.5%[2/21] |
| Cluster14 | 0(14)\|1(4)\|2(3)\| | | | 33.3%[7/21] |
| Cluster140 | 0(20)\|2(1)\| | | | 4.8%[1/21] |

| | | | | |
|---|---|---|---|---|
| Cluster139 | 0(9)\|1(8)\|2(4)\| | | | 57.1%[12/21] |
| Cluster138 | 0(12)\|1(7)\|2(2)\| | | | 42.9%[9/21] |
| Cluster137 | 0(16)\|1(5)\| | | | 23.8%[5/21] |
| Cluster136 | 0(5)\|1(6)\|2(7)\|3(1)\|4(2)\| | | | 76.2%[16/21] |
| Cluster132 | 0(13)\|1(6)\|2(2)\| | | | 38.1%[8/21] |
| Cluster131 | 0(9)\|1(5)\|2(4)\|3(3)\| | | | 57.1%[12/21] |
| Cluster13 | 0(11)\|1(9)\|2(1)\| | | | 47.6%[10/21] |
| Cluster130 | 0(12)\|1(1)\|2(6)\|3(2)\| | | | 42.9%[9/21] |
| Cluster128 | 0(12)\|1(3)\|2(5)\|3(1)\| | | | 42.9%[9/21] |
| Cluster127 | 0(6)\|1(15)\| | | | 71.4%[15/21] |
| Cluster126 | 0(17)\|1(3)\|3(1)\| | | | 19.0%[4/21] |
| Cluster125 | 0(17)\|1(3)\|3(1)\| | | | 19.0%[4/21] |
| Cluster124 | 0(20)\|3(1)\| | | | 4.8%[1/21] |
| Cluster123 | 0(20)\|3(1)\| | | | 4.8%[1/21] |
| Cluster122 | 0(7)\|1(11)\|2(3)\| | | | 66.7%[14/21] |
| Cluster121 | 0(20)\|3(1)\| | | | 4.8%[1/21] |
| Cluster12 | 0(20)\|2(1)\| | | | 4.8%[1/21] |
| Cluster120 | 0(11)\|1(9)\|2(1)\| | | | 47.6%[10/21] |
| Cluster119 | 0(14)\|1(5)\|2(2)\| | | | 33.3%[7/21] |
| Cluster118 | 0(15)\|1(5)\|2(1)\| | | | 28.6%[6/21] |
| Cluster116 | 0(12)\|1(7)\|2(2)\| | | | 42.9%[9/21] |
| Cluster115 | 0(16)\|1(4)\|2(1)\| | | | 23.8%[5/21] |
| Cluster114 | 0(20)\|2(1)\| | | | 4.8%[1/21] |
| Cluster113 | 0(20)\|2(1)\| | | | 4.8%[1/21] |
| Cluster112 | 0(7)\|1(13)\|2(1)\| | | | 66.7%[14/21] |

| | | | | |
|---|---|---|---|---|
| Cluster111 | 0(16)\|1(4)\|2(1)\| | | | 23.8%[5/21] |
| Cluster110 | 0(20)\|5(1)\| | | | 4.8%[1/21] |
| Cluster106 | 0(5)\|1(4)\|2(11)\|3(1)\| | | | 76.2%[16/21] |
| Cluster105 | 0(14)\|1(6)\|2(1)\| | | | 33.3%[7/21] |
| Cluster104 | 0(18)\|2(1)\|3(2)\| | | | 14.3%[3/21] |
| Cluster103 | 0(16)\|1(2)\|2(3)\| | | | 23.8%[5/21] |
| Cluster102 | 0(12)\|1(1)\|2(6)\|3(1)\|4(1)\| | | | 42.9%[9/21] |
| Cluster1 | 0(13)\|1(7)\|2(1)\| | | | 38.1%[8/21] |
| Cluster101 | 0(16)\|1(3)\|2(2)\| | | | 23.8%[5/21] |
| Cluster10 | 0(12)\|1(8)\|2(1)\| | | | 42.9%[9/21] |
| Cluster0 | 0(10)\|1(10)\|2(1)\| | | | 52.4%[11/21] |

**Supplementary Table7.**

**Cluster sets of identified undiscovered multicopy regions from Legionella pneumophila.**

| ClusterID | Distribution | Marker | Percent_of_strains | Weighted_Average_Copy |
|---|---|---|---|---|
| Cluster66 | 7(1)\|12(1)\|23(1)\|28(1)\|30(4)\|31(1)\| | * | 100.0%[9/9] | 24.55555556 |
| Cluster67 | 5(1)\|12(1)\|21(1)\|27(1)\|28(2)\|29(2)\|30(1)\| | * | 100.0%[9/9] | 23.22222222 |
| Cluster491 | 4(1)\|11(1)\|20(1)\|26(1)\|27(2)\|29(2)\|30(1)\| | * | 100.0%[9/9] | 22.55555556 |
| Cluster65 | 6(1)\|9(1)\|19(1)\|26(2)\|28(4)\| | * | 100.0%[9/9] | 22 |
| Cluster351 | 5(1)\|10(1)\|18(1)\|23(2)\|24(1)\|27(2)\|29(1)\| | * | 100.0%[9/9] | 20.66666667 |
| Cluster180 | 0(1)\|2(1)\|13(1)\|15(1)\|19(1)\|20(1)\|21(2)\|22(1)\| | * | 88.9%[8/9] | 14.77777778 |
| Cluster280 | 4(2)\|6(1)\|8(1)\|9(2)\|11(1)\|12(2)\| | * | 100.0%[9/9] | 8.333333333 |
| Cluster111 | 3(1)\|4(1)\|6(1)\|7(3)\|9(3)\| | * | 100.0%[9/9] | 6.777777778 |
| Cluster253 | 3(2)\|4(2)\|5(5)\| | * | 100.0%[9/9] | 4.333333333 |
| Cluster208 | 1(1)\|2(1)\|3(1)\|4(2)\|5(4)\| | * | 100.0%[9/9] | 3.777777778 |
| Cluster185 | 1(1)\|2(3)\|3(1)\|4(1)\|5(2)\|6(1)\| | * | 100.0%[9/9] | 3.333333333 |
| Cluster250 | 1(2)\|3(4)\|4(2)\|6(1)\| | * | 100.0%[9/9] | 3.111111111 |
| Cluster364 | 0(1)\|2(2)\|3(3)\|4(1)\|5(2)\| | * | 88.9%[8/9] | 3 |
| Cluster100 | 1(3)\|3(5)\|4(1)\| | * | 100.0%[9/9] | 2.444444444 |
| Cluster186 | 1(2)\|2(2)\|3(5)\| | * | 100.0%[9/9] | 2.333333333 |
| Cluster678 | 1(4)\|3(4)\|4(1)\| | * | 100.0%[9/9] | 2.222222222 |
| Cluster31 | 1(3)\|2(1)\|3(5)\| | * | 100.0%[9/9] | 2.222222222 |
| Cluster137 | 1(1)\|2(5)\|3(3)\| | * | 100.0%[9/9] | 2.222222222 |
| Cluster120 | 2(7)\|3(2)\| | * | 100.0%[9/9] | 2.222222222 |
| Cluster60 | 1(2)\|2(4)\|3(3)\| | * | 100.0%[9/9] | 2.111111111 |

| | | | | |
|---|---|---|---|---|
| Cluster54 | 1(2)\|2(4)\|3(3)\| | * | 100.0%[9/9] | 2.111111111 |
| Cluster631 | 1(5)\|3(3)\|4(1)\| | * | 100.0%[9/9] | 2 |
| Cluster533 | 2(9)\| | * | 100.0%[9/9] | 2 |

Supplementary Table8.

Cluster sets of identified undiscovered multicopy regions from Candida auris.

| ClusterID | Distribution | Marker | Percent_of_strains | Weighted_Average_Copy |
|---|---|---|---|---|
| Cluster538 | 6(1)\|7(2)\|10(1)\|13(1)\| | * | 100.0%[5/5] | 8.6 |
| Cluster151 | 6(2)\|7(1)\|8(1)\|12(1)\| | * | 100.0%[5/5] | 7.8 |
| Cluster206 | 2(1)\|7(2)\|9(1)\|12(1)\| | * | 100.0%[5/5] | 7.4 |
| Cluster843 | 1(1)\|2(1)\|3(1)\|10(1)\|20(1)\| | * | 100.0%[5/5] | 7.2 |
| Cluster236 | 0(1)\|2(1)\|4(1)\|10(1)\|20(1)\| | * | 80.0%[4/5] | 7.2 |
| Cluster28 | 4(1)\|5(1)\|6(1)\|9(1)\|11(1)\| | * | 100.0%[5/5] | 7 |
| Cluster16 | 5(1)\|6(2)\|7(1)\|11(1)\| | * | 100.0%[5/5] | 7 |
| Cluster896 | 1(2)\|4(1)\|8(1)\|20(1)\| | * | 100.0%[5/5] | 6.8 |
| Cluster706 | 1(2)\|4(1)\|8(1)\|20(1)\| | * | 100.0%[5/5] | 6.8 |
| Cluster39 | 1(2)\|4(1)\|8(1)\|20(1)\| | * | 100.0%[5/5] | 6.8 |
| Cluster959 | 1(2)\|3(1)\|8(1)\|20(1)\| | * | 100.0%[5/5] | 6.6 |
| Cluster544 | 1(2)\|2(1)\|9(1)\|19(1)\| | * | 100.0%[5/5] | 6.4 |
| Cluster960 | 1(2)\|2(1)\|8(1)\|19(1)\| | * | 100.0%[5/5] | 6.2 |
| Cluster895 | 1(2)\|2(1)\|8(1)\|19(1)\| | * | 100.0%[5/5] | 6.2 |
| Cluster86 | 1(2)\|2(1)\|8(1)\|19(1)\| | * | 100.0%[5/5] | 6.2 |
| Cluster802 | 1(2)\|2(1)\|8(1)\|19(1)\| | * | 100.0%[5/5] | 6.2 |
| Cluster731 | 1(2)\|2(1)\|8(1)\|19(1)\| | * | 100.0%[5/5] | 6.2 |
| Cluster36 | 1(2)\|2(1)\|8(1)\|19(1)\| | * | 100.0%[5/5] | 6.2 |
| Cluster11 | 1(2)\|2(1)\|8(1)\|19(1)\| | * | 100.0%[5/5] | 6.2 |
| Cluster649 | 1(2)\|2(1)\|7(1)\|19(1)\| | * | 100.0%[5/5] | 6 |

| | | | | |
|---|---|---|---|---|
| Cluster569 | 1(2)\|2(1)\|7(1)\|19(1)\| | * | 100.0%[5/5] | 6 |
| Cluster349 | 1(3)\|8(1)\|19(1)\| | * | 100.0%[5/5] | 6 |
| Cluster962 | 0(1)\|1(2)\|8(1)\|19(1)\| | * | 80.0%[4/5] | 5.8 |
| Cluster905 | 1(3)\|7(1)\|19(1)\| | * | 100.0%[5/5] | 5.8 |
| Cluster893 | 1(3)\|7(1)\|19(1)\| | * | 100.0%[5/5] | 5.8 |
| Cluster735 | 1(2)\|2(1)\|6(1)\|19(1)\| | * | 100.0%[5/5] | 5.8 |
| Cluster72 | 1(3)\|7(1)\|19(1)\| | * | 100.0%[5/5] | 5.8 |
| Cluster71 | 1(3)\|7(1)\|19(1)\| | * | 100.0%[5/5] | 5.8 |
| Cluster65 | 1(3)\|7(1)\|19(1)\| | * | 100.0%[5/5] | 5.8 |
| Cluster554 | 1(3)\|7(1)\|19(1)\| | * | 100.0%[5/5] | 5.8 |
| Cluster550 | 1(3)\|7(1)\|19(1)\| | * | 100.0%[5/5] | 5.8 |
| Cluster545 | 1(3)\|7(1)\|19(1)\| | * | 100.0%[5/5] | 5.8 |
| Cluster352 | 1(3)\|7(1)\|19(1)\| | * | 100.0%[5/5] | 5.8 |
| Cluster234 | 1(3)\|7(1)\|19(1)\| | * | 100.0%[5/5] | 5.8 |
| Cluster226 | 1(3)\|7(1)\|19(1)\| | * | 100.0%[5/5] | 5.8 |
| Cluster223 | 1(3)\|7(1)\|19(1)\| | * | 100.0%[5/5] | 5.8 |
| Cluster219 | 1(3)\|7(1)\|19(1)\| | * | 100.0%[5/5] | 5.8 |
| Cluster215 | 1(3)\|7(1)\|19(1)\| | * | 100.0%[5/5] | 5.8 |
| Cluster214 | 1(3)\|7(1)\|19(1)\| | * | 100.0%[5/5] | 5.8 |
| Cluster213 | 1(3)\|7(1)\|19(1)\| | * | 100.0%[5/5] | 5.8 |
| Cluster91 | 0(1)\|1(2)\|7(1)\|19(1)\| | * | 80.0%[4/5] | 5.6 |
| Cluster902 | 1(3)\|7(1)\|18(1)\| | * | 100.0%[5/5] | 5.6 |
| Cluster82 | 1(3)\|7(1)\|18(1)\| | * | 100.0%[5/5] | 5.6 |
| Cluster803 | 0(1)\|1(2)\|8(1)\|18(1)\| | * | 80.0%[4/5] | 5.6 |
| Cluster738 | 1(3)\|6(1)\|19(1)\| | * | 100.0%[5/5] | 5.6 |

| | | | | |
|---|---|---|---|---|
| Cluster454 | 1(2)\|2(1)\|6(1)\|18(1)\| | * | 100.0%[5/5] | 5.6 |
| Cluster377 | 0(1)\|1(2)\|7(1)\|19(1)\| | * | 80.0%[4/5] | 5.6 |
| Cluster360 | 0(1)\|1(2)\|7(1)\|19(1)\| | * | 80.0%[4/5] | 5.6 |
| Cluster356 | 1(3)\|6(1)\|19(1)\| | * | 100.0%[5/5] | 5.6 |
| Cluster354 | 1(2)\|2(1)\|6(1)\|18(1)\| | * | 100.0%[5/5] | 5.6 |
| Cluster350 | 1(3)\|7(1)\|18(1)\| | * | 100.0%[5/5] | 5.6 |
| Cluster211 | 0(1)\|1(2)\|7(1)\|19(1)\| | * | 80.0%[4/5] | 5.6 |
| Cluster210 | 1(3)\|7(1)\|18(1)\| | * | 100.0%[5/5] | 5.6 |
| Cluster801 | 1(3)\|7(1)\|17(1)\| | * | 100.0%[5/5] | 5.4 |
| Cluster73 | 0(1)\|1(1)\|2(1)\|5(1)\|19(1)\| | * | 80.0%[4/5] | 5.4 |
| Cluster521 | 1(2)\|2(1)\|7(1)\|16(1)\| | * | 100.0%[5/5] | 5.4 |
| Cluster452 | 0(1)\|1(2)\|7(1)\|18(1)\| | * | 80.0%[4/5] | 5.4 |
| Cluster353 | 0(1)\|1(2)\|5(1)\|19(1)\| | * | 80.0%[4/5] | 5.2 |
| Cluster208 | 1(1)\|5(1)\|6(2)\|8(1)\| | * | 100.0%[5/5] | 5.2 |
| Cluster165 | 2(1)\|4(2)\|7(1)\|9(1)\| | * | 100.0%[5/5] | 5.2 |
| Cluster967 | 0(1)\|1(2)\|4(1)\|19(1)\| | * | 80.0%[4/5] | 5 |
| Cluster645 | 0(1)\|1(2)\|7(1)\|16(1)\| | * | 80.0%[4/5] | 5 |
| Cluster346 | 0(1)\|1(2)\|4(1)\|17(1)\| | * | 80.0%[4/5] | 4.6 |
| Cluster361 | 0(1)\|1(2)\|6(1)\|14(1)\| | * | 80.0%[4/5] | 4.4 |
| Cluster968 | 0(1)\|1(3)\|18(1)\| | * | 80.0%[4/5] | 4.2 |
| Cluster647 | 0(1)\|1(3)\|18(1)\| | * | 80.0%[4/5] | 4.2 |
| Cluster63 | 0(1)\|1(1)\|2(1)\|4(1)\|14(1)\| | * | 80.0%[4/5] | 4.2 |
| Cluster644 | 0(1)\|1(3)\|17(1)\| | * | 80.0%[4/5] | 4 |
| Cluster87 | 0(1)\|1(2)\|7(1)\|10(1)\| | * | 80.0%[4/5] | 3.8 |
| Cluster772 | 0(1)\|4(2)\|5(1)\|6(1)\| | * | 80.0%[4/5] | 3.8 |

| | | | | |
|---|---|---|---|---|
| Cluster400 | 0(1)\|4(2)\|5(1)\|6(1)\| | * | 80.0%[4/5] | 3.8 |
| Cluster813 | 1(1)\|3(1)\|4(2)\|6(1)\| | * | 100.0%[5/5] | 3.6 |
| Cluster778 | 0(1)\|4(2)\|5(2)\| | * | 80.0%[4/5] | 3.6 |
| Cluster585 | 2(1)\|3(1)\|4(2)\|5(1)\| | * | 100.0%[5/5] | 3.6 |
| Cluster523 | 1(3)\|2(1)\|13(1)\| | * | 100.0%[5/5] | 3.6 |
| Cluster125 | 0(1)\|3(1)\|5(3)\| | * | 80.0%[4/5] | 3.6 |
| Cluster7 | 2(1)\|3(1)\|4(3)\| | * | 100.0%[5/5] | 3.4 |
| Cluster204 | 2(1)\|3(1)\|4(3)\| | * | 100.0%[5/5] | 3.4 |
| Cluster804 | 0(1)\|3(1)\|4(2)\|5(1)\| | * | 80.0%[4/5] | 3.2 |
| Cluster715 | 0(1)\|1(1)\|3(1)\|5(1)\|6(1)\| | * | 80.0%[4/5] | 3 |
| Cluster620 | 0(1)\|2(1)\|4(2)\|5(1)\| | * | 80.0%[4/5] | 3 |
| Cluster414 | 0(1)\|2(1)\|4(2)\|5(1)\| | * | 80.0%[4/5] | 3 |
| Cluster966 | 0(1)\|1(1)\|4(2)\|5(1)\| | * | 80.0%[4/5] | 2.8 |
| Cluster398 | 1(1)\|2(1)\|3(2)\|5(1)\| | * | 100.0%[5/5] | 2.8 |
| Cluster865 | 1(3)\|2(1)\|8(1)\| | * | 100.0%[5/5] | 2.6 |
| Cluster435 | 1(1)\|2(2)\|3(1)\|5(1)\| | * | 100.0%[5/5] | 2.6 |
| Cluster235 | 0(1)\|1(2)\|3(1)\|8(1)\| | * | 80.0%[4/5] | 2.6 |
| Cluster660 | 0(1)\|1(1)\|3(1)\|4(2)\| | * | 80.0%[4/5] | 2.4 |
| Cluster518 | 0(1)\|2(1)\|3(2)\|4(1)\| | * | 80.0%[4/5] | 2.4 |
| Cluster51 | 1(2)\|3(2)\|4(1)\| | * | 100.0%[5/5] | 2.4 |
| Cluster899 | 0(1)\|1(1)\|2(1)\|4(2)\| | * | 80.0%[4/5] | 2.2 |
| Cluster595 | 1(2)\|2(2)\|5(1)\| | * | 100.0%[5/5] | 2.2 |
| Cluster388 | 0(1)\|1(2)\|4(1)\|5(1)\| | * | 80.0%[4/5] | 2.2 |
| Cluster205 | 1(1)\|2(2)\|3(2)\| | * | 100.0%[5/5] | 2.2 |
| Cluster199 | 0(1)\|1(1)\|2(1)\|4(2)\| | * | 80.0%[4/5] | 2.2 |

| | | | | |
|---|---|---|---|---|
| Cluster173 | 1(3)\|3(1)\|5(1)\| | * | 100.0%[5/5] | 2.2 |
| Cluster640 | 0(1)\|1(2)\|3(1)\|5(1)\| | * | 80.0%[4/5] | 2 |
| Cluster52 | 0(1)\|2(3)\|4(1)\| | * | 80.0%[4/5] | 2 |
| Cluster460 | 0(1)\|1(2)\|3(1)\|5(1)\| | * | 80.0%[4/5] | 2 |
| Cluster321 | 0(1)\|1(2)\|3(1)\|5(1)\| | * | 80.0%[4/5] | 2 |
| Cluster104 | 0(1)\|1(2)\|3(1)\|5(1)\| | * | 80.0%[4/5] | 2 |
| Cluster953 | 1(1)\|2(4)\| | * | 100.0%[5/5] | 1.8 |
| Cluster941 | 1(3)\|2(1)\|4(1)\| | * | 100.0%[5/5] | 1.8 |
| Cluster869 | 0(1)\|1(2)\|2(1)\|5(1)\| | * | 80.0%[4/5] | 1.8 |
| Cluster8 | 0(1)\|1(1)\|2(1)\|3(2)\| | * | 80.0%[4/5] | 1.8 |
| Cluster769 | 1(1)\|2(4)\| | * | 100.0%[5/5] | 1.8 |
| Cluster708 | 1(1)\|2(4)\| | * | 100.0%[5/5] | 1.8 |
| Cluster707 | 0(1)\|1(1)\|2(1)\|3(2)\| | * | 80.0%[4/5] | 1.8 |
| Cluster700 | 0(1)\|1(2)\|2(1)\|5(1)\| | * | 80.0%[4/5] | 1.8 |
| Cluster675 | 0(1)\|1(1)\|2(1)\|3(2)\| | * | 80.0%[4/5] | 1.8 |
| Cluster540 | 0(1)\|1(2)\|2(1)\|5(1)\| | * | 80.0%[4/5] | 1.8 |
| Cluster539 | 1(1)\|2(4)\| | * | 100.0%[5/5] | 1.8 |
| Cluster42 | 0(1)\|1(1)\|2(1)\|3(2)\| | * | 80.0%[4/5] | 1.8 |
| Cluster337 | 1(1)\|2(4)\| | * | 100.0%[5/5] | 1.8 |
| Cluster313 | 0(1)\|2(3)\|3(1)\| | * | 80.0%[4/5] | 1.8 |
| Cluster200 | 1(1)\|2(4)\| | * | 100.0%[5/5] | 1.8 |
| Cluster17 | 1(1)\|2(4)\| | * | 100.0%[5/5] | 1.8 |
| Cluster168 | 0(1)\|1(2)\|3(1)\|4(1)\| | * | 80.0%[4/5] | 1.8 |
| Cluster995 | 1(2)\|2(3)\| | * | 100.0%[5/5] | 1.6 |
| Cluster994 | 0(1)\|1(1)\|2(2)\|3(1)\| | * | 80.0%[4/5] | 1.6 |

| | | | | |
|---|---|---|---|---|
| Cluster961 | 1(2)\|2(3)\| | * | 100.0%[5/5] | 1.6 |
| Cluster957 | 0(1)\|1(2)\|2(1)\|4(1)\| | * | 80.0%[4/5] | 1.6 |
| Cluster949 | 1(3)\|2(1)\|3(1)\| | * | 100.0%[5/5] | 1.6 |
| Cluster934 | 1(2)\|2(3)\| | * | 100.0%[5/5] | 1.6 |
| Cluster874 | 1(2)\|2(3)\| | * | 100.0%[5/5] | 1.6 |
| Cluster812 | 1(2)\|2(3)\| | * | 100.0%[5/5] | 1.6 |
| Cluster782 | 1(2)\|2(3)\| | * | 100.0%[5/5] | 1.6 |
| Cluster774 | 1(2)\|2(3)\| | * | 100.0%[5/5] | 1.6 |
| Cluster724 | 1(2)\|2(3)\| | * | 100.0%[5/5] | 1.6 |
| Cluster676 | 0(1)\|1(1)\|2(2)\|3(1)\| | * | 80.0%[4/5] | 1.6 |
| Cluster433 | 1(2)\|2(3)\| | * | 100.0%[5/5] | 1.6 |
| Cluster424 | 1(2)\|2(3)\| | * | 100.0%[5/5] | 1.6 |
| Cluster330 | 0(1)\|1(2)\|3(2)\| | * | 80.0%[4/5] | 1.6 |
| Cluster292 | 1(2)\|2(3)\| | * | 100.0%[5/5] | 1.6 |
| Cluster170 | 1(2)\|2(3)\| | * | 100.0%[5/5] | 1.6 |
| Cluster990 | 1(4)\|3(1)\| | * | 100.0%[5/5] | 1.4 |
| Cluster988 | 1(4)\|3(1)\| | * | 100.0%[5/5] | 1.4 |
| Cluster986 | 1(4)\|3(1)\| | * | 100.0%[5/5] | 1.4 |
| Cluster984 | 1(4)\|3(1)\| | * | 100.0%[5/5] | 1.4 |
| Cluster978 | 1(4)\|3(1)\| | * | 100.0%[5/5] | 1.4 |
| Cluster977 | 1(4)\|3(1)\| | * | 100.0%[5/5] | 1.4 |
| Cluster973 | 1(4)\|3(1)\| | * | 100.0%[5/5] | 1.4 |
| Cluster951 | 0(1)\|1(1)\|2(3)\| | * | 80.0%[4/5] | 1.4 |
| Cluster926 | 1(4)\|3(1)\| | * | 100.0%[5/5] | 1.4 |
| Cluster925 | 1(4)\|3(1)\| | * | 100.0%[5/5] | 1.4 |

| | | | | |
|---|---|---|---|---|
| Cluster924 | 1(4)\|3(1)\| | * | 100.0%[5/5] | 1.4 |
| Cluster919 | 1(4)\|3(1)\| | * | 100.0%[5/5] | 1.4 |
| Cluster917 | 1(4)\|3(1)\| | * | 100.0%[5/5] | 1.4 |
| Cluster910 | 1(4)\|3(1)\| | * | 100.0%[5/5] | 1.4 |
| Cluster909 | 1(4)\|3(1)\| | * | 100.0%[5/5] | 1.4 |
| Cluster901 | 1(4)\|3(1)\| | * | 100.0%[5/5] | 1.4 |
| Cluster878 | 1(3)\|2(2)\| | * | 100.0%[5/5] | 1.4 |
| Cluster875 | 0(1)\|1(2)\|2(1)\|3(1)\| | * | 80.0%[4/5] | 1.4 |
| Cluster847 | 1(4)\|3(1)\| | * | 100.0%[5/5] | 1.4 |
| Cluster846 | 1(4)\|3(1)\| | * | 100.0%[5/5] | 1.4 |
| Cluster844 | 1(4)\|3(1)\| | * | 100.0%[5/5] | 1.4 |
| Cluster836 | 1(4)\|3(1)\| | * | 100.0%[5/5] | 1.4 |
| Cluster832 | 1(4)\|3(1)\| | * | 100.0%[5/5] | 1.4 |
| Cluster831 | 1(4)\|3(1)\| | * | 100.0%[5/5] | 1.4 |
| Cluster830 | 1(4)\|3(1)\| | * | 100.0%[5/5] | 1.4 |
| Cluster828 | 1(4)\|3(1)\| | * | 100.0%[5/5] | 1.4 |
| Cluster827 | 1(4)\|3(1)\| | * | 100.0%[5/5] | 1.4 |
| Cluster824 | 1(4)\|3(1)\| | * | 100.0%[5/5] | 1.4 |
| Cluster821 | 1(4)\|3(1)\| | * | 100.0%[5/5] | 1.4 |
| Cluster811 | 0(1)\|1(1)\|2(3)\| | * | 80.0%[4/5] | 1.4 |
| Cluster786 | 0(1)\|1(2)\|2(1)\|3(1)\| | * | 80.0%[4/5] | 1.4 |
| Cluster760 | 1(4)\|3(1)\| | * | 100.0%[5/5] | 1.4 |
| Cluster756 | 1(4)\|3(1)\| | * | 100.0%[5/5] | 1.4 |
| Cluster754 | 1(4)\|3(1)\| | * | 100.0%[5/5] | 1.4 |
| Cluster749 | 1(4)\|3(1)\| | * | 100.0%[5/5] | 1.4 |

| | | | | |
|---|---|---|---|---|
| Cluster745 | 1(4)\|3(1)\| | * | 100.0%[5/5] | 1.4 |
| Cluster727 | 0(1)\|1(3)\|4(1)\| | * | 80.0%[4/5] | 1.4 |
| Cluster714 | 0(1)\|1(2)\|2(1)\|3(1)\| | * | 80.0%[4/5] | 1.4 |
| Cluster673 | 1(4)\|3(1)\| | * | 100.0%[5/5] | 1.4 |
| Cluster672 | 1(4)\|3(1)\| | * | 100.0%[5/5] | 1.4 |
| Cluster671 | 1(4)\|3(1)\| | * | 100.0%[5/5] | 1.4 |
| Cluster658 | 1(4)\|3(1)\| | * | 100.0%[5/5] | 1.4 |
| Cluster655 | 1(4)\|3(1)\| | * | 100.0%[5/5] | 1.4 |
| Cluster654 | 1(4)\|3(1)\| | * | 100.0%[5/5] | 1.4 |
| Cluster653 | 1(4)\|3(1)\| | * | 100.0%[5/5] | 1.4 |
| Cluster642 | 1(4)\|3(1)\| | * | 100.0%[5/5] | 1.4 |
| Cluster622 | 1(4)\|3(1)\| | * | 100.0%[5/5] | 1.4 |
| Cluster584 | 1(3)\|2(2)\| | * | 100.0%[5/5] | 1.4 |
| Cluster578 | 1(4)\|3(1)\| | * | 100.0%[5/5] | 1.4 |
| Cluster577 | 1(4)\|3(1)\| | * | 100.0%[5/5] | 1.4 |
| Cluster574 | 1(4)\|3(1)\| | * | 100.0%[5/5] | 1.4 |
| Cluster573 | 1(4)\|3(1)\| | * | 100.0%[5/5] | 1.4 |
| Cluster570 | 1(4)\|3(1)\| | * | 100.0%[5/5] | 1.4 |
| Cluster563 | 1(4)\|3(1)\| | * | 100.0%[5/5] | 1.4 |
| Cluster562 | 1(4)\|3(1)\| | * | 100.0%[5/5] | 1.4 |
| Cluster560 | 1(4)\|3(1)\| | * | 100.0%[5/5] | 1.4 |
| Cluster559 | 1(4)\|3(1)\| | * | 100.0%[5/5] | 1.4 |
| Cluster558 | 1(4)\|3(1)\| | * | 100.0%[5/5] | 1.4 |
| Cluster556 | 1(4)\|3(1)\| | * | 100.0%[5/5] | 1.4 |
| Cluster511 | 0(1)\|1(3)\|4(1)\| | * | 80.0%[4/5] | 1.4 |

| | | | | |
|---|---|---|---|---|
| Cluster505 | 0(1)\|1(1)\|2(3)\| | * | 80.0%[4/5] | 1.4 |
| Cluster480 | 1(4)\|3(1)\| | * | 100.0%[5/5] | 1.4 |
| Cluster479 | 1(4)\|3(1)\| | * | 100.0%[5/5] | 1.4 |
| Cluster475 | 1(4)\|3(1)\| | * | 100.0%[5/5] | 1.4 |
| Cluster474 | 1(4)\|3(1)\| | * | 100.0%[5/5] | 1.4 |
| Cluster472 | 1(4)\|3(1)\| | * | 100.0%[5/5] | 1.4 |
| Cluster471 | 1(4)\|3(1)\| | * | 100.0%[5/5] | 1.4 |
| Cluster466 | 1(4)\|3(1)\| | * | 100.0%[5/5] | 1.4 |
| Cluster465 | 1(4)\|3(1)\| | * | 100.0%[5/5] | 1.4 |
| Cluster464 | 1(4)\|3(1)\| | * | 100.0%[5/5] | 1.4 |
| Cluster463 | 1(4)\|3(1)\| | * | 100.0%[5/5] | 1.4 |
| Cluster43 | 1(3)\|2(2)\| | * | 100.0%[5/5] | 1.4 |
| Cluster425 | 1(3)\|2(2)\| | * | 100.0%[5/5] | 1.4 |
| Cluster422 | 0(1)\|1(1)\|2(3)\| | * | 80.0%[4/5] | 1.4 |
| Cluster412 | 0(1)\|1(1)\|2(3)\| | * | 80.0%[4/5] | 1.4 |
| Cluster383 | 1(4)\|3(1)\| | * | 100.0%[5/5] | 1.4 |
| Cluster382 | 1(4)\|3(1)\| | * | 100.0%[5/5] | 1.4 |
| Cluster381 | 1(4)\|3(1)\| | * | 100.0%[5/5] | 1.4 |
| Cluster380 | 1(4)\|3(1)\| | * | 100.0%[5/5] | 1.4 |
| Cluster375 | 1(4)\|3(1)\| | * | 100.0%[5/5] | 1.4 |
| Cluster308 | 1(3)\|2(2)\| | * | 100.0%[5/5] | 1.4 |
| Cluster297 | 0(1)\|1(3)\|4(1)\| | * | 80.0%[4/5] | 1.4 |
| Cluster29 | 0(1)\|1(2)\|2(1)\|3(1)\| | * | 80.0%[4/5] | 1.4 |
| Cluster259 | 1(4)\|3(1)\| | * | 100.0%[5/5] | 1.4 |
| Cluster247 | 1(4)\|3(1)\| | * | 100.0%[5/5] | 1.4 |

| | | | | |
|---|---|---|---|---|
| Cluster241 | 1(4)\|3(1)\| | * | 100.0%[5/5] | 1.4 |
| Cluster240 | 1(4)\|3(1)\| | * | 100.0%[5/5] | 1.4 |
| Cluster238 | 1(4)\|3(1)\| | * | 100.0%[5/5] | 1.4 |
| Cluster237 | 1(4)\|3(1)\| | * | 100.0%[5/5] | 1.4 |
| Cluster202 | 0(1)\|1(2)\|2(1)\|3(1)\| | * | 80.0%[4/5] | 1.4 |
| Cluster183 | 0(1)\|1(1)\|2(3)\| | * | 80.0%[4/5] | 1.4 |
| Cluster180 | 0(1)\|1(2)\|2(1)\|3(1)\| | * | 80.0%[4/5] | 1.4 |
| Cluster164 | 1(3)\|2(2)\| | * | 100.0%[5/5] | 1.4 |
| Cluster132 | 1(3)\|2(2)\| | * | 100.0%[5/5] | 1.4 |
| Cluster130 | 1(4)\|3(1)\| | * | 100.0%[5/5] | 1.4 |
| Cluster128 | 1(4)\|3(1)\| | * | 100.0%[5/5] | 1.4 |
| Cluster127 | 1(4)\|3(1)\| | * | 100.0%[5/5] | 1.4 |
| Cluster122 | 1(4)\|3(1)\| | * | 100.0%[5/5] | 1.4 |
| Cluster12 | 1(3)\|2(2)\| | * | 100.0%[5/5] | 1.4 |
| Cluster121 | 1(4)\|3(1)\| | * | 100.0%[5/5] | 1.4 |
| Cluster120 | 1(4)\|3(1)\| | * | 100.0%[5/5] | 1.4 |
| Cluster119 | 1(4)\|3(1)\| | * | 100.0%[5/5] | 1.4 |
| Cluster108 | 1(4)\|3(1)\| | * | 100.0%[5/5] | 1.4 |
| Cluster101 | 0(1)\|1(2)\|2(1)\|3(1)\| | * | 80.0%[4/5] | 1.4 |
| Cluster989 | 1(4)\|2(1)\| | * | 100.0%[5/5] | 1.2 |
| Cluster987 | 0(1)\|1(3)\|3(1)\| | * | 80.0%[4/5] | 1.2 |
| Cluster985 | 0(1)\|1(2)\|2(2)\| | * | 80.0%[4/5] | 1.2 |
| Cluster983 | 1(4)\|2(1)\| | * | 100.0%[5/5] | 1.2 |
| Cluster981 | 1(4)\|2(1)\| | * | 100.0%[5/5] | 1.2 |
| Cluster980 | 1(4)\|2(1)\| | * | 100.0%[5/5] | 1.2 |

| | | | | |
|---|---|---|---|---|
| Cluster976 | 0(1)\|1(3)\|3(1)\| | * | 80.0%[4/5] | 1.2 |
| Cluster975 | 0(1)\|1(3)\|3(1)\| | * | 80.0%[4/5] | 1.2 |
| Cluster972 | 0(1)\|1(2)\|2(2)\| | * | 80.0%[4/5] | 1.2 |
| Cluster963 | 0(1)\|1(2)\|2(2)\| | * | 80.0%[4/5] | 1.2 |
| Cluster947 | 0(1)\|1(2)\|2(2)\| | * | 80.0%[4/5] | 1.2 |
| Cluster923 | 1(4)\|2(1)\| | * | 100.0%[5/5] | 1.2 |
| Cluster922 | 1(4)\|2(1)\| | * | 100.0%[5/5] | 1.2 |
| Cluster918 | 0(1)\|1(3)\|3(1)\| | * | 80.0%[4/5] | 1.2 |
| Cluster913 | 1(4)\|2(1)\| | * | 100.0%[5/5] | 1.2 |
| Cluster911 | 1(4)\|2(1)\| | * | 100.0%[5/5] | 1.2 |
| Cluster908 | 0(1)\|1(3)\|3(1)\| | * | 80.0%[4/5] | 1.2 |
| Cluster897 | 1(4)\|2(1)\| | * | 100.0%[5/5] | 1.2 |
| Cluster885 | 1(4)\|2(1)\| | * | 100.0%[5/5] | 1.2 |
| Cluster884 | 1(4)\|2(1)\| | * | 100.0%[5/5] | 1.2 |
| Cluster883 | 1(4)\|2(1)\| | * | 100.0%[5/5] | 1.2 |
| Cluster882 | 1(4)\|2(1)\| | * | 100.0%[5/5] | 1.2 |
| Cluster848 | 1(4)\|2(1)\| | * | 100.0%[5/5] | 1.2 |
| Cluster845 | 0(1)\|1(3)\|3(1)\| | * | 80.0%[4/5] | 1.2 |
| Cluster837 | 1(4)\|2(1)\| | * | 100.0%[5/5] | 1.2 |
| Cluster826 | 0(1)\|1(3)\|3(1)\| | * | 80.0%[4/5] | 1.2 |
| Cluster808 | 1(4)\|2(1)\| | * | 100.0%[5/5] | 1.2 |
| Cluster787 | 0(1)\|1(2)\|2(2)\| | * | 80.0%[4/5] | 1.2 |
| Cluster775 | 1(4)\|2(1)\| | * | 100.0%[5/5] | 1.2 |
| Cluster759 | 1(4)\|2(1)\| | * | 100.0%[5/5] | 1.2 |
| Cluster755 | 0(1)\|1(3)\|3(1)\| | * | 80.0%[4/5] | 1.2 |

| | | | | |
|---|---|---|---|---|
| Cluster753 | 0(1)\|1(3)\|3(1)\| | * | 80.0%[4/5] | 1.2 |
| Cluster751 | 1(4)\|2(1)\| | * | 100.0%[5/5] | 1.2 |
| Cluster744 | 0(1)\|1(3)\|3(1)\| | * | 80.0%[4/5] | 1.2 |
| Cluster743 | 0(1)\|1(3)\|3(1)\| | * | 80.0%[4/5] | 1.2 |
| Cluster742 | 0(1)\|1(3)\|3(1)\| | * | 80.0%[4/5] | 1.2 |
| Cluster725 | 0(1)\|1(2)\|2(2)\| | * | 80.0%[4/5] | 1.2 |
| Cluster722 | 1(4)\|2(1)\| | * | 100.0%[5/5] | 1.2 |
| Cluster720 | 1(4)\|2(1)\| | * | 100.0%[5/5] | 1.2 |
| Cluster716 | 0(1)\|1(2)\|2(2)\| | * | 80.0%[4/5] | 1.2 |
| Cluster709 | 1(4)\|2(1)\| | * | 100.0%[5/5] | 1.2 |
| Cluster667 | 0(1)\|1(3)\|3(1)\| | * | 80.0%[4/5] | 1.2 |
| Cluster665 | 1(4)\|2(1)\| | * | 100.0%[5/5] | 1.2 |
| Cluster664 | 0(1)\|1(3)\|3(1)\| | * | 80.0%[4/5] | 1.2 |
| Cluster657 | 1(4)\|2(1)\| | * | 100.0%[5/5] | 1.2 |
| Cluster652 | 0(1)\|1(3)\|3(1)\| | * | 80.0%[4/5] | 1.2 |
| Cluster630 | 0(1)\|1(2)\|2(2)\| | * | 80.0%[4/5] | 1.2 |
| Cluster616 | 1(4)\|2(1)\| | * | 100.0%[5/5] | 1.2 |
| Cluster587 | 1(4)\|2(1)\| | * | 100.0%[5/5] | 1.2 |
| Cluster583 | 0(1)\|1(2)\|2(2)\| | * | 80.0%[4/5] | 1.2 |
| Cluster576 | 1(4)\|2(1)\| | * | 100.0%[5/5] | 1.2 |
| Cluster575 | 0(1)\|1(3)\|3(1)\| | * | 80.0%[4/5] | 1.2 |
| Cluster571 | 1(4)\|2(1)\| | * | 100.0%[5/5] | 1.2 |
| Cluster568 | 0(1)\|1(3)\|3(1)\| | * | 80.0%[4/5] | 1.2 |
| Cluster567 | 1(4)\|2(1)\| | * | 100.0%[5/5] | 1.2 |
| Cluster565 | 1(4)\|2(1)\| | * | 100.0%[5/5] | 1.2 |

| | | | | |
|---|---|---|---|---|
| Cluster564 | 1(4)\|2(1)\| | * | 100.0%[5/5] | 1.2 |
| Cluster561 | 0(1)\|1(3)\|3(1)\| | * | 80.0%[4/5] | 1.2 |
| Cluster55 | 1(4)\|2(1)\| | * | 100.0%[5/5] | 1.2 |
| Cluster531 | 0(1)\|1(2)\|2(2)\| | * | 80.0%[4/5] | 1.2 |
| Cluster528 | 1(4)\|2(1)\| | * | 100.0%[5/5] | 1.2 |
| Cluster526 | 1(4)\|2(1)\| | * | 100.0%[5/5] | 1.2 |
| Cluster516 | 1(4)\|2(1)\| | * | 100.0%[5/5] | 1.2 |
| Cluster514 | 1(4)\|2(1)\| | * | 100.0%[5/5] | 1.2 |
| Cluster512 | 1(4)\|2(1)\| | * | 100.0%[5/5] | 1.2 |
| Cluster5 | 0(1)\|1(3)\|3(1)\| | * | 80.0%[4/5] | 1.2 |
| Cluster49 | 1(4)\|2(1)\| | * | 100.0%[5/5] | 1.2 |
| Cluster482 | 0(1)\|1(2)\|2(2)\| | * | 80.0%[4/5] | 1.2 |
| Cluster478 | 1(4)\|2(1)\| | * | 100.0%[5/5] | 1.2 |
| Cluster477 | 1(4)\|2(1)\| | * | 100.0%[5/5] | 1.2 |
| Cluster469 | 1(4)\|2(1)\| | * | 100.0%[5/5] | 1.2 |
| Cluster467 | 1(4)\|2(1)\| | * | 100.0%[5/5] | 1.2 |
| Cluster434 | 0(1)\|1(2)\|2(2)\| | * | 80.0%[4/5] | 1.2 |
| Cluster418 | 0(1)\|1(2)\|2(2)\| | * | 80.0%[4/5] | 1.2 |
| Cluster397 | 1(4)\|2(1)\| | * | 100.0%[5/5] | 1.2 |
| Cluster396 | 1(4)\|2(1)\| | * | 100.0%[5/5] | 1.2 |
| Cluster387 | 0(1)\|1(3)\|3(1)\| | * | 80.0%[4/5] | 1.2 |
| Cluster379 | 1(4)\|2(1)\| | * | 100.0%[5/5] | 1.2 |
| Cluster373 | 0(1)\|1(2)\|2(2)\| | * | 80.0%[4/5] | 1.2 |
| Cluster372 | 1(4)\|2(1)\| | * | 100.0%[5/5] | 1.2 |
| Cluster371 | 1(4)\|2(1)\| | * | 100.0%[5/5] | 1.2 |

| | | | | |
|---|---|---|---|---|
| Cluster368 | 0(1)\|1(3)\|3(1)\| | * | 80.0%[4/5] | 1.2 |
| Cluster367 | 0(1)\|1(3)\|3(1)\| | * | 80.0%[4/5] | 1.2 |
| Cluster366 | 0(1)\|1(3)\|3(1)\| | * | 80.0%[4/5] | 1.2 |
| Cluster340 | 1(4)\|2(1)\| | * | 100.0%[5/5] | 1.2 |
| Cluster332 | 1(4)\|2(1)\| | * | 100.0%[5/5] | 1.2 |
| Cluster329 | 1(4)\|2(1)\| | * | 100.0%[5/5] | 1.2 |
| Cluster318 | 1(4)\|2(1)\| | * | 100.0%[5/5] | 1.2 |
| Cluster314 | 0(1)\|1(2)\|2(2)\| | * | 80.0%[4/5] | 1.2 |
| Cluster312 | 0(1)\|1(2)\|2(2)\| | * | 80.0%[4/5] | 1.2 |
| Cluster31 | 0(1)\|1(3)\|3(1)\| | * | 80.0%[4/5] | 1.2 |
| Cluster301 | 0(1)\|1(2)\|2(2)\| | * | 80.0%[4/5] | 1.2 |
| Cluster269 | 1(4)\|2(1)\| | * | 100.0%[5/5] | 1.2 |
| Cluster267 | 1(4)\|2(1)\| | * | 100.0%[5/5] | 1.2 |
| Cluster260 | 1(4)\|2(1)\| | * | 100.0%[5/5] | 1.2 |
| Cluster255 | 0(1)\|1(3)\|3(1)\| | * | 80.0%[4/5] | 1.2 |
| Cluster254 | 0(1)\|1(3)\|3(1)\| | * | 80.0%[4/5] | 1.2 |
| Cluster253 | 0(1)\|1(3)\|3(1)\| | * | 80.0%[4/5] | 1.2 |
| Cluster252 | 1(4)\|2(1)\| | * | 100.0%[5/5] | 1.2 |
| Cluster250 | 0(1)\|1(3)\|3(1)\| | * | 80.0%[4/5] | 1.2 |
| Cluster248 | 1(4)\|2(1)\| | * | 100.0%[5/5] | 1.2 |
| Cluster244 | 0(1)\|1(3)\|3(1)\| | * | 80.0%[4/5] | 1.2 |
| Cluster242 | 0(1)\|1(3)\|3(1)\| | * | 80.0%[4/5] | 1.2 |
| Cluster198 | 1(4)\|2(1)\| | * | 100.0%[5/5] | 1.2 |
| Cluster197 | 1(4)\|2(1)\| | * | 100.0%[5/5] | 1.2 |
| Cluster182 | 1(4)\|2(1)\| | * | 100.0%[5/5] | 1.2 |

| | | | | |
|---|---|---|---|---|
| Cluster161 | 1(4)\|2(1)\| | * | 100.0%[5/5] | 1.2 |
| Cluster160 | 0(1)\|1(2)\|2(2)\| | * | 80.0%[4/5] | 1.2 |
| Cluster152 | 0(1)\|1(2)\|2(2)\| | * | 80.0%[4/5] | 1.2 |
| Cluster147 | 1(4)\|2(1)\| | * | 100.0%[5/5] | 1.2 |
| Cluster146 | 0(1)\|1(2)\|2(2)\| | * | 80.0%[4/5] | 1.2 |
| Cluster129 | 0(1)\|1(3)\|3(1)\| | * | 80.0%[4/5] | 1.2 |
| Cluster126 | 0(1)\|1(3)\|3(1)\| | * | 80.0%[4/5] | 1.2 |
| Cluster115 | 1(4)\|2(1)\| | * | 100.0%[5/5] | 1.2 |
| Cluster113 | 1(4)\|2(1)\| | * | 100.0%[5/5] | 1.2 |
| Cluster111 | 0(1)\|1(3)\|3(1)\| | * | 80.0%[4/5] | 1.2 |
| Cluster110 | 1(4)\|2(1)\| | * | 100.0%[5/5] | 1.2 |
| Cluster107 | 1(4)\|2(1)\| | * | 100.0%[5/5] | 1.2 |
| Cluster1002 | 1(4)\|2(1)\| | * | 100.0%[5/5] | 1.2 |
| Cluster982 | 0(1)\|1(3)\|2(1)\| | * | 80.0%[4/5] | 1 |
| Cluster979 | 0(1)\|1(3)\|2(1)\| | * | 80.0%[4/5] | 1 |
| Cluster955 | 0(1)\|1(3)\|2(1)\| | * | 80.0%[4/5] | 1 |
| Cluster952 | 1(5)\| | * | 100.0%[5/5] | 1 |
| Cluster948 | 0(1)\|1(3)\|2(1)\| | * | 80.0%[4/5] | 1 |
| Cluster937 | 1(5)\| | * | 100.0%[5/5] | 1 |
| Cluster932 | 0(1)\|1(3)\|2(1)\| | * | 80.0%[4/5] | 1 |
| Cluster929 | 0(1)\|1(3)\|2(1)\| | * | 80.0%[4/5] | 1 |
| Cluster912 | 0(1)\|1(3)\|2(1)\| | * | 80.0%[4/5] | 1 |
| Cluster900 | 0(1)\|1(3)\|2(1)\| | * | 80.0%[4/5] | 1 |
| Cluster898 | 0(1)\|1(3)\|2(1)\| | * | 80.0%[4/5] | 1 |
| Cluster876 | 1(5)\| | * | 100.0%[5/5] | 1 |

| | | | | |
|---|---|---|---|---|
| Cluster864 | 1(5)\| | * | 100.0%[5/5] | 1 |
| Cluster853 | 1(5)\| | * | 100.0%[5/5] | 1 |
| Cluster840 | 0(1)\|1(3)\|2(1)\| | * | 80.0%[4/5] | 1 |
| Cluster838 | 0(1)\|1(3)\|2(1)\| | * | 80.0%[4/5] | 1 |
| Cluster809 | 0(1)\|1(3)\|2(1)\| | * | 80.0%[4/5] | 1 |
| Cluster805 | 0(1)\|1(3)\|2(1)\| | * | 80.0%[4/5] | 1 |
| Cluster798 | 0(1)\|1(3)\|2(1)\| | * | 80.0%[4/5] | 1 |
| Cluster768 | 0(1)\|1(3)\|2(1)\| | * | 80.0%[4/5] | 1 |
| Cluster767 | 1(5)\| | * | 100.0%[5/5] | 1 |
| Cluster746 | 0(1)\|1(3)\|2(1)\| | * | 80.0%[4/5] | 1 |
| Cluster702 | 0(1)\|1(3)\|2(1)\| | * | 80.0%[4/5] | 1 |
| Cluster697 | 1(5)\| | * | 100.0%[5/5] | 1 |
| Cluster695 | 1(5)\| | * | 100.0%[5/5] | 1 |
| Cluster687 | 1(5)\| | * | 100.0%[5/5] | 1 |
| Cluster683 | 1(5)\| | * | 100.0%[5/5] | 1 |
| Cluster682 | 1(5)\| | * | 100.0%[5/5] | 1 |
| Cluster681 | 1(5)\| | * | 100.0%[5/5] | 1 |
| Cluster678 | 1(5)\| | * | 100.0%[5/5] | 1 |
| Cluster670 | 0(1)\|1(3)\|2(1)\| | * | 80.0%[4/5] | 1 |
| Cluster656 | 0(1)\|1(3)\|2(1)\| | * | 80.0%[4/5] | 1 |
| Cluster638 | 1(5)\| | * | 100.0%[5/5] | 1 |
| Cluster637 | 1(5)\| | * | 100.0%[5/5] | 1 |
| Cluster635 | 0(1)\|1(3)\|2(1)\| | * | 80.0%[4/5] | 1 |
| Cluster634 | 0(1)\|1(3)\|2(1)\| | * | 80.0%[4/5] | 1 |
| Cluster631 | 0(1)\|1(3)\|2(1)\| | * | 80.0%[4/5] | 1 |

| | | | | |
|---|---|---|---|---|
| Cluster614 | 1(5)\| | * | 100.0%[5/5] | 1 |
| Cluster612 | 0(1)\|1(3)\|2(1)\| | * | 80.0%[4/5] | 1 |
| Cluster608 | 0(1)\|1(3)\|2(1)\| | * | 80.0%[4/5] | 1 |
| Cluster604 | 0(1)\|1(3)\|2(1)\| | * | 80.0%[4/5] | 1 |
| Cluster589 | 1(5)\| | * | 100.0%[5/5] | 1 |
| Cluster579 | 0(1)\|1(3)\|2(1)\| | * | 80.0%[4/5] | 1 |
| Cluster572 | 0(1)\|1(3)\|2(1)\| | * | 80.0%[4/5] | 1 |
| Cluster541 | 0(1)\|1(3)\|2(1)\| | * | 80.0%[4/5] | 1 |
| Cluster515 | 0(1)\|1(3)\|2(1)\| | * | 80.0%[4/5] | 1 |
| Cluster508 | 1(5)\| | * | 100.0%[5/5] | 1 |
| Cluster488 | 1(5)\| | * | 100.0%[5/5] | 1 |
| Cluster486 | 1(5)\| | * | 100.0%[5/5] | 1 |
| Cluster485 | 1(5)\| | * | 100.0%[5/5] | 1 |
| Cluster473 | 0(1)\|1(3)\|2(1)\| | * | 80.0%[4/5] | 1 |
| Cluster468 | 0(1)\|1(3)\|2(1)\| | * | 80.0%[4/5] | 1 |
| Cluster449 | 1(5)\| | * | 100.0%[5/5] | 1 |
| Cluster444 | 0(1)\|1(3)\|2(1)\| | * | 80.0%[4/5] | 1 |
| Cluster443 | 0(1)\|1(3)\|2(1)\| | * | 80.0%[4/5] | 1 |
| Cluster430 | 0(1)\|1(3)\|2(1)\| | * | 80.0%[4/5] | 1 |
| Cluster429 | 0(1)\|1(3)\|2(1)\| | * | 80.0%[4/5] | 1 |
| Cluster415 | 0(1)\|1(3)\|2(1)\| | * | 80.0%[4/5] | 1 |
| Cluster405 | 1(5)\| | * | 100.0%[5/5] | 1 |
| Cluster404 | 1(5)\| | * | 100.0%[5/5] | 1 |
| Cluster394 | 0(1)\|1(3)\|2(1)\| | * | 80.0%[4/5] | 1 |
| Cluster385 | 0(1)\|1(3)\|2(1)\| | * | 80.0%[4/5] | 1 |

| | | | | |
|---|---|---|---|---|
| Cluster316 | 0(1)\|1(3)\|2(1)\| | * | 80.0%[4/5] | 1 |
| Cluster309 | 1(5)\| | * | 100.0%[5/5] | 1 |
| Cluster283 | 1(5)\| | * | 100.0%[5/5] | 1 |
| Cluster278 | 1(5)\| | * | 100.0%[5/5] | 1 |
| Cluster273 | 0(1)\|1(3)\|2(1)\| | * | 80.0%[4/5] | 1 |
| Cluster271 | 0(1)\|1(3)\|2(1)\| | * | 80.0%[4/5] | 1 |
| Cluster27 | 0(1)\|1(3)\|2(1)\| | * | 80.0%[4/5] | 1 |
| Cluster266 | 1(5)\| | * | 100.0%[5/5] | 1 |
| Cluster264 | 1(5)\| | * | 100.0%[5/5] | 1 |
| Cluster263 | 0(1)\|1(3)\|2(1)\| | * | 80.0%[4/5] | 1 |
| Cluster258 | 0(1)\|1(3)\|2(1)\| | * | 80.0%[4/5] | 1 |
| Cluster249 | 0(1)\|1(3)\|2(1)\| | * | 80.0%[4/5] | 1 |
| Cluster245 | 0(1)\|1(3)\|2(1)\| | * | 80.0%[4/5] | 1 |
| Cluster176 | 0(1)\|1(3)\|2(1)\| | * | 80.0%[4/5] | 1 |
| Cluster154 | 0(1)\|1(3)\|2(1)\| | * | 80.0%[4/5] | 1 |
| Cluster123 | 0(1)\|1(3)\|2(1)\| | * | 80.0%[4/5] | 1 |
| Cluster118 | 0(1)\|1(3)\|2(1)\| | * | 80.0%[4/5] | 1 |
| Cluster1003 | 1(5)\| | * | 100.0%[5/5] | 1 |
| Cluster1000 | 1(5)\| | * | 100.0%[5/5] | 1 |
| Cluster944 | 0(1)\|1(4)\| | * | 80.0%[4/5] | 0.8 |
| Cluster943 | 0(1)\|1(4)\| | * | 80.0%[4/5] | 0.8 |
| Cluster933 | 0(1)\|1(4)\| | * | 80.0%[4/5] | 0.8 |
| Cluster930 | 0(1)\|1(4)\| | * | 80.0%[4/5] | 0.8 |
| Cluster877 | 0(1)\|1(4)\| | * | 80.0%[4/5] | 0.8 |
| Cluster863 | 0(1)\|1(4)\| | * | 80.0%[4/5] | 0.8 |

| | | | | |
|---|---|---|---|---|
| Cluster781 | 0(1)\|1(4)\| | * | 80.0%[4/5] | 0.8 |
| Cluster770 | 0(1)\|1(4)\| | * | 80.0%[4/5] | 0.8 |
| Cluster713 | 0(1)\|1(4)\| | * | 80.0%[4/5] | 0.8 |
| Cluster712 | 0(1)\|1(4)\| | * | 80.0%[4/5] | 0.8 |
| Cluster680 | 0(1)\|1(4)\| | * | 80.0%[4/5] | 0.8 |
| Cluster677 | 0(1)\|1(4)\| | * | 80.0%[4/5] | 0.8 |
| Cluster632 | 0(1)\|1(4)\| | * | 80.0%[4/5] | 0.8 |
| Cluster615 | 0(1)\|1(4)\| | * | 80.0%[4/5] | 0.8 |
| Cluster599 | 0(1)\|1(4)\| | * | 80.0%[4/5] | 0.8 |
| Cluster591 | 0(1)\|1(4)\| | * | 80.0%[4/5] | 0.8 |
| Cluster537 | 0(1)\|1(4)\| | * | 80.0%[4/5] | 0.8 |
| Cluster446 | 0(1)\|1(4)\| | * | 80.0%[4/5] | 0.8 |
| Cluster426 | 0(1)\|1(4)\| | * | 80.0%[4/5] | 0.8 |
| Cluster417 | 0(1)\|1(4)\| | * | 80.0%[4/5] | 0.8 |
| Cluster406 | 0(1)\|1(4)\| | * | 80.0%[4/5] | 0.8 |
| Cluster392 | 0(1)\|1(4)\| | * | 80.0%[4/5] | 0.8 |
| Cluster336 | 0(1)\|1(4)\| | * | 80.0%[4/5] | 0.8 |
| Cluster279 | 0(1)\|1(4)\| | * | 80.0%[4/5] | 0.8 |
| Cluster277 | 0(1)\|1(4)\| | * | 80.0%[4/5] | 0.8 |
| Cluster265 | 0(1)\|1(4)\| | * | 80.0%[4/5] | 0.8 |
| Cluster207 | 0(1)\|1(4)\| | * | 80.0%[4/5] | 0.8 |
| Cluster177 | 0(1)\|1(4)\| | * | 80.0%[4/5] | 0.8 |
